# A first-in-class multimodal organomercury compound demonstrates preferential blast reduction with improved hematopoietic and immune health in ALL

**DOI:** 10.64898/2026.07.18.739301

**Authors:** Sougata Mondal, Oyendrila Ghosh, Basab Bagchi, Pratima Jana, Bidisha Maiti, Kalyan Kusum Mukherjee, Supratim Ghosh

## Abstract

Acute leukemia treatment requires effective blast reduction while improving hematopoietic and immune health. As reported previously, we developed a structurally distinct organomercury derivative of curcumin, α-Mercurin, having improved aqueous solubility and intravenous applicability. Here we comprehensively evaluated therapeutic efficacy, pharmacological response, systemic toxicity, blood Hg pharmacokinetics (PK), Hg biodistribution and elimination profile for α-Mercurin in a N-nitroso-N-ethylurea (ENU)-induced autochthonous Acute lymphoblastic leukemia (ALL) rat model, comparing with clinical standard cytarabine. Results showed recovery in body-weight, along with longer time-to-humane-endpoint from 13 to 39 days. Treatment successfully reduced immature blast count, while promoting erythroid, myeloid, and megakaryocytic compartments and improving immune mediators across peripheral blood, bone marrow, thymus, spleen, and lymph nodes. Histopathology showed reduced leukemic infiltration with preservation of major organ architecture, while cytarabine produced evident myelosuppression, neurotoxicity, and cardiotoxicity. PK study showed blood Hg reached T_max_ at 15 min, followed by urinary and fecal elimination with residual retention in kidney tissue and no detectable Hg in brain. Collectively, α-Mercurin demonstrated a promising multimodal therapeutic index, combining cytoreductive activity with improvement in hematopoietic and immune health, along with favorable safety profile for future ALL therapy.

**GRAPHICAL ABSTRACT:** 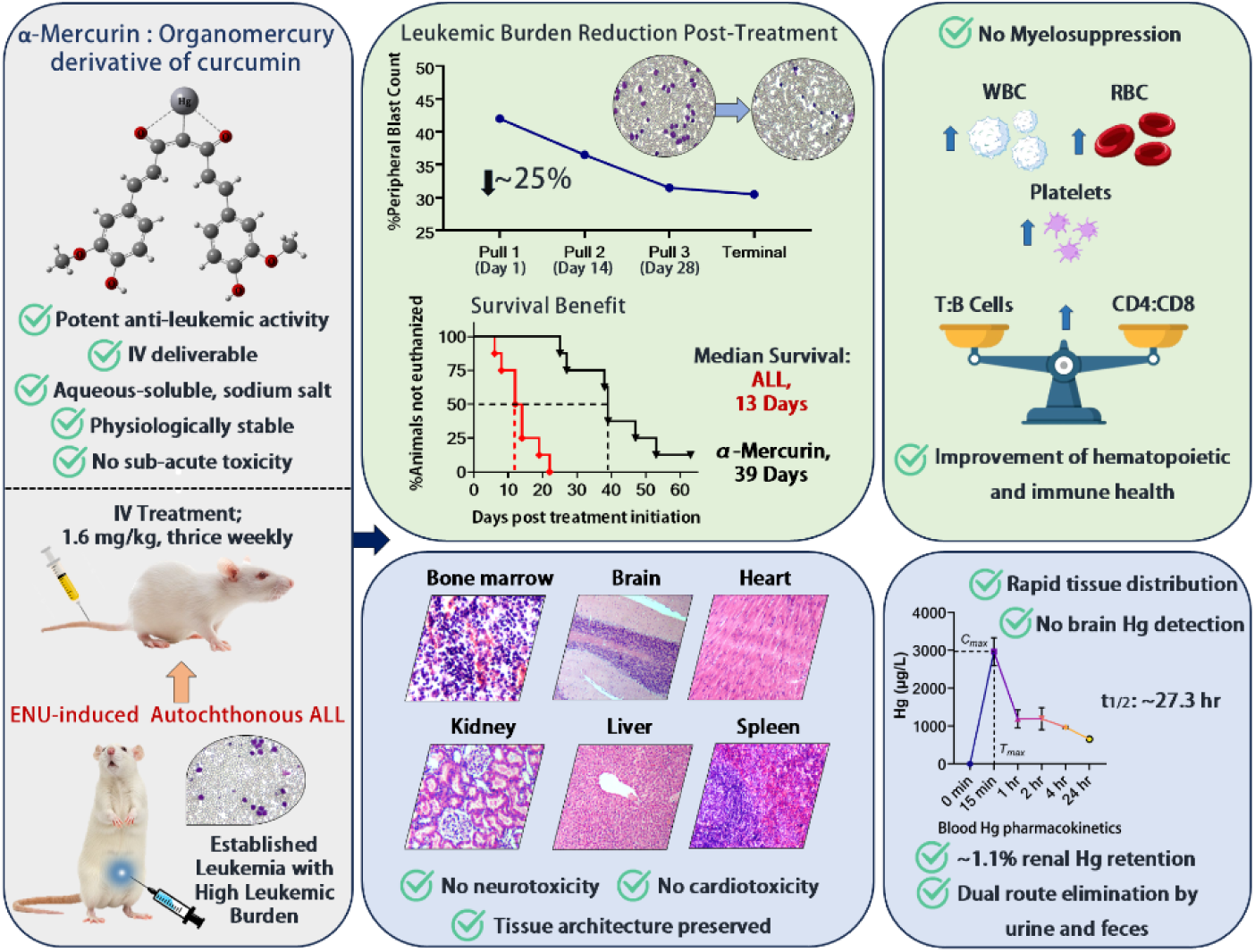

## 1. INTRODUCTION

Acute leukemia remains a major therapeutic challenge(1,2), especially in children and elderly people(3–8). Besides direct leukemic burden, acute leukemia profoundly disrupts hematopoietic activity and immune balance through suppression of normal bone marrow function(9–12). This disease progression can result in anemia, thrombocytopenia, abnormal leukocyte counts, and immune dysfunction(13–15). Current treatment mainly relies on combination chemotherapy, supported by targeted inhibitors and immunotherapies in selected subtypes(16–21). However, effective targeted therapeutics are not available for many biological subtypes, thus dependent on cytotoxic chemotherapeutic modality as an important therapeutic backbone for a substantial proportion of patients(22). Although these agents can achieve significant cytoreduction, their therapeutic activity is accompanied by treatment-associated toxicities, particularly myelosuppression, which can affect erythroid, myeloid, and megakaryocytic compartments and compromise treatment outcome(23–25). Neurological, hepatic, cardiac, and other systemic toxicities have also been reported depending on treatment intensity and exposure(26,27).

Metal-containing therapeutics provide a promising perspective in this context(28,29), as several metal and metalloid compounds have demonstrated therapeutic activity in hematological malignancies(30,31), with arsenic trioxide being an established example for acute promyelocytic leukemia(32). However, mercury (Hg)-containing compounds have historically been associated with systemic toxicity(33,34) and consequently received very limited attention as therapeutic candidate. We therefore hypothesized that this toxicity-centered perception could be addressed with novel formulation, suitable route of administration and optimal dosing to achieve therapeutic benefits.

Curcumin, a naturally occurring polyphenolic molecule, has attracted considerable interest because of its anti-proliferative, anti-inflammatory, and immunomodulatory properties(35–37). Its pharmacological activities have been associated with modulation of multiple cellular pathways, including NF-κB and STAT3, inflammatory mediators, and ROS-associated mitochondrial apoptosis(38–42). However, the clinical application of curcumin remains limited due to poor aqueous solubility and chemical instability(43,44). Several approaches including nanoformulations have been explored to improve its bioavailability(45,46). However, formulation-related limitations such as increased particle size, variable biodistribution, and complex delivery requirements remained as important constrains for systemic administration(47,48). Another approach has been the conjugation of metal or metalloid ions to β-diketo oxygens of curcumin, however, resultant derivatives either changed the biochemical properties of curcumin or require amphiphilic solvents such as DMSO for solubility(49–52). The β-diketone moiety also undergoes keto-enol tautomerism, which can make conventional O-chelate complexes susceptible in ionic solvent(43,53,54). Additionally, the α-carbon of the conjugated curcumin framework remains vulnerable to hydrolytic and oxidative degradation in aqueous biological conditions(55,56). To address these limitations, we previously developed α-Mercurin as a structurally distinct organometallic derivative of curcumin by directly binding Hg to the α-carbon via formation of a stable C_α_–Hg bond(57). This strategy modifies the carbon framework of curcumin, while maintaining its overall conjugated structure. α-Mercurin is soluble in the aqueous medium as sodium salt and suitable for intravenous (IV) administration. In our previous study, we demonstrated that α-Mercurin preferentially eliminated leukemic cells through ROS-associated mitochondrial dysfunction and apoptosis, while showing no significant cytotoxicity towards peripheral blood cells from healthy individuals in vitro, ex vivo, and in vivo(57). These findings provided an initial pharmacodynamic rationale for further evaluation of α-Mercurin under physiological context.

The present study is designed to comprehensively evaluate therapeutic potential of α-Mercurin in a N-nitroso-N-ethylurea (ENU)-induced autochthonous Acute Lymphoblastic Leukemia (ALL) rat model having intact immune and hematopoietic physiology(58–61). Therapeutic benefit of α-Mercurin is investigated in terms of disease progression, body-weight, survival duration, peripheral hematological parameters, bone marrow cellularity, and immune health across peripheral blood (PB) and lymphoid compartments. We further examined tissue-level compatibility through histopathology, and serum biochemical profiling. Hg biodistribution is studied using cold vapor atomic absorption spectrometry (CV-AAS), in combination with blood Hg pharmacokinetics (PK) and urinary as well as fecal Hg excretion. Oxidative stress to liver and kidney tissues following α-Mercurin treatment is assessed via GSH:GSSG ratio. Across the study, cytarabine is used as the clinically established therapeutic standard to understand the overall therapeutic potential of α-Mercurin. To our knowledge, no previous work has reported any intravenously deliverable formulation of Hg with comprehensive in vivo assessment. This study therefore provides a pharmacological foundation with pharmacokinetics/pharmacodynamics (PK/PD) characterization and comprehensive translational assessment of an organomercury compound, α-Mercurin for therapeutic application against acute leukemia.

## 2. MATERIALS AND METHODS

### 2.1. MATERIALS

Curcumin (Sigma-Aldrich, Cat No. C7727); Mercury(II) chloride (Sigma-Aldrich, Cat No. 215465); Ammonium hydroxide (Supelco, Cat No. 1935000521); N-Nitroso-N-ethylurea (Sigma-Aldrich, Cat No. N8509-5G); Cytarabine, ≥98% (HPLC) (Sigma-Aldrich, Cat No. 251010); Mercury Standard for AAS (Supelco, Cat No. 16482); Tin(II) chloride dihydrate, ACS, ≥98% (Sigma-Aldrich, Cat No. 31669); Hydrochloric acid about 37%, ACS (Supelco, Cat No. 1.93001.0522); Nitric acid about 69% (Supelco, Cat No. 1.93406.0521); Sodium Chloride ACS, 99.9% (SRL, Cat No. 41721); Potassium Chloride ACS, 99.5% (SRL, Cat No. 38630); Sodium Phosphate Dibasic Dihydrate extrapure AR, 99.5% (SRL, Cat No. 87258); Potassium phosphate monobasic (Supelco, Cat No. 1.4873.0250); Citric acid ACS reagent grade, ≥99.5%, crystals (Sigma-Aldrich, Cat No. 251275-500G); K2 EDTA 2ml vacutainer blood collection tube (BD, Cat No. 367841); K2 EDTA 0.5ml microtainer blood collection tube (BD, Cat No. 365974); PerCP/Cyanine5.5 anti-rat CD3 Antibody (Biolegend, Cat No. 201418); CD19 Antibody (F-3) Alexa Fluor 680 (Santa Cruz, Cat No. sc-373897 AF680); NCAM/CD56 Antibody (123C3) Alexa Fluor 647 (Santa Cruz, Cat No. sc-7326 AF647); FITC anti-rat CD4 Antibody (Biolegend, Cat No. 201505); PE anti-rat CD8a Antibody (Biolegend, Cat No. 200608); CD45 Monoclonal Antibody (OX1), FITC (Invitrogen, Cat No. 11-0461-82); CD34 (Hematopoietic Stem Cell and Endothelial Marker) Monoclonal Antibody (ICO-115), PE (Invitrogen, Cat No. 947-MSM1-PE-100T); Histosec 60 pastilles (without DMSO) (Sigma-Aldrich, Cat No. 1.01676.2504); Hematoxylin solution (Harris) for microscopy (SRL, Cat No. 40362); Eosin Yellow (water soluble), ACS (SRL, Cat No. 29391); DAPI (Invitrogen, Cat No. D1306); DPX Mountant for histology (SRL, Cat No. 88147); Xylene extrapure AR, ACS (SRL, Cat No. 89159); EtOH Absolute (Adventol, Cat No. 90056); Micro 75mmx25mm Super Deluxe 1.10mm Slide (Blue Star); tert-Butanol (Supelco, Cat No. 8.22264.0521); Glycerol (Glycerine) (SRL, Cat No. 62417); Formaldehyde solution min 37% (Emplura, Cat No. 1.94989.0521); Cholesterol Clinical Chemistry (Vanguard, Cat No. 300038); Albumin Clinical Chemistry (Vanguard, Cat No. 300046); Urea UV Clinical Chemistry (Vanguard, Cat No. 300042); Alkaline Phosphatase Clinical Chemistry (Vanguard, Cat No. 300045); SGOT Clinical Chemistry (Vanguard, Cat No. 300043); SGPT Clinical Chemistry (Vanguard, Cat No. 300044); Bilirubin T&D Clinical Chemistry (Vanguard, Cat No. 300021A); Acetic Acid Glacial ACS, 99.9% (SRL, Cat No. 85801), Glutathione GSH/GSSG Assay Kit (Sigma-Aldrich, Cat No. MAK440-1KT), meta-Phosphoric acid ACS reagent, chips, 33.5-36.5% (Sigma-Aldrich, Cat No. 239275-5G).

### 2.2. METHODS

#### 2.2.1. Animals handling and monitoring

Male Wistar rats (age: 5 - 6 weeks old, body weight: 125 ± 20 g) were obtained from the Centre for Laboratory Animal Research and Training (CLART), West Bengal Livestock Development Corporation Ltd., India, and housed in the animal house facility of the Chittaranjan National Cancer Institute (CNCI). All procedures were approved by the Institutional Animal Ethics Committee of the CNCI (IAEC-1774/SG-8(Rev)/2024/09) and conducted in accordance with CCSEA and ARRIVE 2.0 guidelines. Rats were acquired and maintained under specific pathogen-free conditions in polypropylene cages (410 × 282 × 153 mm) with stainless steel wire tops, housing two animals per cage.

Environmental conditions were controlled at 24 ± 2°C with 40 - 70% relative humidity under a 12 h light/12 h dark cycle. Animals were acclimatized for one week prior to initiation of experiments. Standard rodent chow and water were provided as necessary. Bedding and cages were changed regularly. Body weights were recorded at procurement (day 0) and weekly throughout the study. All animals were monitored daily for general health, behavioral changes and signs of leukemia progression or treatment-associated toxicity.

#### 2.2.2. Preparation and administration of α-Mercurin and cytarabine

##### 2.2.2.1. α-Mercurin preparation and dosing

α-Mercurin was synthesized as previously reported(57). For in vivo administration, 1.3 mg of dry α-Mercurin was dissolved in 1.6 mL of 5 mM aqueous NaOH, followed by centrifugation at 8,650×g for 7 min (REMI NEYA 16R centrifuge). The supernatant was filtered using a 0.2 µm PES syringe filter prior to intravenous (IV) administration. Fresh preparations were made before each injection. α-Mercurin was administered at 1.6 mg/kg body weight, three times per week with a one-day interval between doses. The treatment regimen was selected based on our previous sub-acute toxicity study(57). Treatment duration extended up to 8 weeks or until humane endpoint criteria were met.

##### 2.2.2.2. Cytarabine preparation and dosing

Cytarabine (cytosine arabinoside; Ara-C) was dissolved in sterile saline and filtered prior to IV administration. It was administered at 5 mg/kg body weight, thrice weekly, consistent with low-dose maintenance regimens in acute lymphoblastic leukemia (ALL) to achieve a cumulative dose of ∼100–150 mg/kg body weight(62). Treatment duration extended up to 8 weeks or until humane endpoint criteria were met.

#### 2.2.3. ENU preparation and induction of leukemia

ENU (N-Nitroso-N-ethylurea) was stored at −20°C in the dark and freshly prepared prior to each administration. The required quantity was aliquoted under an argon-filled glove bag, dissolved in 95% ethanol (5% water), and subsequently diluted in 0.15 M phosphate-citrate buffer (pH ∼5.0)(63). Solutions were protected from light throughout preparation and administration. Leukemia was induced by intraperitoneal (IP) injection of ENU (80 mg/kg body weight) once weekly for 15 weeks using 24-gauge needle. Animals were monitored weekly by peripheral blood smear analysis for the presence of immature blast cells. Additional assessments included palpation of spleen and liver for any enlargement, along with measurement of total leukocyte count, differential counts, erythrocytes, platelets and hemoglobin (Hb) levels. Animals exhibiting white blood Cell (WBC) counts ≥25×10^3^/mm^3^ with ≥35% circulating blasts were considered to have advanced ALL and were enrolled for treatment.

#### 2.2.4. Experimental design and animal allocation

A total of 54 rats were used, of which 46 received ENU. Following leukemia establishment, 24 animals with advanced ALL were randomized into three groups (n = 8 each): ALL control (ALL Con), ALL treatment with cytarabine (ALL+Cyt) and ALL treatment with α-Mercurin (ALL+α-Mer). Additional groups included healthy control (Healthy Con; n = 4) and α-Mercurin control (α-Mer Con; n = 4). Longitudinal sampling was performed at defined timepoints: hereafter referred to as Pull 1 (day 1, just before treatment initiation), Pull 2 (day 14, post-treatment), Pull 3 (day 28, post-treatment) and terminal analysis (at euthanasia). Time dependent variation in animal numbers across different groups occurred due to humane-endpoint-based euthanasia during the study. Where applicable, investigators were blinded during group allocation, data acquisition and downstream analysis.

##### 2.2.4.1. Number of animals (n) across timepoints

Variation in animal numbers (n) across timepoints occurred due to humane endpoint-based euthanasia during disease progression.

| Timepoint | Healthy Control | α-Mercurin Control | Leukemia Control | Leukemia+ Cytarabine | Leukemia+α-Mercurin |
| --- | --- | --- | --- | --- | --- |
| Pull 1 (day 1, just before treatment) | 4 | 4 | 8 | 8 | 8 |
| Pull 2 (day 14, post-treatment) | 4 | 4 | 4 | 8 | 8 |
| Pull 3 (day 28, post-treatment) | 4 | 4 | No survival | 4 | 6 |
| Terminal (at euthanasia) | 4 | 4 | 8 | 8 | 8 |

#### 2.2.5. Baseline body weight determination

Baseline body weight was defined as the body weight measured on the day of group randomization for treatment initiation. This body weight was used as the reference point for calculating disease-associated body-weight loss during the study. Percentage body-weight loss was calculated using the formula:

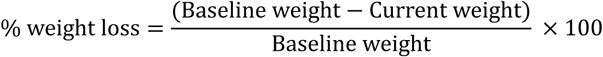

#### 2.2.6. Humane endpoint criteria and monitoring

Animals were humanely euthanized upon meeting any one or combination of the following predefined criteria(64) as indication of severe leukemia-associated clinical deterioration:

a. Sustained loss of 10 - 15% of body-weight from the baseline, accompanied by additional clinical signs of morbidity.
b. Severe lethargy or impaired mobility, characterized by significant reduction of spontaneous movement or inability to maintain normal posture.
c. Inability to access food or water, evidenced by failure to reach feeding or drinking sources despite availability.
d. Pale discoloration of extremities such as feet, tail, ears, nose or eyes; often accompanied by reduced peripheral temperature.

#### 2.2.7. Monitoring and euthanasia procedure

Clinical assessments, including body weight measurement and behavioural observation were conducted daily. Upon exhibiting rapid deterioration or multiple concurrent signs, animals were euthanized immediately using an institutionally approved method (Compendium of CCSEA(65)), and tissues were collected and fixed in 10% formalin for downstream analyses. Time-to-humane-endpoint analysis was performed using Kaplan-Meier methodology.

#### 2.2.8. Tissue collection and downstream analyses

At euthanasia, blood, major organs (brain, thymus, heart, lung, liver, kidney, spleen, lymph node, femur bone) and bone marrow were collected for histological, immunological and biochemical analysis. Bone marrow aspirates and blood samples were used for smear preparation and flow cytometry. Bone marrow aspirates were also used for immunofluorescence studies. Serum was isolated for clinical biochemistry assessment.

#### 2.2.9. Peripheral blood sampling

Peripheral blood smearing was done by dropping ∼10 µl blood on clean glass slide and spreading with another slide. All the smears were air dried and further stained with Leishman’s stain. CBC along with Hb concentration and flow cytometric analysis for immune response were performed and represented as Pull 1 (day 1, just before treatment), Pull 2 (day 14, post-treatment) and Pull 3 (day 28, post-treatment) as well as terminal analysis upon euthanasia. Clinical biochemistry assessment was done using the serum isolated from terminally collected ∼2 ml blood samples in polypropylene tubes.

#### 2.2.10. Investigation of hematological parameters

Hematological analyses from peripheral blood smear were performed to evaluate leukemia progression and treatment response. Total WBC along with differential leukocyte counts were performed to quantify neutrophils, eosinophils, basophils, lymphocytes and monocytes, along with the proportion of circulating blasts. Circulating blasts were identified based on nuclear-to-cytoplasmic (N/C) ratio, chromatin pattern, nucleolar prominence and irregular cell morphology. In addition, erythrocyte-related parameters, including red blood cell (RBC) count, Hb concentration and platelet count, were measured to assess anemia and thrombocytopenia-associated disease severity. All hematological parameters were expressed as median with IQR (25 - 75 percentile) in box plots (showing all points); whiskers represent minimum-maximum values, and compared between experimental groups to determine treatment-associated normalization of leukemia-induced hematological abnormalities. All Leishman’s-stained slides were visualised under Zeiss Primovert inverted microscope equipped with Tucsen GT 5.0 5MP USB2.0 CMOS Camera, using 10X, 20X, 40X objective lens.

#### 2.2.11. Clinical biochemistry

Clinical biochemistry was analyzed using serum samples of terminally collected blood samples. Following retroorbital bleeding, blood samples were allowed to clot at room temperature for 30 min and subsequently centrifuged at 2,000×g for 10 min at 4⁰C. The serum layer was collected carefully and stored at −80⁰C until further analysis. Serum biochemical parameters including total cholesterol, albumin, urea, blood urea nitrogen (BUN), alkaline phosphatase (ALP), aspartate aminotransferase (AST/SGOT), alanine aminotransferase (ALT/SGPT) and total bilirubin were quantified using commercially available diagnostic kits, according to the manufacturer’s instructions.

#### 2.2.12. Flow cytometry analysis

For all the flow cytometric acquisition, antibodies were used at concentrations instructed by manufacturers. For peripheral blood, following blood sampling in K2 EDTA blood collection tubes, 1X RBC lysis solution was added to whole blood at the ratio of whole blood:RBC lysis solution, 1:10. Following incubation at room temperature for 10 min in the dark, cells were washed twice using sterile PBS, centrifuging at 600×g for 7 min. Further, cells (∼1X10^6^) were incubated with predetermined antibodies for 45 min at room temperature (RT) in the dark and again washed two times with sterile PBS before flow cytometry. Cells were further resuspended in sterile PBS and analyzed using LSRFortessa (BD Biosciences). For isolated single cells from mentioned organs, excluding RBC lysis all the steps were followed as described above. Single stain controls were used to properly compensate for spill over between different fluorochromes. Final data analysis was performed with the FlowJo software version 10.8.1.

Whole blood after excluding RBCs, was stained with mAbs specific for CD3, CD19, CD56, CD4 and CD8 followed by flow cytometric analysis. After gating on lymphocyte compartment and excluding doublets, CD3^+^ T cells, CD19^+^ B cells, CD56^+^ NK and CD3^-^ CD19^-^CD56^-^ (triple-negative; TN^-^) cells were presented as percent of total lymphocytes. CD4^+^, CD8^+^ and CD4^+^CD8^+^ T cell subsets were further studied and presented as percent of total CD3^+^ T cells. To understand immune homeostasis as well as dysregulation in contrast to malignant expansion of leukemic blasts, CD4^+^:CD8^+^ and CD3^+^ T:CD19^+^ B ratio (T:B) was calculated. Alongside, immune index was also assessed from (CD3^+^+CD19^+^+CD56^+^)/CD3^-^CD19^-^CD56^-^.

#### 2.2.13. Histopathological analysis of tissues

Following fixation in 10% formalin, tissue samples were dehydrated by standard procedure and further embedded in paraffin. For hematoxylin and eosin (H&E) of bone with marrow, formalin-fixed femurs were first decalcified using 14% EDTA (pH adjusted to ∼7.4 using ammonium hydroxide) for a period of ∼15 days. Following decalcification, femurs were washed with sterile PBS for 30 min and two times again with ddH_2_O for 30 min each to remove excess EDTA. Further, the samples were dehydrated and embedded in paraffin wax. Tissue sections were cut at 5 μm thickness using Leica RM2125RT microtome. All histopathological evaluation was performed by H&E staining. Stained slides were visualized under Zeiss Primovert inverted microscope equipped with Tucsen GT 5.0 5MP USB2.0 CMOS Camera, using 10X, 20X and 40X objective lens.

#### 2.2.14. Immunofiuorescence analysis of bone marrow

For immunofluorescence analysis of the bone marrow smear, freshly isolated bone marrow was fixed in 4% paraformaldehyde for 15 - 20 min and then smeared on glass slide. Following air-drying, slides were washed two times with sterile PBS (5 min each) and further permeabilized with 0.1% Triton-X for 3 min. Following permeabilization, cells were further washed with sterile PBS for two times (5 min each) and 3% bovine serum albumin (BSA) was used to block non-specific antibody binding by incubating for 30 min. Further cells were washed with sterile PBS for three times, 5 min each. Then, FITC-CD45 and PE-CD34 antibodies were used at concentrations recommended by manufacturer and cells were incubated in a dark humidified chamber for 1 h at RT. Cells were further washed with sterile PBS for three times (5 min each) and 300 nM DAPI was used to stain nuclei. Incubation was done for 3 min in dark at RT and further washed three times with sterile PBS (5 min each). Following nuclear staining, 90% glycerol was used to mount cover slip and edges were sealed with transparent nail-polish. Slides were visualized under Olympus FLUOVIEW FV4000 Confocal Laser Scanning Microscope, using 20X and 40X objective lenses.

#### 2.2.15. Experimental strategy for blood Hg pharmacokinetics

For the blood Hg pharmacokinetic (PK) study, a single dose of α-Mercurin (1.6 mg/kg body weight) was administered intravenously to rats (n=3). Blood samples were collected at 0 min, 15 min, 1 h, 2 h, 4 h, and 24 h in K2 EDTA blood collection tubes. Immediately after collection, 100 µL of each blood sample was transferred to polypropylene tube pretreated with 10 µL of concentrated HNO₃ (69%). The Hg content was subsequently measured using CV-AAS. From the calculated values, we further estimated C_max_, T_max_, k (elimination; log-linear regression 1-24 h), t_1/2_ (0.693/k), AUC_0-24 h_ (linear trapezoidal), and AUC_0-∞_ (AUC_0-24 h_ + C_last_/k).

#### 2.2.16. Experimental strategy for urinary/fecal Hg elimination study

For the urinary and fecal Hg elimination study, α-Mercurin was administered intravenously at a dose of 1.6 mg/kg body weight, three times weekly for two weeks (Healthy, n=3; α-Mercurin, n=3). Urine and fecal samples were collected once weekly. Baseline samples were collected at week 0 (day 1, before treatment initiation), followed by sample collection at week 1 (day 7, 24 h after the third intravenous dose) and week 2 (day 14, 24 h after the sixth intravenous dose). Following the second week of treatment, animals underwent a one-week clearance/washout period without further treatment. Urine and fecal samples were collected again at week 3 (day 21) following the washout period. Hg content was measured using CV-AAS.

#### 2.2.17. Mercury (Hg) estimation by CV-AAS

Mercury (Hg) quantification was performed using cold vapour atomic absorption spectroscopy (CV-AAS) on a Varian AA240FS atomic absorption spectrophotometer (AAS) equipped with a Hg hollow cathode lamp and a quartz cold vapour cell. Working standards of Hg were freshly prepared each day by serial dilution of the stock solution (1000 mg/L; 1000 ppm) using the same acid matrix with which the samples and calibration curves were generated. A stannous chloride (SnCl₂) solution (25% w/v) was prepared freshly each day by dissolving in 20% hydrochloric acid and used as the reducing agent. All solutions were prepared using ddH_2_O.

Excised rat organs (brain, thymus, heart, lung, liver, spleen, kidney, lymph and femur bone) and feces were weighed accurately (80–120 mg for tissues and 500–600 mg for feces; wet weight). For blood and urine samples, 100 µl of samples were collected in polypropylene tubes pretreated with 10 µl concentrated HNO_3_ (69%). Further, all the samples were digested with concentrated HNO_3_ (69%), heating at 85°C until complete digestion and formation of clear solution. Following digestion, samples were cooled to RT and diluted with ddH_2_O to a final volume of 20 mL, and analyzed on the same day. Hg concentrations were determined from standard curve and corrected for dilution. Final results were expressed as µg/kg wet tissue for tissue/feces and µg/Liter for blood/urine.

#### 2.2.18. GSH/GSSG assay of liver and kidney tissues

For the analysis of tissue (liver and kidney) GSH:GSSG ratio, freshly isolated tissue sections from two animal groups (healthy control, n=3; α-Mercurin control, n=3) were immediately snap-frozen in liquid nitrogen (−196 ⁰C). Further the samples were weighed within 40 – 50 mg, and homogenized using cold homogenization buffer (50 mM phosphate, pH 7, 1 mM EDTA) with probe sonicator keeping on ice, as instructed by manufacturer. Samples were then centrifuged at 10,000×g for 5 min at 4 ⁰C. Following centrifugation, supernatant was collected and redistributed in two tubes. For GSSG measurement, one tube was incubated with instructed scavenger, while for total glutathione measurement samples were further diluted using 1X assay buffer. Following that, deproteination was done by 5% meta- Phosphoric Acid (MPA) solution and centrifuged at 14,000×g for 5 min at RT. Next, supernatant was collected and diluted with 1X assay buffer as instructed and distributed in clear flat-bottom 96-well plates and 100 μL of working reagent was added to each sample as instructed by manufacturer. Further, optical density (OD) was recorded for each well at 412 nm at zero and 10 minutes using Tecan Infinite 200 PRO Multimode Microplate Reader. From the OD values, GSSG and GSH concentrations of the samples were measured as per manufacturer’s instruction.

#### 2.2.17. Statistical analysis

All experimental data are presented as median or mean ± SEM with interquartile range (IQR; 25th–75th percentile). Box plots represent median values with IQR, and whiskers indicate minimum-maximum values. Statistical analyses were performed using GraphPad Prism (version 8.4.2) and Microsoft Excel. For comparisons between two independent groups, unpaired two-tailed t-tests were used for parametric data, while non-parametric data were analyzed using Mann-Whitney U test. For multiple group comparisons, one-way ANOVA followed by post-hoc Tukey tests was used for parametric datasets. Due to unequal sample sizes and survival-associated attrition across timepoints, analyses at individual timepoints were performed independently using non-parametric Kruskal-Wallis test with Dunn’s multiple comparisons test. Time-to-humane-endpoint for survival analysis was performed using the Kaplan-Meier method, and differences between groups were evaluated using the log-rank (Mantel-Cox) test. Median survival with range is reported. A p-value <0.05 was considered statistically significant, with significance levels indicated as, *p <0.05, **p <0.01, ***p <0.001, and ****p <0.0001.

## 3. RESULTS

### 3.1. ENU predominantly induces ALL

ENU administration induced leukemia in 31 of 46 animals, with ALL being the predominant subtype (26 of 31 animals). Among these, 24 animals met the criteria for advanced disease, with peripheral blast count ≥35% and WBC count ≥25×10^3^/mm^3^, and we selected these animals for further therapeutic evaluation (Figure 1A).

**Figure 1.**
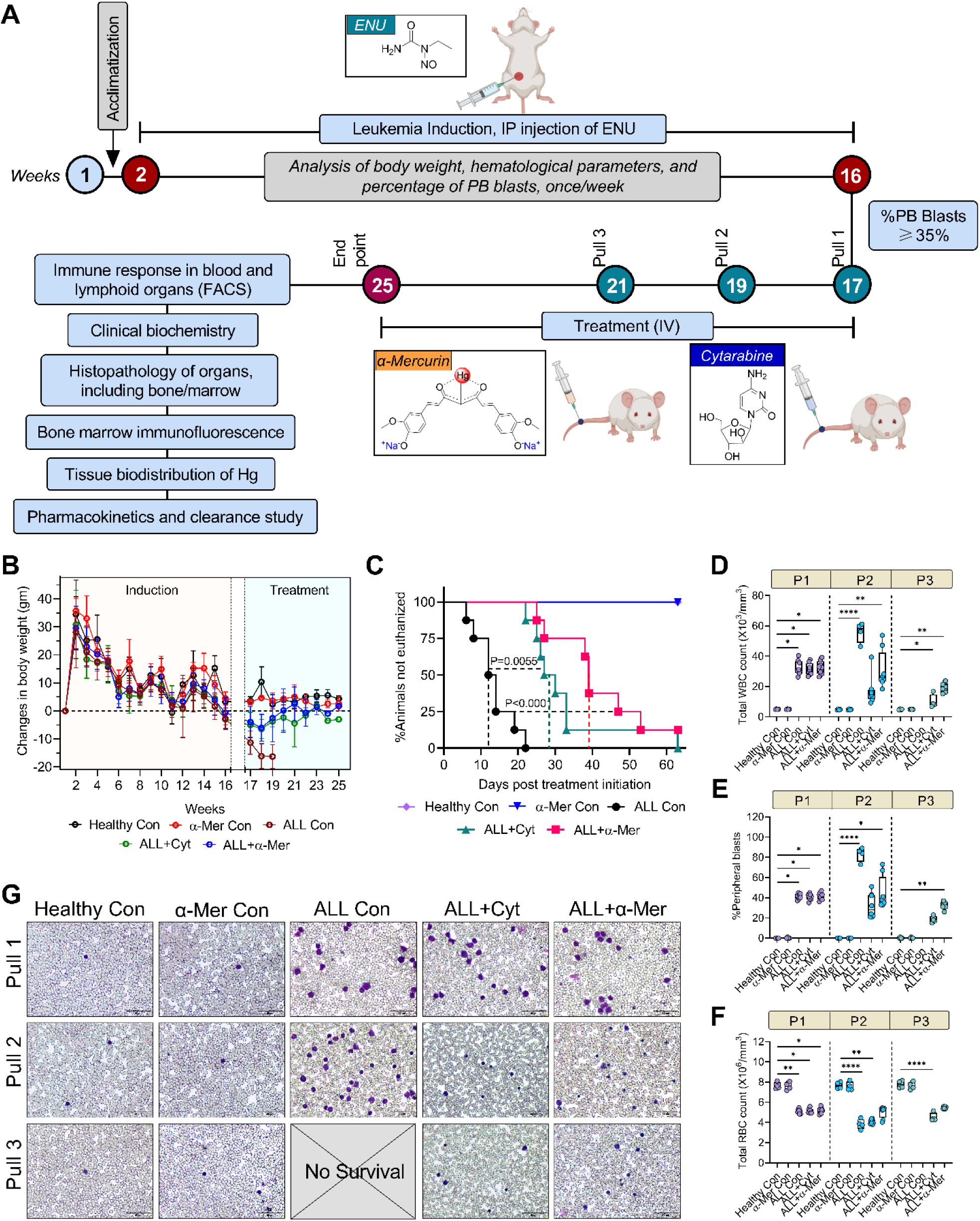
Overview of experimental design and treatment-associated improvement on ALL progression following treatment. **A.** Schematic overview of the experimental design, including ALL induction, treatment schedule and terminal analyses. **B.** Body weight changes (g) during disease progression and treatment. ALL induction results in progressive weight loss from week 2 to week 16, whereas treatment with α-Mercurin or cytarabine attenuates this decline and partially restored body weight toward control levels. **C.** Kaplan-Meier survival analysis showing time-to-humane-endpoint. ALL controls reached endpoint rapidly (13 days), compared with cytarabine (28.5 days) and α-Mercurin (39 days). **D-F.** Hematological parameters including total WBC (**D**), peripheral blasts (**E**) and total RBC (**F**). ALL controls showing rapid disease progression and no survival to P3. For extended data, see Figure S4 and Table S1. All data are presented as median with IQR (25- 75 percentile) in box plots; whiskers represent minimum-maximum values. Kruskal-Wallis test followed by Dunn’s multiple comparisons test was performed for statistical analysis. Sample sizes (n) varied across timepoints (P1: n = 4-8; P2: n = 4-8; P3: n = 0-6) due to survival-associated attrition. Statistical significance is denoted as *p ≤ 0.05, **p ≤ 0.01, ***p ≤ 0.001, ****p ≤ 0.0001. **G.** Representative peripheral blood smears at Pull 1, Pull 2 and Pull 3, illustrating disease progression and therapeutic response. ALL controls suggest increased blast population and abnormal morphology, whereas treated groups exhibit reduced blast burden. Scale bar: 50 µm. Pull 1: P1 (day 1, just before treatment), Pull 2: P2 (day 14, post-treatment), Pull 3: P3 (day 28, post-treatment).

### 3.2. Improved body weight profile following α-Mercurin treatment

We observed comparable body weight across all groups during the initial 12–13 weeks, indicating normal growth and absence of systemic toxicity during this period (Figure 1B). With progression to advanced ALL at ∼15 weeks, ALL controls showed a progressive decline in body weight. Both α-Mercurin and cytarabine treated ALL showed recovery of body weight during the treatment period. In contrast, ALL controls continued to lose body weight and reached the predefined humane endpoints by ∼20^th^ week. Overall, we observed better body weight maintenance in both ALL-treated groups comparing to ALL controls.

### 3.3. α-Mercurin prolongs survival

We performed Kaplan-Meier analysis to evaluate the effect of treatment on disease progression in ALL-bearing rats, using the predefined humane endpoint as the event (Figure 1C). ALL controls showed rapid disease progression, with a median time-to-humane-endpoint of 13 days (range, 6–22 days). Cytarabine treatment delayed disease progression and increased the median time-to-humane-endpoint to 28.5 days (range, 22–63 days). α-Mercurin treatment further extended the median time-to-humane-endpoint to 39 days (range, 25–63 days). We therefore observed longer survival in the α-Mercurin-treated ALL compared with both ALL controls and cytarabine-treated ALL.

### 3.4. α-Mercurin treatment suggests sustained leukemia control and progressive improvement of hematological indices

We monitored hematological parameters throughout the study and found stable indices in healthy and α-Mercurin controls (Figure 1D–1G and Figure S4A–S4G). At Pull 1, we found a similar disease profile in all ALL-bearing animals, with increased peripheral blasts and WBC counts, thrombocytopenia, anemia, and reduced lymphoid and myeloid components (Table S1). At Pull 2, ALL controls showed further progression of disease. Levels of peripheral blast increased from ∼41.5% to ∼85% at Pull 2, while WBC counts increased from ∼30.7×10^3^/mm^3^ to ∼58.0×10^3^/mm^3^. We also observed a decline in platelet and Hb levels from ∼2.71×10^5^/mm^3^ to ∼1.43×10^5^/mm^3^ and from ∼8.05 to ∼6.06 g/dL, respectively. RBC, lymphocyte, and neutrophil levels also decreased during disease progression. Cytarabine treatment reduced peripheral blasts from ∼41% at Pull 1 to ∼17.5% at Pull 3 and WBC count from ∼30.2×10^3^/mm^3^ to ∼8.4×10^3^/mm^3^. We also observed recovery of neutrophils and lymphocytes, which reached ∼16% and ∼53%, respectively at Pull 3. However, RBC and Hb levels remained low at ∼4.35×10^6^/mm^3^ and ∼6.3 g/dL, respectively. Platelet counts showed partial recovery at Pull 3. In contrast, α- Mercurin treatment reduced peripheral blasts from ∼42% at Pull 1 to ∼31.5% at Pull 3. We also observed recovery of lymphocyte and neutrophil levels to ∼43.5% and ∼17.5%, respectively. Platelet counts increased from ∼2.34×10^5^/mm^3^ at Pull 1 to ∼2.65×10^5^/mm^3^ at Pull 3, while RBC and Hb levels reached ∼5.47×10^6^/mm^3^ and ∼8.19 g/dL, respectively.

### 3.5. Terminal hematological and bone marrow evaluation suggest hematopoietic recovery

We further evaluated terminal peripheral blood parameters to assess the hematological response to treatment (Figure 2A and 2B). ALL controls showed a severe leukemic phenotype comparing to healthy controls, with peripheral blasts of ∼85.5% and WBC counts of ∼59.2×10^3^/mm^3^ (P=0.0011). We also found marked thrombocytopenia (∼0.88×10^5^/mm^3^; P=0.0003), anemia with RBC counts of ∼3.21×10^6^/mm^3^ (P=0.0006) and Hb of ∼5.16 g/dL (P=0.0024), along with reduced lymphocyte (∼9%), neutrophil (∼4%), and monocyte (∼1%) levels. We also observed dense lymphoblast infiltration in the bone marrow with marked loss of normal hematopoietic compartments (Figure 2C and Figure S4H). Cytarabine treatment reduced peripheral blasts to ∼13.5% (P<0.024) and WBC counts to ∼11×10^3^/mm^3^ comparing to ALL controls. We also found partial recovery of platelet counts to ∼1.97×10^5^/mm^3^. However, RBC and Hb levels remained low at ∼4.45×10^6^/mm^3^ and ∼5.93 g/dL, respectively. Bone marrow examination showed reduced leukemic infiltration following cytarabine treatment, but we observed relatively hypocellular marrow with limited erythroid recovery. In contrast, α-Mercurin treatment reduced peripheral blast burden to ∼30.5% compared with ∼85.5% in ALL controls. We also observed a reduction in WBC count to ∼16.5×10^3^/mm^3^. At the same time, α-Mercurin treatment improved erythroid and megakaryocytic parameters, with RBC counts of ∼5.60×10^6^/mm^3^, Hb of ∼8.01 g/dL, and platelet counts of ∼2.6×10^5^/mm^3^. We further observed reduced leukemic infiltration in the bone marrow, together with scattered reappearances of erythroid precursors and megakaryocytes.

**Figure 2.**
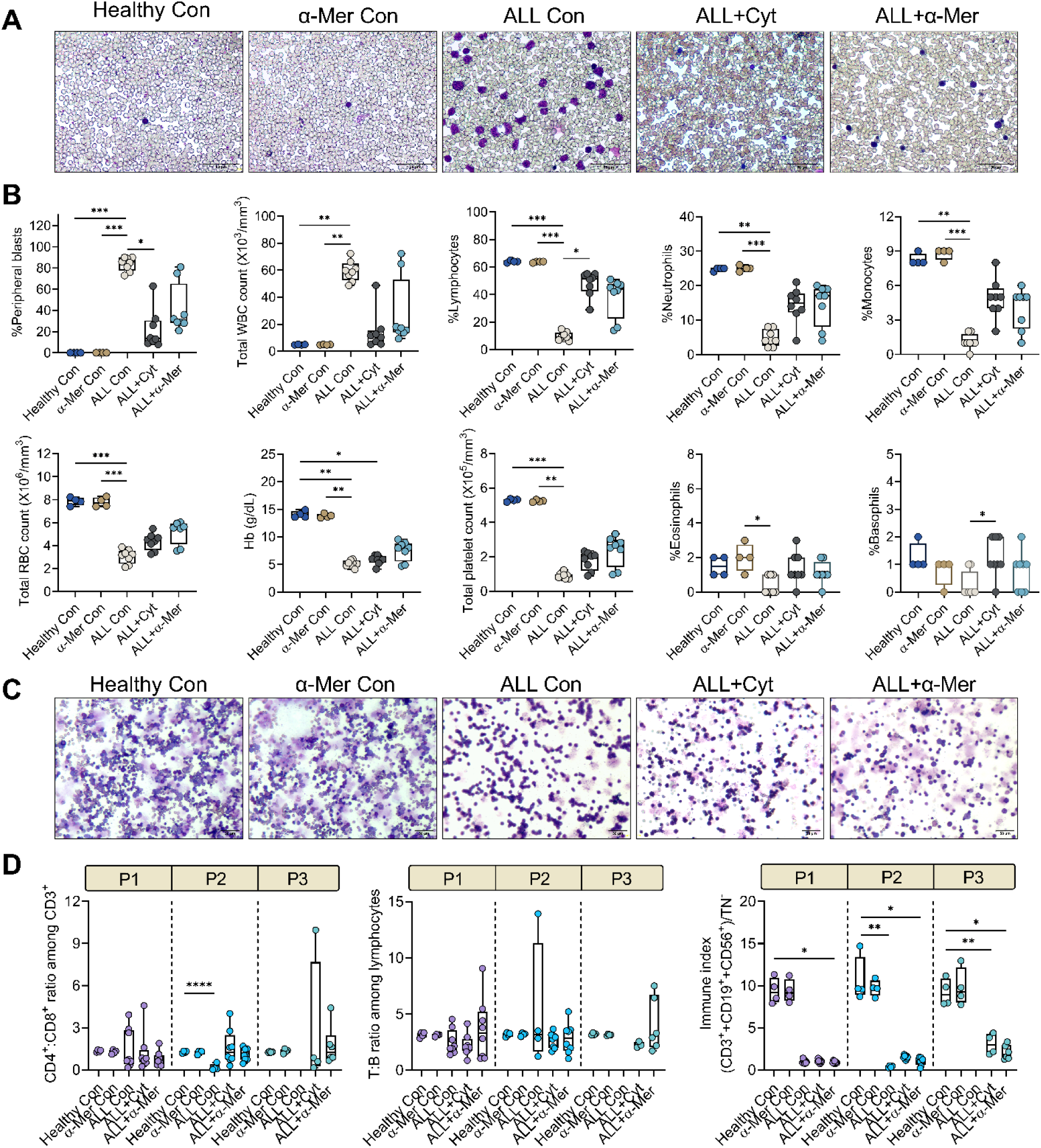
α-Mercurin reduces leukemic burden with improvement of hematopoietic and immune health. **A.** Representative peripheral blood smears collected prior to euthanasia showing marked leukocytosis and circulating blasts in ALL controls. Treatment with α-Mercurin significantly reduces leukocytosis and blast burden. Scale bar: 50 µm. **B.** Graphical depiction of complete blood count (CBC) parameters from peripheral blood samples across all groups at endpoint. **C.** Representative Leishman-stained bone marrow smears showing extensive immature blasts and effacement of normal architecture in ALL controls. Cytarabine-treated animals exhibit marked cytoreduction with associated erythroid suppression, whereas α-Mercurin treatment suggests reduced leukemic burden with recovery of multilineage hematopoiesis. Scale bar: 50 µm. **D.** Immune profiling of peripheral blood at P1, P2 and P3, showing improvement of immune balance following α- Mercurin treatment. Panels include CD4⁺:CD8⁺ and CD3⁺ T:CD19⁺ B-cell (T:B) ratio along with immune index. ALL controls did not survive to P3. For extended data, see Figure S6 and Table S2. All data are presented as median with IQR (25-75 percentile) in box plots; whiskers represent minimum-maximum values. Kruskal-Wallis test followed by Dunn’s multiple comparisons test was performed for statistical analysis. Sample sizes (n) varied across timepoints (P1: n = 4-8; P2: n = 4-8; P3: n = 0-6; Terminal: n = 4-8) due to survival-associated attrition. Statistical significance is denoted as *p ≤ 0.05, **p ≤ 0.01, ***p ≤ 0.001, ****p ≤ 0.0001. Pull 1: P1 (day 1, just before treatment), Pull 2: P2 (day 14, post-treatment), Pull 3: P3 (day 28, post-treatment).

### 3.6. α-Mercurin suggests progressive improvement in peripheral immune parameters

At Pull 1, we observed marked changes in peripheral immune parameters in ALL controls compared with healthy controls (Figure 2D). ALL controls showed ∼23.15% CD3⁺ T cells, with CD8⁺ cells of ∼42.95%, higher than CD4⁺ cells of ∼29.96%, resulting in a reduced CD4⁺:CD8⁺ ratio of ∼0.7 comparing to ∼1.3 in healthy controls. We also found reduced CD19⁺ cells of ∼10.2% and an increased triple-negative population of ∼51.5% (P=0.0248), with the immune index decreasing to ∼0.95 (P=0.0696) (Figure S6 and Table S2). At Pull 2, we observed further deterioration of peripheral immune parameters in ALL controls. CD3⁺, CD4⁺, and CD19⁺ populations decreased to ∼15.75%, ∼7.14%, and ∼5.63%, respectively, while CD8⁺ cells increased to ∼74.53%. The immune index also decreased to ∼0.26 (P=0.005), comparing to healthy controls. Cytarabine treatment increased CD3⁺ and CD4⁺ populations to ∼34.25% and ∼50%, respectively, and CD19⁺ cells to ∼12.9%, resulting in an immune index of ∼1.60 comparing to ALL controls. α-Mercurin treatment also increased CD3⁺ and CD4⁺ populations to ∼30.5% and ∼41.8%, respectively, while CD19⁺ cells increased to ∼9.96% and the immune index to ∼1.23. We also observed a reduction in the triple-negative population following both treatments compared with ALL controls. At Pull 3, cytarabine treatment further increased CD3⁺ and CD19⁺ populations to ∼40.86% and ∼17.19%, respectively, with the immune index reaching ∼3.01. α-Mercurin treatment increased CD3⁺, CD4⁺, and CD19⁺ populations to ∼35.05%, ∼46.6%, and ∼13.19%, respectively, with the immune index increasing to ∼2.2. We observed stable and comparable immune parameters in the healthy and α-Mercurin controls throughout the study.

### 3.7. α-Mercurin improves immune parameters across hematopoietic and lymphoid compartments

Terminal immune profiling further showed changes in immune cell populations across peripheral blood (PB), bone marrow, thymus, lymph nodes, and spleen (Figure 3A–3F and Figures S7–S11). In PB (Figure 3A and Figure S7) of ALL controls, we found reduced CD3⁺ and CD4⁺ T-cell population of ∼16.1% (P=0.0006) and ∼7.27%, respectively; along with increased CD8⁺ cells (∼66.35%), resulting in a low CD4⁺:CD8⁺ ratio of ∼0.1 comparing to healthy controls. We also found reduced CD19⁺ B cells (∼4.18%, P=0.003) and an increased triple-negative population (∼79.08%, P=0.0001). Comparing to ALL controls, cytarabine treatment increased CD3⁺, CD4⁺, and CD19⁺ population to ∼39.25%, ∼39.06%, and ∼16.74% (P=0.0115), respectively, and improved the CD4⁺:CD8⁺ ratio to ∼0.8 while reducing triple-negative cells to ∼31.54% (P=0.0337). α-Mercurin treatment increased CD3⁺ and CD4⁺ population to ∼33.7% and ∼37.7%, respectively, and reduced CD8⁺ cells to ∼43% comparing to ALL controls. We also observed an increase in the CD4⁺:CD8⁺ ratio to ∼0.94 and CD19⁺ cells to ∼11.52%, while the triple-negative population decreased to ∼39.54%. We observed similar changes in bone marrow (Figure 3C and Figure S8). ALL controls showed reduced CD19⁺ cells (∼10.76%) and increased CD8⁺ cells (∼58.58%), with a CD4⁺:CD8⁺ ratio of ∼0.32. Comparing to ALL controls, cytarabine treatment increased CD19⁺ cells to ∼38.2% (P=0.0152) and the CD4⁺:CD8⁺ ratio to ∼1.7. α- Mercurin treatment increased CD19⁺ cells to ∼27.39% and reduced CD8⁺ cells to ∼38.6%, resulting in a CD4⁺:CD8⁺ ratio of ∼0.65. We also evaluated thymic immune population and found recovery following α-Mercurin treatment (Figure 3D and Figure S9). ALL controls showed marked reductions in CD3⁺ thymocytes (∼21.95%, P=0.0006), CD4⁺ (∼2.31%), and CD8⁺ (∼2.7%) population, with a CD4⁺:CD8⁺ ratio of ∼0.35 comparing to healthy controls. Comparing to ALL controls, cytarabine treatment increased CD3⁺ thymocytes to ∼54.23%, but DP (CD4⁺CD8⁺) cells remained at ∼24.45% (P=0.0351). In comparison, α-Mercurin treatment increased CD3⁺ thymocytes to ∼56.35% and DP cells to ∼53.88%. In lymph nodes of ALL controls (Figure 3E and Figure S10), we found reduced CD3⁺ (P=0.0172) and CD4⁺ population of ∼23.85% and ∼12.4%, respectively; along with an increased triple-negative population (∼51.33%, P=0.0054) and a reduced CD4⁺:CD8⁺ ratio (∼0.42) comparing to healthy controls. α-Mercurin treatment increased CD3⁺, CD4⁺, and CD19⁺ population to ∼37.4%, ∼36.01%, and ∼25.45%, respectively, and restored the CD4⁺:CD8⁺ ratio to ∼1.3 comparing to ALL controls. Cytarabine treatment also increased these population, with CD3⁺, CD4⁺, and CD19⁺ cells reaching ∼39.36%, ∼31.38%, and ∼14.23%, respectively, and the CD4⁺:CD8⁺ ratio reaching ∼0.8. In the spleen, ALL controls showed reduced CD19⁺ cells (∼20.34%) and an increased triple-negative population (∼31.2%) and CD3⁺ cells ∼39.85% while CD4^+^:CD8^+^ remained ∼1.28 (Figure 3F and Figure S11). α-Mercurin treatment increased CD4⁺ and CD19⁺ cells to ∼42.42% and ∼29.73%, respectively, and increased the CD4⁺:CD8⁺ ratio to ∼1.77. In contrast, cytarabine treatment produced a more CD8⁺-dominant profile, with CD8⁺ cells at ∼38.09%, a CD4⁺:CD8⁺ ratio of ∼0.53, and CD19⁺ cells at ∼10.49% (Table S3).

**Figure 3.**
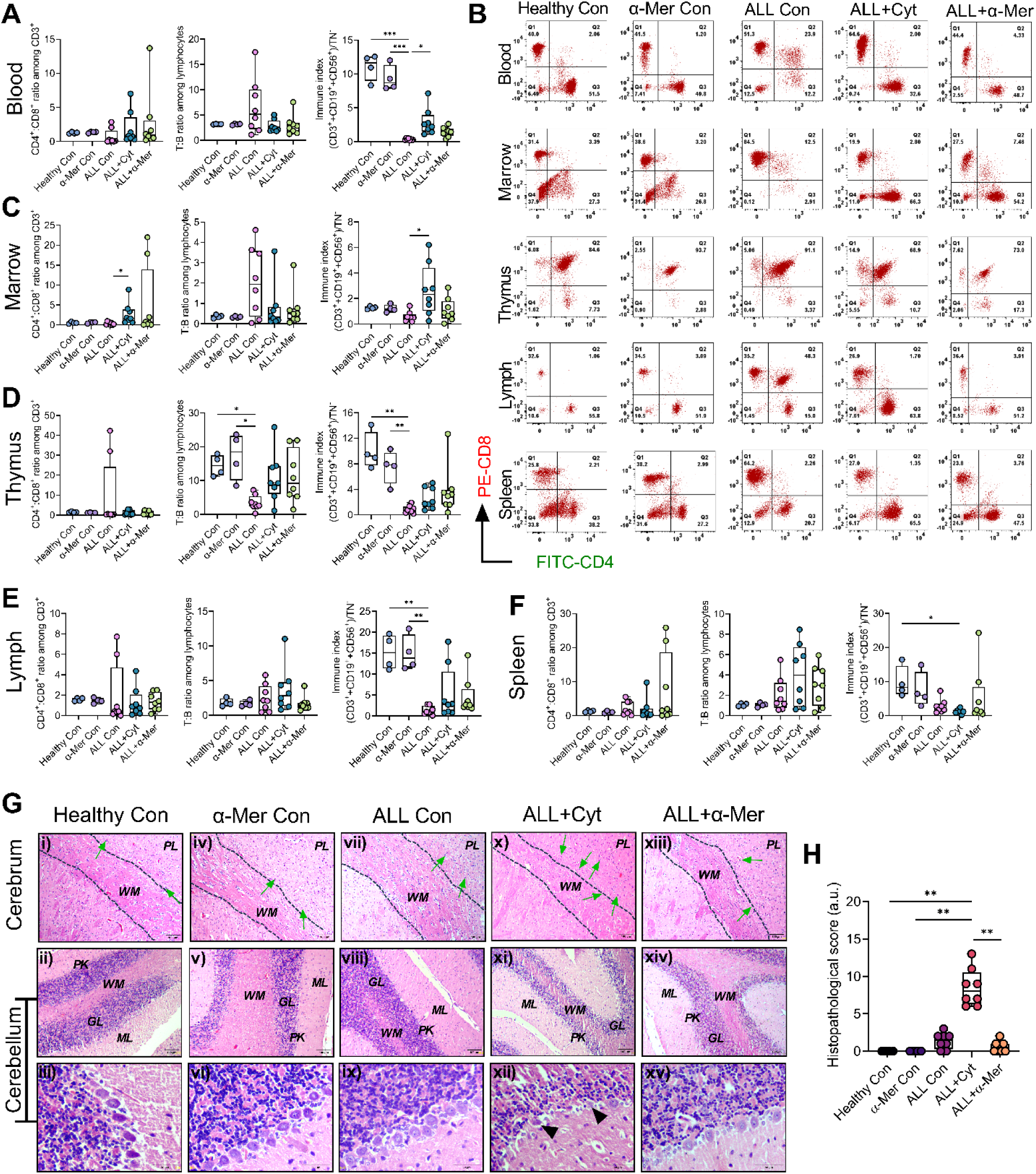
α-Mercurin suggests improvement of immune health across lymphoid compartments with preserved brain architecture. **A.** Peripheral blood immune profiling at endpoint showing proportions of major immune cell populations and derived ratios across experimental groups. **B.** Distribution of CD3⁺ T-cell subsets (CD4^+^, CD8^+^, CD4^+^CD8^+^ and CD4^-^CD8^-^) across peripheral blood, bone marrow, thymus, lymph nodes and spleen. **C-F.** Flow cytometric analysis of single-cell suspensions from bone marrow (**C**), thymus (**D**), lymph nodes (**E**) and spleen (**F**), showing immune balance indices, including CD4⁺:CD8⁺ ratio, CD3⁺ T:CD19⁺ B-cell (T:B) ratio and comprehensive immune index [(CD3^+^+CD19^+^+CD56^+^)/(CD3^-^CD19^-^CD56^-^)] to assess overall immune health improvement. ALL controls exhibit systemic immune dysregulation characterized by T-cell depletion, reduced marrow B-cell populations, CD8^+^ skewing and impaired immune indices across compartments. Cytarabine treatment partially improved marrow B-cell and peripheral CD4^+^ populations, whereas α-Mercurin suggests broader immune balance, including normalization of thymic and splenic compartments. For extended data, see Figure S7–S11 and Table S3. **G.** Representative hematoxylin and eosin (H&E) stained sections of cerebrum and cerebellum across all experimental groups. ALL controls exhibit minimal neuronal shrinkage and scattered microgliosis without leukemic infiltration. Cytarabine treatment displays neuronal degeneration, microgliosis, demyelination and Purkinje cell injury, indicating treatment-associated neurotoxicity. In contrast, α-Mercurin maintains intact neuronal morphology and cerebellar architecture with no detectable neurotoxic lesions. Scale bar: 50, 100 µm. ML: Molecular Layer, GL: Granular Layer, PL: Pyramidal Layer, AM: Arachnoid Mater, WM: White Mater, PM: Pia Mater, PK: Purkinje cells, Green arrow: Microgliosis, Black arrowhead: Purkinje degeneration. **H.** Quantitative histopathological scoring of brain tissue based on neuronal degeneration, white matter demyelination, microgliosis, vacuolation and Purkinje cell loss; data suggests no neurotoxicity and CNS safety of α-Mercurin at given dose. For extended data, see Figure S12. All data are presented as median with IQR (25-75 percentile) in box plots; whiskers represent minimum-maximum values. Kruskal-Wallis test followed by Dunn’s multiple comparisons test was performed for statistical analysis. Statistical significance is denoted as *p ≤ 0.05, **p ≤ 0.01, ***p ≤ 0.001, ****p ≤ 0.0001.

### 3.8. α-Mercurin preserves brain architecture and myocardial integrity

We evaluated histopathology of the cerebrum and cerebellum to assess any neurological toxicity (Figure 3G–3H and Figure S12). We observed intact neuronal architecture in both healthy and α-Mercurin controls, with preserved cortical layers, Purkinje cells, and white matter. ALL controls showed only mild changes, including occasional neuronal shrinkage and limited microglial activation, without leukemic infiltration, suggesting total histopathological score (HS) of 1.5 (IQR: 0.25–2). In contrast, cytarabine-treated ALL showed neuronal degeneration, white matter demyelination, molecular layer vacuolation, and Purkinje cell loss, resulting in a higher total HS of 8 (IQR: 6.25–10.5) comparing to healthy controls (P=0.0013). α-Mercurin treatment maintained normal neuronal morphology without evident gliosis or demyelination, with a total HS of 0.5 (IQR: 0–1).

We also evaluated cardiac histology to assess myocardial safety (Figure 4A–4B and Figure S13). We observed preserved myocardial structure in healthy and α-Mercurin controls, while ALL controls showed mild edema and scattered inflammatory cell congestion within intramyocardial vessels, with a total HS of 1 (IQR: 1–2). Cytarabine treatment produced marked cardiac alterations, including myocyte necrosis, myocarditis, inflammatory changes, and fiber disorganization, with a total HS of 7 (IQR: 6–8) comparing to healthy controls (P=0.0002). In contrast, α-Mercurin treatment maintained myocardial architecture with only minimal inflammatory changes, resulting in a total HS of 1 (IQR: 0.25–1).

**Figure 4.**
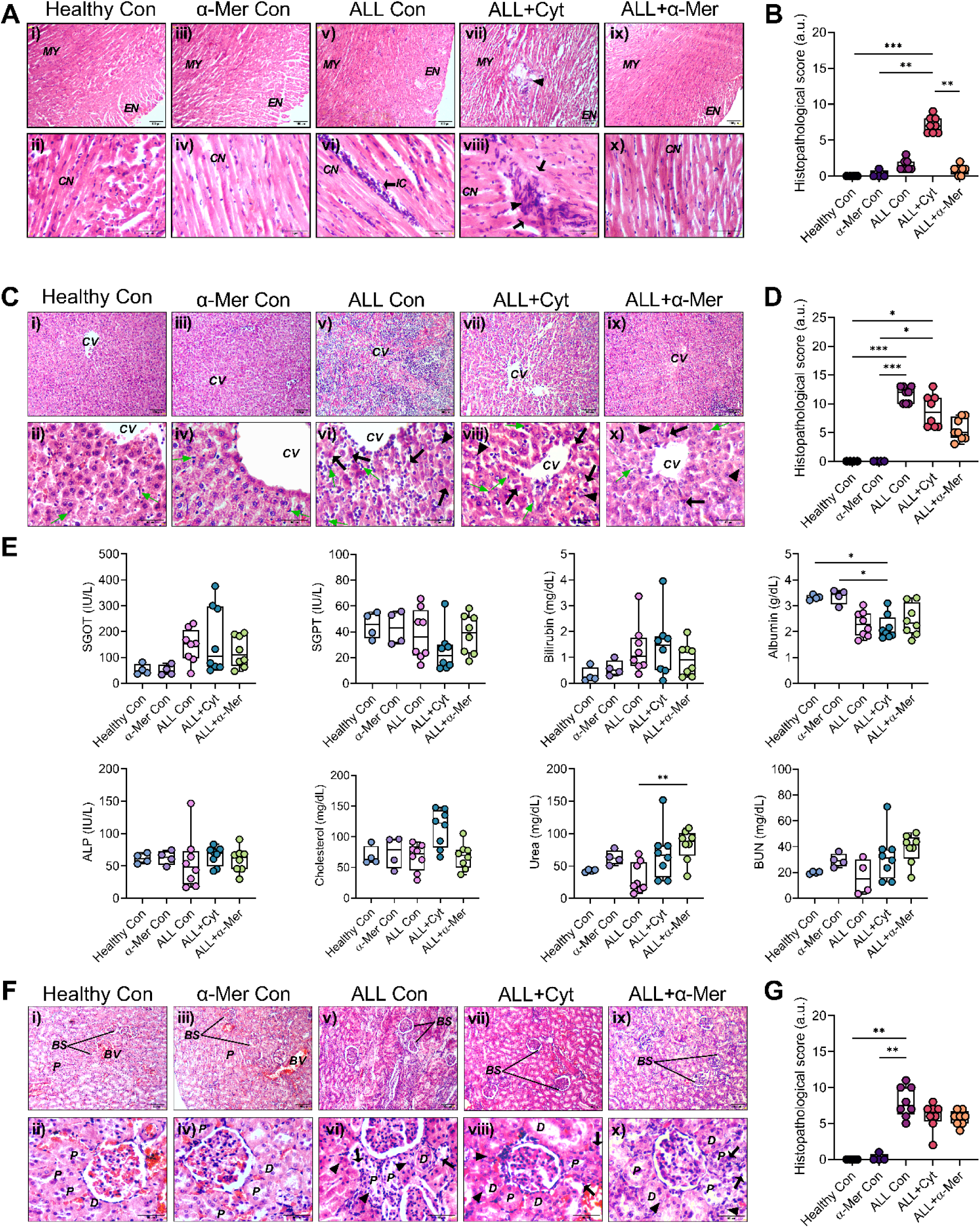
α-Mercurin demonstrates preserved cardiac, hepatic and renal tissue integrity with improved biochemical parameters. **A.** Representative hematoxylin and eosin (H&E)-stained cardiac sections across all experimental groups. ALL controls showing mild interstitial edema, focal inflammatory cell infiltration and vascular congestion. Cytarabine treatment exhibits pronounced myocardial injury as shown by myofiber vacuolation, patchy necrosis and endothelial swelling with dense inflammatory infiltrates, consistent with myocarditis. In contrast, α-Mercurin-treated ALL depicts largely preserved myocardial structure with no significant necrosis or fibrosis and only occasional mild interstitial inflammatory cell infiltration. Scale bar: 50, 100 µm. MY: Myocardium, EN: Endocardium, CN: Central Nucleus, IC: Inflammatory cells, Black arrow: Myofibre necrosis, Black arrowhead: Myocarditis. **B.** Quantitative histopathological scoring of cardiac tissue based on myocarditis, necrosis, myofiber degeneration and myocardial swelling, suggesting no myocardial toxicity of α-Mercurin at given dose. For extended data, see Figure S13. **C.** Representative liver histology from each experimental group with H&E staining. Severe leukemic cell infiltration in ALL controls is observed. Treatment with cytarabine reduces the disease burden, including extramedullary hematopoiesis, simultaneously it also exerts marked hepatocellular toxicity.

### 3.9. α-Mercurin reduces hepatic leukemic burden while preserving liver architecture and improves biochemical parameters

We evaluated liver histology to assess leukemic infiltration and hepatic safety following treatment (Figure 4C–4D and Figure S14). Healthy and α-Mercurin controls showed intact hepatic architecture, with well-organized hepatic cords and normal hepatocyte morphology. In contrast, ALL controls showed extensive sinusoidal infiltration, disruption of lobular architecture, extramedullary hematopoiesis, and Kupffer cell hypertrophy, with a total HS of 12 (IQR: 10–13; P=0.009 vs healthy control). Cytarabine treatment reduced leukemic infiltration but caused evident hepatocellular injury, including hydropic degeneration, focal necrosis, and loss of normal hepatocyte organization with total HS of 8 (IQR: 6–11; P=0.0411 vs healthy control). α-Mercurin treatment reduced hepatic leukemic infiltration while maintaining hepatic architecture, with minimal inflammatory changes and no evident fibrosis or steatosis (total HS: 5, IQR: 4–7.75).

We next evaluated serum biochemical parameters to assess overall organ health (Figure 4E). ALL controls showed increased SGOT of ∼152.3 IU/L and bilirubin of ∼1.04 mg/dL, together with reduced albumin to ∼2.27 g/dL and ALP to ∼48.64 IU/L. Cytarabine treatment reduced SGOT to ∼105.3 IU/L but showed an increase in bilirubin to ∼1.47 mg/dL and lower albumin to ∼2.0 g/dL, while SGPT and ALP were ∼21.65 IU/L and ∼69.87 IU/L, respectively. α-Mercurin treatment reduced SGOT to ∼110.6 IU/L and bilirubin to ∼0.91 mg/dL, while SGPT, ALP, and albumin were ∼39.44 IU/L, ∼62.9 IU/L, and ∼2.3 g/dL, respectively. Cytarabine treatment also significantly elevated cholesterol to ∼122.3 mg/dL compared to ∼62.53 mg/dL in healthy controls, whereas the ALL controls showed ∼72.36 mg/dL. α-Mercurin treatment maintained cholesterol at ∼69.47 mg/dL, close to the level observed in healthy controls. Further, ALL controls showed a reduced urea level of ∼21.15 mg/dL comparing to healthy control of ∼43.16 mg/dL. Cytarabine treatment showed ∼66.76 mg/dL, while ALL animals treated with α-Mercurin elevated the level further to ∼88.09 mg/dL.

### 3.10. α-Mercurin maintains renal and pulmonary tissue integrity

We evaluated kidney histology to assess renal effects following treatment (Figure 4F–4G and Figure S15). Healthy and α-Mercurin controls showed normal glomerular and tubular architecture. ALL controls showed leukemic infiltration, glomerular hypercellularity, and tubular injury, with a total HS of 7.5 (IQR: 6.25–10; P=0.0022 vs healthy control). We also observed congestion of capillaries with lymphocytes and neutrophils, suggesting endocapillary glomerulopathy. Cytarabine treatment reduced leukemic infiltration but showed mild tubular and epithelial changes, with a total HS of 6.5 (IQR: 5.25–7). Similarly, α-Mercurin treatment reduced renal leukemic infiltration and also showed mild tubular stress, with a total HS of 6 (IQR: 5–6.75).

However, no intrinsic hepatotoxicity is detected following α-Mercurin treatment. Scale bar: 50, 100 µm. CV: Central Vein, Green arrow: Kupffer cell, Black arrow: Apoptotic cell, Black arrowhead: Karyorrhexis. **D.** Quantitative histopathological scoring of liver tissues based on hepatocellular injury, extramedullary hematopoiesis, sinusoidal congestion/dilation, Kupffer cell hypertrophy, hyperplasia and steatosis, suggested therapeutic benefit of α-Mercurin treatment. For extended data, see Figure S14. **E.** Clinical biochemistry data across all groups, showing levels for serum SGOT, SGPT, Bilirubin, Albumin, ALP, Cholesterol, Urea, and BUN. ALL induction suggests altered transaminases, bilirubin, albumin and BUN levels. Though cytarabine significantly reduces immature lymphoblasts, but it also exhibits significant hepatic stress. α-Mercurin treatment improves functional hepatic parameters toward normal range, despite mild elevation in Urea and BUN level. **F.** Representative H&E-stained kidney sections across experimental groups. ALL controls exhibit glomerular inflammation, interstitial infiltration and tubular injury. Both cytarabine and α-Mercurin treatments suggest reduced leukemic burden and associated inflammatory changes, with α-Mercurin maintaining overall structural integrity and showing no evidence of intrinsic nephrotoxicity. Scale bar: 50, 100 µm. BS: Bowman’s space, BV: blood vessels, G: glomerulus, P: proximal tubule, D: distal tubule, Black arrow: apoptotic cell, Black arrowhead: karyorrhexis. **G.** Quantitative histopathological scoring of kidney tissues based on endocapillary glomerulopathy, interstitial inflammation, glomerular congestion, tubular injury and atrophy; demonstrating improved renal pathology following treatment. For extended data, see Figure S15. All histopathological score and clinical biochemistry data are presented as median with IQR (25-75 percentile) in box plots; whiskers represent minimum-maximum values. Kruskal-Wallis test followed by Dunn’s multiple comparisons test was performed for statistical analysis. Statistical significance is denoted as *p ≤ 0.05, **p ≤ 0.01, ***p ≤ 0.001, ****p ≤ 0.0001.

We also evaluated lung histology to assess pulmonary tissue changes (Figure 5A–5B and Figure S16). Healthy and α-Mercurin controls showed preserved alveolar architecture. ALL controls showed mild inflammatory changes, vascular congestion, type II pneumocyte hyperplasia, and limited alveolar edema, with a total HS of 8 (IQR: 6.25–8; P=0.0002 vs healthy control). Both treatments reduced these pathological changes. α-Mercurin treatment showed near-preserved lung architecture with a total HS of 3 (IQR: 3–4.75), comparable to cytarabine with total HS of 3 (IQR: 2.25–4).

**Figure 5.**
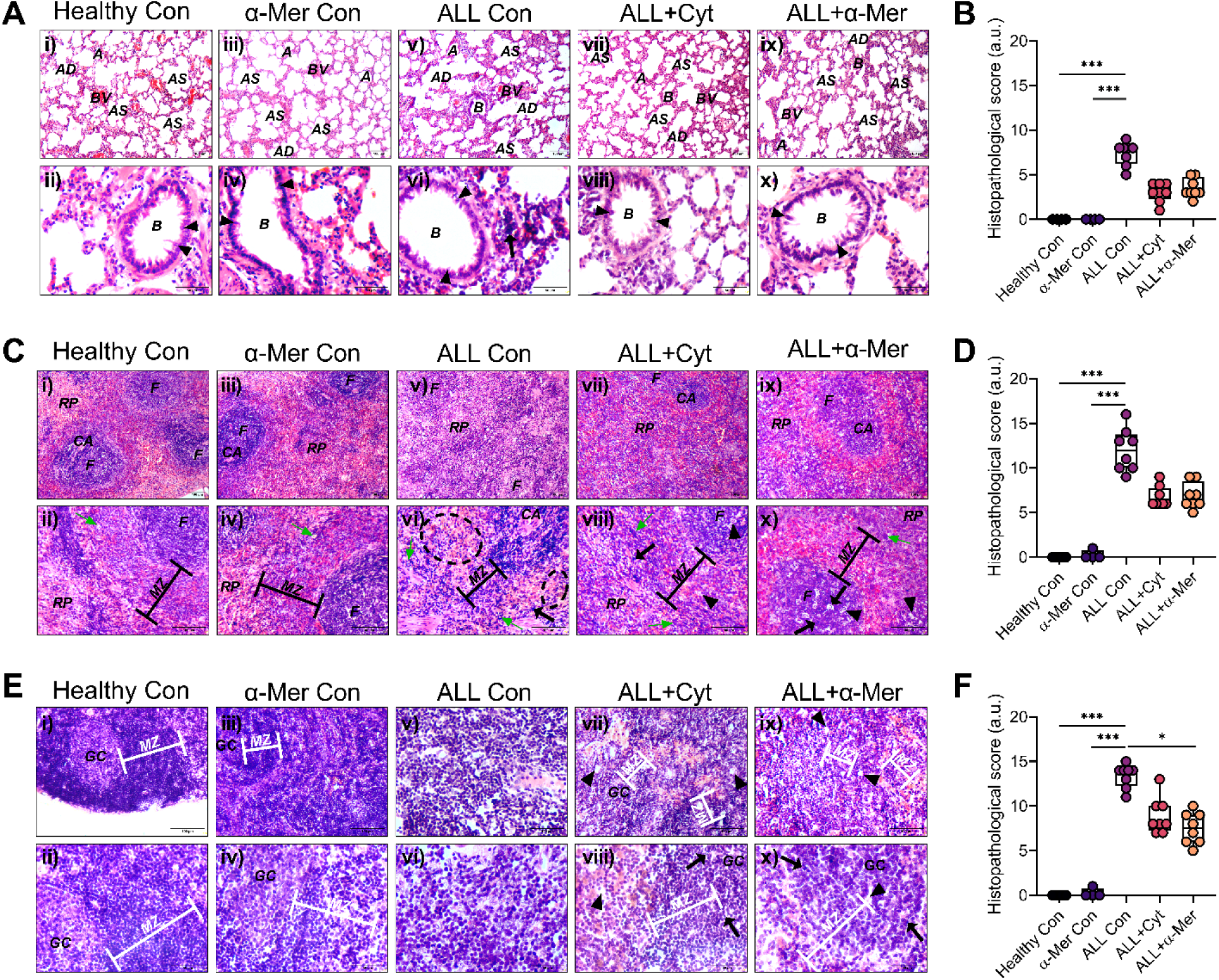
α-Mercurin preserves alveolar, splenic, and lymph node structures with reduced leukemic pathology. **A.** Representative hematoxylin and eosin (H&E)-stained lung sections across experimental groups. ALL controls exhibit mild interstitial inflammation and focal alveolitis. In contrast, treated groups suggest minimal residual changes without evidence of hemorrhage or bronchiolar injury. Scale bar: 50, 100 µm. A: alveolus, AS: alveolar sac, AD: alveolar duct, BV: blood vessel, B: bronchiole, Black arrowhead: Goblet cell. **B.** Quantitative histopathological scoring of lung tissue based on alveolitis, congestion, edema, and Type II pneumocyte hyperplasia, showing no evidence of treatment-associated pulmonary toxicity. For extended data, see Figure S16. **C.** Representative H&E-stained spleen sections across experimental groups. ALL controls indicate diffuse leukemic infiltration, red pulp expansion and loss of white pulp organization. Cytarabine treatment reduces leukemic burden but is associated with lymphoid depletion and hypocellular white pulp, whereas α-Mercurin treatment reduces leukemic infiltration with partial improvement of white pulp architecture. Scale bar: 100 µm. F: follicle, RP: red pulp, CA: central artery, MZ: marginal zone, green arrow: megakaryocytes, black arrow: apoptosis, black arrowhead: vacuoles. Black dashed circle: hemosiderin. **D.** Quantitative histopathological scoring of spleen sections based on leukemic infiltration, white pulp disruption, red pulp expansion/congestion, extramedullary hematopoiesis and necrosis/apoptosis, demonstrating significant disease burden in ALL controls and improvement following treatment. See Figure S17 for extended data. **E.** Representative H&E-stained lymph node sections across experimental groups. ALL controls suggest severe architectural disruption with diffuse leukemic infiltration and loss of follicular organization. Cytarabine treatment results in partial recovery but is associated with lymphoid depletion, whereas α-Mercurin treatment reduces leukemic infiltration and promotes reappearance of follicles and germinal centers. Scale bar: 50, 100 µm. GC: Germinal center, MZ: mantle zone, black arrow: apoptosis, black arrowhead: vacuoles. **F.** Quantitative histopathological scoring of lymph nodes based on structural integrity, leukemic infiltration, follicular status, cellularity, and cell death, demonstrating improved architectural recovery following α-Mercurin treatment for ALL animals. For extended data, see Figure S18. All histopathological scores are presented as median with IQR (25-75 percentile) in box plots; whiskers represent minimum-maximum values. Kruskal-Wallis test followed by Dunn’s multiple comparisons test was performed for statistical analysis. Statistical significance is denoted as *p ≤ 0.05, **p ≤ 0.01, ***p ≤ 0.001, ****p ≤ 0.0001.

### 3.11. α-Mercurin reduces splenic leukemic infiltration and preserves lymph node architecture

Further we evaluated spleen histology to assess leukemic infiltration and changes in splenic architecture following treatment (Figure 5C–5D and Figure S17). Healthy and α-Mercurin controls showed preserved splenic architecture with intact white pulp and organized red pulp. In contrast, ALL controls showed marked splenomegaly, diffuse leukemic infiltration, white pulp atrophy, and extramedullary hematopoiesis, with a total HS of 12 (IQR: 10–13.75; P=0.0002 vs healthy control). Cytarabine treatment reduced leukemic infiltration but only partially restored splenic architecture, with residual structural disorganization and lymphoid depletion, suggesting total HS of 6 (IQR: 6–7.75). α-Mercurin treatment also reduced leukemic infiltration and restored the white pulp and follicles, with better control of extramedullary hematopoiesis, reflecting total HS of 6.5 (IQR: 6–8.5).

We next evaluated lymph node architecture (Figure 5E–5F and Figure S18). Healthy and α-Mercurin controls showed intact cortical, paracortical, and medullary regions with preserved follicles and germinal centres. ALL controls showed extensive leukemic infiltration with loss of follicles, disruption of germinal centres, and expansion of the medullary region, resulting in a total HS of 14 (IQR: 12.25–14; P=0.0002 vs healthy control). Cytarabine treatment reduced leukemic infiltration but showed marked lymphoid depletion, hypocellularity, and residual architectural distortion, with a total HS of 8 (IQR: 7.25–10). α-Mercurin treatment reduced leukemic infiltration and was associated with reappearance of follicles and germinal centres, with a total HS of 7.5 (IQR: 6–9). We also observed mature lymphoid cells with less apparent lymphoid depletion in the α-Mercurin-treated ALL group.

### 3.12. α-Mercurin reduces leukemic infiltration and supports bone marrow and thymic recovery

We further evaluated bone marrow histology to assess leukemic infiltration and hematopoietic changes following treatment (Figure 6A–6B and Figure S19). Healthy and α-Mercurin controls showed heterogeneous marrow with intact bone structure and normal distribution of hematopoietic lineages. ALL controls showed dense blast infiltration suppression of erythroid and megakaryocytic populations, and increased osteoclastic activity, with a total HS of 15.5 (IQR: 13–17; P=0.0022 vs healthy control). Cytarabine treatment markedly reduced blast infiltration and osteoclastic activity but resulted in hypocellular marrow with reduced representation of hematopoietic populations, indicating total HS of 9.5 (IQR: 7.25–10). In contrast, α-Mercurin treatment reduced leukemic infiltration and indicated reappearance of erythroid and myeloid populations with improved marrow heterogeneity and bone remodeling, reflecting total HS of 8.5 (IQR: 6.25–9.75).

**Figure 6.**
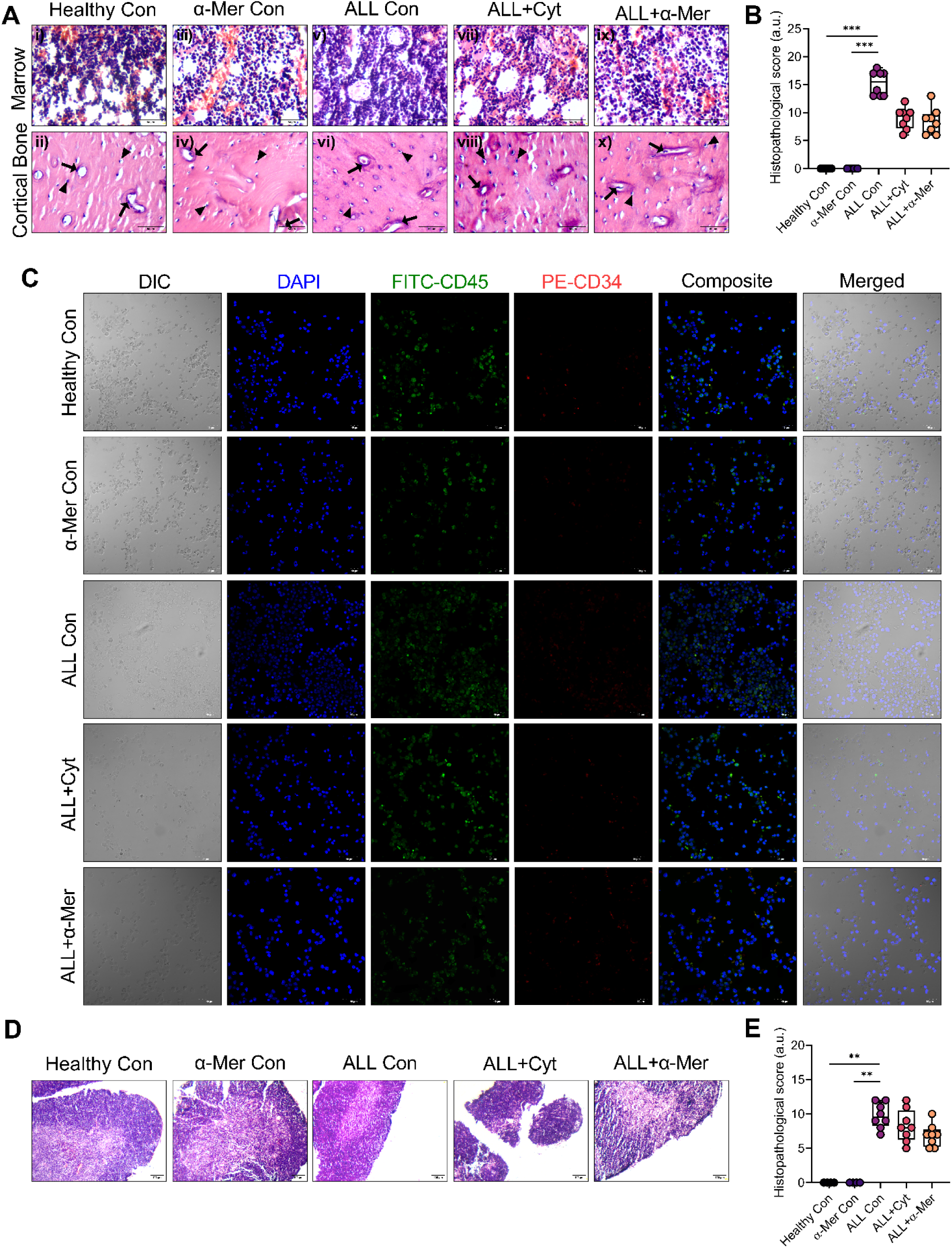
α-Mercurin reduces leukemic burden with improvement of bone marrow architecture and thymic integrity. **A.** Representative hematoxylin and eosin (H&E)-stained sections showing bone marrow and cortical bone architecture across experimental groups. ALL controls suggest diffuse blast infiltration, marrow hypercellularity and disrupted bone remodeling with increased osteoclastic activity. Cytarabine treatment results in blast clearance but induces marrow hypocellularity, whereas α-Mercurin treatment reduces leukemic infiltration while restoring hematopoietic heterogeneity and marrow architecture. Scale bar: 50 µm. Black arrow: Osteoblast, Black arrowhead: Osteoclast. **B.** Quantitative histopathological scoring of bone marrow and cortical bone based on leukemic infiltration, marrow cellularity, hematopoietic heterogeneity, and osteoclastic activity, demonstrating improved marrow integrity following α-Mercurin treatment to ALL animals. See Figure S19 for extended data. **C.** Representative DIC/confocal images of CD45⁺ and CD34⁺ cells from bone marrow smears. Healthy and α-Mercurin control animals depict preserved erythroid lineage and sparse CD45⁺CD34⁺ progenitors. In contrast, ALL control animals exhibit erythroid suppression and expansion of CD45⁺CD34⁺ leukemic blasts. Cytarabine treatment reduces blast burden but is associated with persistent erythroid suppression, whereas α-Mercurin treatment reduces CD45⁺CD34⁺ populations while restoring marrow heterogeneity and erythropoiesis. Scale bar: 40 µm. **D.** Representative H&E-stained thymus sections across experimental groups. ALL controls exhibit thymic atrophy with loss of corticomedullary distinction. Cytarabine treatment results in cortical lymphocyte depletion and structural disruption, whereas α-Mercurin treatment partially restores thymic architecture and cortical thickness. Scale bar: 100 µm. **E.** Quantitative histopathological scoring of thymus based on atrophy, corticomedullary distinction, and lymphoid depletion, demonstrating improved thymic integrity following α- Mercurin treatment. See Figure S20 for extended data. All histopathological scores and clinical biochemistry data are presented as median with IQR (25-75 percentile) in box plots; whiskers represent minimum-maximum values. Kruskal-Wallis test followed by Dunn’s multiple comparisons test was performed for statistical analysis. Statistical significance is denoted as *p ≤ 0.05, **p ≤ 0.01, ***p ≤ 0.001, ****p ≤ 0.0001.

We further evaluated CD45⁺CD34⁺ cells by immunofluorescence (Figure 6C). Healthy and α-Mercurin controls showed sparse CD45⁺CD34⁺ cells within a heterogeneous marrow background. In contrast, ALL controls showed dense CD45⁺CD34⁺ cell clusters with reduced representation of mature hematopoietic populations. Cytarabine treatment reduced these clusters but resulted in a sparsely populated marrow with limited recovery of erythroid cells. α-Mercurin treatment reduced CD45⁺CD34⁺ cell accumulation and indicated a restored heterogeneous cellular distribution, with reappearance of differentiated hematopoietic populations.

We also evaluated thymic histology to assess changes in lymphoid tissue following treatment (Figure 6D–6E and Figure S20). Healthy and α-Mercurin controls showed preserved thymic architecture with clear corticomedullary organization, dense cortical thymocytes, and an organized medulla. ALL controls showed marked thymic atrophy, cortical thinning, loss of corticomedullary demarcation, lymphoid depletion, and vascular congestion, with a total HS of 9.55 (IQR: 8.25–11.75; P=0.0027 vs healthy control). Cytarabine treatment reduced leukemic burden but showed further cortical depletion, apoptotic bodies, and poorly defined thymic architecture (total HS: 8, IQR: 6.25–10.5). In contrast, α-Mercurin treatment showed improved cortical thickness and re-established corticomedullary organization, with mild residual lymphoid depletion and a total HS of 7 (IQR: 5.25–7.75).

### 3.13. α-Mercurin shows predominant renal distribution with no detectable Hg in brain

We quantified tissue Hg levels using CV-AAS to evaluate the distribution of Hg in ALL animals following α-Mercurin treatment, terminally (Figure 7A). We observed a marked organ-specific distribution pattern, with the highest Hg level in the kidney (∼21,998 µg/kg), accounting for ∼94.81% of the total detected Hg across the analyzed tissues. Considering a combined kidney weight of ∼2 g, the measured renal Hg corresponded to ∼44 µg, which represented only ∼1.1% of the total administered Hg (∼4 mg). We detected lower Hg levels in the liver (∼643.3 µg/kg), spleen (∼435.5 µg/kg), lung (∼141.4 µg/kg), heart (∼50.76 µg/kg), and bone (∼43.46 µg/kg). Thymus and lymph nodes showed Hg levels of ∼243.4 µg/kg (∼0.36%) and ∼323.0 µg/kg (∼0.38%), respectively. Hg level in the brain remained below the detection limit. Therefore, we found no measurable brain Hg exposure under the conditions used in this study. We further compared tissue Hg levels between α-Mercurin-treated ALL animals and the corresponding controls to assess disease-associated Hg distribution (Figure 7B). We observed higher Hg levels in several tissues of α-Mercurin-treated ALL animals, including ∼9-fold higher level in liver (P=0.0485), ∼3-fold in kidney (P=0.004), ∼13-fold in spleen (P=0.0485), and ∼5-fold in bone, with similar trends in other tissues.

**Figure 7.**
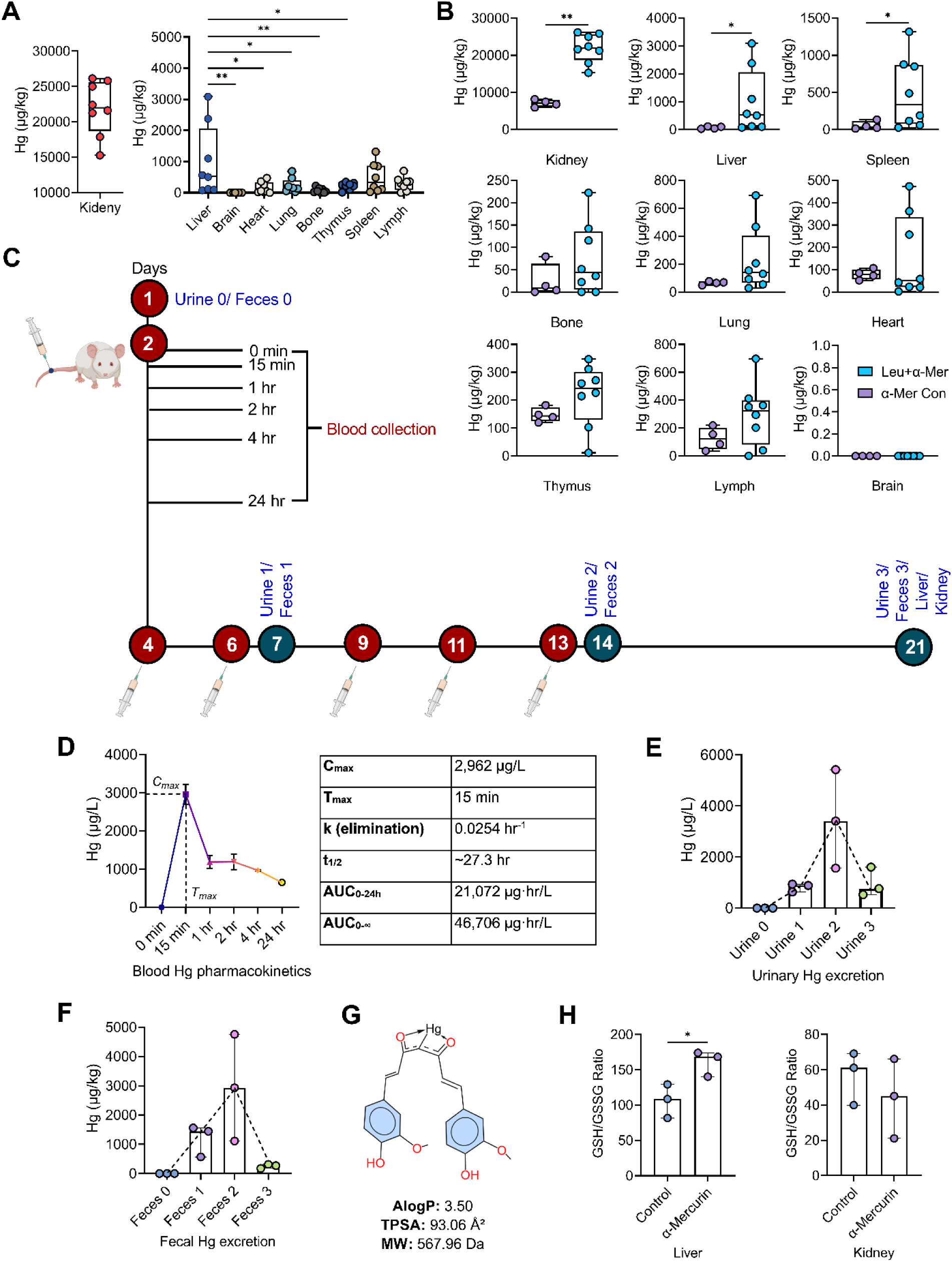
α-Mercurin demonstrates rapid tissue distribution and dual route (urine/feces) elimination of Hg with mild oxidative response. **A.** Graphical representation of total Hg content (µg/kg, wet weight) in each tissue as determined using CV-AAS, terminally. α-Mercurin-treated ALL animals demonstrates major Hg distribution in kidney, indicating renal clearance, while minute levels in liver and spleen, and trace level in thymus, lymph, lung, heart, bone marrow, with no detectable Hg in CNS. **B.** Comparative analysis of organ specific Hg biodistribution for α-Mercurin treated ALL and control animals, showing increased Hg level in ALL bearing animals. Statistical analysis was performed using Mann-Whitney U test. Statistical significance is denoted as *p ≤ 0.05, **p ≤ 0.01, ***p ≤ 0.001, ****p ≤ 0.0001. **C.** Schematic overview of the experimental design for blood Hg pharmacokinetics, urinary/fecal sample collection for Hg elimination study, and reduced glutathione to oxidized glutathione disulfide (GSH/GSSG) assay of liver and kidney, following α-Mercurin treatment. **D.** Blood Hg pharmacokentics was analyzed with CV-AAS from a single dose (1.6 mg/kg body weight) IV administration, followed by blood samples collection at 0 min, 15 min, 1 hr, 2 hr, 4 hr and 24 hr; demonstrating rapid tissue distribution including C_max_ of 2962 ug/L and T_max_ of 15 min with t_1/2_ of ∼27.3 hr. **E.** Urinary Hg elimination analysis via CV-AAS showing active renal elimination and progressive clearance. **F.** Fecal Hg elimination analysis with CV-AAS showing active clearance of administered Hg by fecal output. **G.** Physicochemical descriptors of neutral α-Mercurin including AlogP, topological polar surface area (TPSA) and molecular weight (MW), calculated by Epik engine of Schrödinger. **H.** The GSH/GSSG ratios of liver and kidney tissues following α-Mercurin treatment, suggesting that liver exhibited a significant adaptive response to the metabolic workload, while kidney indicates a mild oxidative stress. All data sets for pharmacokinetics, Hg elimination and GSH/GSSG ration are presented as median with IQR (25-75 percentile) in box plots; whiskers represent minimum-maximum values. For urine/fecal Hg elimination data, statistical analysis was performed by Kruskal-Wallis test followed by Dunn’s multiple comparisons test. For GSH/GSSG ratio, statistical analysis was performed by Welch’s t test. Statistical significance is denoted as *p ≤ 0.05, **p ≤ 0.01, ***p ≤ 0.001, ****p ≤ 0.0001.

### 3.14. α-Mercurin shows rapid systemic Hg exposure with urinary and fecal elimination

To understand the excretion mechanism of Hg, we further investigated blood Hg PK in combination with its urinary and fecal estimation (Figure 7C). Following single IV administration of α-Mercurin (1.6 mg/kg body weight), blood Hg concentration peaked with C_max_ ∼2,962 µg/L at T_max_ of 15 min, indicating rapid membrane partitioning and tissue distribution (Figure 7D). Blood Hg declined sharply by ∼60% within the first hr, followed by a slower terminal elimination phase with half-life (t_1/2_) of ∼27.3 hr (AUC_0–24h_: ∼21,072 µg·hr/L). Hg remained detectable at 24 hr post-injection, consistent with ongoing tissue-to-blood redistribution during the elimination phase.

We next evaluated urinary and fecal Hg excretion to further characterize Hg elimination profile (Figure 7E–7F). Urinary Hg elevated progressively with repeated dosing (1.6 mg/kg body weight, thrice/weekly), reaching ∼887 µg/L after one week of treatment and peaking at ∼3,407 µg/L after two weeks of treatment. Following cessation of treatment, urinary Hg declined sharply to ∼763 µg/L during the washout period, confirming active renal excretion and progressive clearance. Similarly, fecal Hg also increased progressively, reaching ∼1,445 µg/kg after one week and ∼2,933 µg/kg after two weeks. During washout, fecal Hg declined markedly ∼273 µg/kg, confirming near-complete clearance of fecal Hg output following treatment discontinuation. This observed PK profile is also consistent with in silico physicochemical descriptors of neutral α-Mercurin, calculated using Epik engine of Schrödinger (Figure 7G). The ALogP of 3.50 indicates moderate lipophilicity(66), while the topological polar surface area (TPSA) of 93.06 AÅ^2^ is greater than the established passive BBB permeation threshold of ∼90 AÅ^2^(67,68). The molecular weight (MW) of 567.96 Da, combined with ALogP also supports concurrent renal and hepatobiliary clearance(69,70).

### 3.15. α-Mercurin shows organ-specific GSH/GSSG ratios in liver and kidney

To evaluate the effect of α-Mercurin on cellular oxidative stress, we measured the ratios of reduced glutathione to oxidized glutathione disulfide (GSH/GSSG) in both liver and kidney tissues following the cessation of treatment (Figure 7H). Experimental results showed different pattern of organ-specific responses. α-Mercurin-treated animals showed a significant surge in its antioxidant pool, resulting to a hyper-reductive ratio of ∼168.5 comparing to untreated liver ∼108.6 (P=0.0386). Conversely, in kidney, α-Mercurin-treatment suggested a mild oxidative response, reducing the GSH/GSSG ratio to ∼45.08, comparing to healthy kidney of ∼61.13.

## 4. DISCUSSION

In the present study, we performed a comprehensive therapeutic evaluation of α-Mercurin, an organomercury derivative of curcumin, in an ENU-induced autochthonous ALL rat model. In our previous study, we reported the emergence of a novel organomercury compound, α- Mercurin to address the poor aqueous solubility and limited therapeutic applicability of curcumin. Similarly, therapeutic application of Hg is also restricted due to its potential systemic toxicity of previous formulations and route of administrations. In contrast, direct bonding of Hg at the α-carbon of curcumin (C_α_–Hg) produced a distinct organometallic framework. This modification allowed formulation of α- Mercurin as an aqueous-soluble sodium salt, suitable for IV administration. Our previous study also showed preferential leukemic cell killing through ROS-associated mitochondrial apoptosis, in vitro as well as ex vivo. In resonance, here, we further evaluated translational potential of α-Mercurin in vivo, in terms of overall therapeutic response, including its effect on hematopoietic and immune health, along with organ safety, systemic exposure, tissue Hg distribution, blood Hg pharmacokinetics (PK) and elimination profile.

α-Mercurin produced sustained disease control with improvement in general physiology. ALL-bearing animals progressively lost body weight and rapidly reached predefined humane endpoints, whereas α-Mercurin-treated ALL animals regained body weight during treatment. Cytarabine also improved body weight by delaying disease progression, in accordance with its established cytoreductive activity. Both treatments prolonged time-to-humane-endpoint, with median of 28.5 days for cytarabine and 39 days for α-Mercurin. We consider this finding as evidence of overall therapeutic efficacy under the present experimental conditions.

The hematological findings showed distinct pharmacodynamic (PD) profiles for the two treatment groups. Cytarabine produced a strong nonspecific cytoreductive response with marked reduction in circulating blasts (∼67%) and WBC counts from treatment initiation point. However, persistent erythroid suppression, incomplete platelet recovery, and relatively hypocellular marrow was also evident, in consistent with its known myelosuppressive effects. In comparison, α-Mercurin produced a gradual reduction in leukemic burden (∼25%) together with improvement in RBC, Hb, platelet, lymphocyte, and neutrophil populations from treatment initiation point. Bone marrow examination further showed reduced leukemic infiltration with scattered reappearance of erythroid precursors and megakaryocytes. Thus, the therapeutic response to α-Mercurin involved disease control together with improvement in hematopoietic compartments. We also observed a consistent pattern in immune profiling. ALL induction caused substantial immune dysregulation, including T-cell depletion, CD8⁺ predominance, depletion of CD19⁺ B cells, expansion of triple-negative populations, thymic atrophy, and disruption of lymphoid architecture. Both treatments showed improvement in immune health with different approach. α-Mercurin increased CD3⁺ and CD4⁺ populations, improving CD4⁺:CD8⁺ balance, also partially restored CD19⁺ B cells as well as CD4⁺CD8⁺ DP thymocytes, along with rejuvenated lymph node and thymic architecture. These findings suggest that α-Mercurin can support immune health significantly during leukemia control, without causing nonspecific suppression of normal immune compartments. Histopathology further supported the observed therapeutic and safety profile. α-Mercurin reduced leukemic infiltration in liver, kidney, spleen, lymph nodes, bone marrow, and thymus; thus, helped in preservation of tissue architecture. We also observed preserved lung, brain, and cardiac morphology. In contrast, cytarabine treatment caused neurohistological, myocardial, and myelosuppressive changes, along with its effective cytoreductive response. Overall serum biochemical findings are also consistent with tissue observations, demonstrating that α-Mercurin improved several hepatic parameters. Collectively, these findings suggest promising therapeutic profile of α-Mercurin.

Importantly, tissue biodistribution study of Hg via CV-AAS showed predominant renal Hg content, but the estimated renal Hg was only ∼1.1% of the cumulative administered Hg. Thus, high renal concentration did not represent extensive total renal retention and instead suggested active renal handling and clearance. We detected only trace levels of Hg in other tissues, while brain Hg remained below the detection limit, indicating no blood-brain-barrier (BBB) penetration. The calculated TPSA (∼93.06 Å²) provides supportive physicochemical explanation for restriction upon BBB penetration, although the CV-AAS finding remains the fundamental evidence. Importantly, Hg PK and biodistribution study via CV-AAS showed a biphasic profile for elimination from blood circulation with calculated terminal half-life of ∼27.3 hours. This finding is consistent with a two-compartment PK model in which the initial rapid decline reflects tissue distribution and the prolonged terminal phase reflects slow tissue-to-blood redistribution followed by elimination from the host system, indicating moderate lipophilicity. The MW of 567.96 Da further supports simultaneous renal and hepatobiliary metabolism, as compounds in this MW range are accessible to both glomerular filtration as well as biliary excretion. Significant availability of Hg in both urine and feces demonstrates both renal and hepatobiliary pathways are simultaneously playing role in elimination of Hg, during repeated IV dosing. Critically, both urinary and fecal Hg fell sharply during the one-week washout period, demonstrating that cessation of dosing leads to rapid reduction in active excretion, consistent with a clearance-dominant PK profile rather than progressive accumulation. This dual-route excretion pattern, in combination with minimal detection of Hg in renal tissue at terminal sacrifice suggest that the present formulation of organomercury therapeutic is suitable for active systemic clearance, without progressive organ accumulation. This pharmacokinetic safety feature of rapid tissue distribution, dual-route active elimination and BBB exclusion collectively distinguishes α- Mercurin from inorganic mercury species and other toxic forms of mercury, those are known for CNS accumulation and nephrotoxicity.

Further, the liver demonstrates remarkable physiological resilience to the metabolic workload for α-Mercurin treatment. Significant elevation of GSH/GSSG ratio demonstrates the hepatoprotective potential of our compound most likely due to antioxidant property of curcumin; thus, helped in conservation of histopathological features of the hepatic tissue. However, simultaneous elevations in serum urea and BUN reflect an initial reduction in renal filtration capacity. At the molecular level, the evidential drop in the Kidney GSH/GSSG ratio reveals that transient cellular oxidative stress may drives this functional switch. Importantly, our histopathological analysis confirms no structural distortion in renal tissue. Therefore, the stress remains limited within the functional level and could be managed via time dependent excretion and reabsorption inhibitors, if required. Taken together, the present data sets characterize the α- Mercurin as a promising, well tolerable anti-cancer compound that could be used alone or in combination with other therapeutic molecules.

We also acknowledge that the ENU-induced model provides an autochthonous disease environment with heterogeneous mutations and does not represent a defined molecular subtype of human ALL. Future studies should be conducted in genetically engineered and/or patient-derived animal models. In the next study molecular pharmacology should also be investigated including PK of α-Mercurin, along with metabolite profiling, PK/PD relationship and long-term toxicity.

## SUPPORTING INFORMATION

Descriptive histopathology of major organs and additional figures illustrating detailed findings of hematological profiling, immunophenotyping, and histopathological scoring are provided in the ‘Supporting Information’ document. All supporting information figures are denoted as “[Figure S(number)]” and tables are denoted as “[Table S(number)]”. Two representative videos from in vivo studies are also included as supporting information, demonstrating intravenous (IV) administration of α-Mercurin and cytarabine.

## AUTHOR CONTRIBUTIONS: CRediT

**Sougata Mondal:** Data curation; Formal analysis; Investigation; Software; Methodology; Validation; Visualization; Writing – original draft; Writing – review and editing. **Oyendrila Ghosh:** Data curation; Formal analysis; Methodology; Validation; Writing – review and editing. **Basab Bagchi:** Formal analysis; Methodology; Validation; Writing – review and editing. **Pratima Jana:** Formal analysis; Methodology; Validation; Writing – review and editing. **Bidisha Maiti:** Formal analysis; Methodology; Validation; Writing – review and editing. **Kalyan Kusum Mukherjee:** Conceptualization; Formal analysis; Funding acquisition; Resources; Validation; Writing – review and editing. **Supratim Ghosh:** Conceptualization; Data curation; Formal analysis; Funding acquisition; Investigation; Methodology; Project administration; Resources; Supervision; Validation; Visualization; Writing – original draft; Writing – review and editing

## DECLARATION OF GENERATIVE AI AND AI-ASSISTED TECHNOLOGIES IN MANUSCRIPT PREPARATION

During the preparation of this manuscript, the authors used Claude (Anthropic) and ChatGPT (OpenAI, USA) for assistance in manuscript drafting, structural editing, scientific writing refinement and language clarity improvement. After using these tools, the authors reviewed and edited the content as needed and take full responsibility for the content of the manuscript for publication.

## Supporting information

Supporting information, including additional datasets for better understanding

## ACKNOWLEDGEMENTS

Authors acknowledge funding support from the Indian Council of Medical Research (ICMR) and the Chittaranjan National Cancer Institute (CNCI). Authors also acknowledge the Centre for Laboratory Animal Research and Training (CLART), under West Bengal Livestock Development Corporation Ltd. for providing the Wistar rats for this investigation. Authors are also thankful to Dr. Upasana Das from Wake Forest University School of Medicine, USA, for conceptual contribution and Mr. Rupankar Ghosh and Mr. Subhajit Bhowmik from CNCI for experimental support.

## DATA AVAILABILITY STATEMENT

All data supporting the findings of this study are available within the article and its Supporting Information. Raw hematological, immunophenotyping, and histopathological scoring data are available from the corresponding author upon reasonable request. Representative in vivo videos demonstrating intravenous administration of α-Mercurin and cytarabine are included as Supporting Information.

