## Supporting information, including additional datasets for better understanding for "A first-in-class multimodal organomercury compound demonstrates preferential blast reduction with improved hematopoietic and immune health in ALL"

[1]Department of Anti-Cancer Drug Development and Chemotherapy, Chittaranjan National Cancer Institute, 37, S. P. Mukherjee Road, Bhowanipore, Kolkata 700026, West Bengal, India. [2]Department of Medical Oncology, Street Number 299, DJ Block, Action Area 1D, New Town, Kolkata 700160, West Bengal, India. [3]Department of Medical Oncology, Chittaranjan National Cancer Institute, 37, S. P. Mukherjee Road, Bhowanipore, Kolkata 700026, West Bengal, India.

***Corresponding author:** Supratim Ghosh. Department of Anti-Cancer Drug Development and Chemotherapy, Chittaranjan National Cancer Institute, 37, S. P. Mukherjee Road, Bhowanipore, Kolkata 700026, West Bengal, India.

#### S1. Detailed histopathological evaluation

##### S1.1. α-Mercurin demonstrates preserved brain architecture

Histopathological investigation in combination with total histopathological score (Figure 3G–3H and Figure S12) of the cerebellum and cerebrum sections revealed similar patterns in both healthy control as well as α-Mercurin control animals. Findings exhibited completely preserved cortical and cerebellar architecture with intact neuronal layers and normal white matter. In contrast, cerebrum sections from the ALL control animals were found with occasional neuronal shrinkage and scattered microglial activation, while cerebellar section showed mild edema in the molecular layer and minimal changes in the structure of Purkinje neurons. Notably, no leukemic infiltration was detected in either region of the brain of ALL control animals. Overall, histopathological architecture of both regions suggests no ENU-induced toxicity in CNS. On the contrary, cytarabine-treated ALL rats demonstrated the highest degree of neurotoxicity. The cerebrum showed clear neuronal degeneration, marked microgliosis and observable white matter demyelination. The cerebellum exhibited Purkinje cell degeneration/apoptosis, vacuolation of molecular layer and demyelination of white matter. These findings are well aligned with the known neurotoxic profile generated by cytarabine. Significantly, ALL animals treated with α-Mercurin did not produce any noticeable neurotoxic lesions in either brain region. Neuronal morphology remained intact with preserved white matter, cerebellar layers and Purkinje cells. No gliosis, vascular injury or demyelination was observed. The observations from cerebrum and cerebellum of α-Mercurin-treated ALL animals also showed strong contrast to that of cytarabine-treated ALL animals. Histopathological analysis suggested no toxic effect on CNS of α-Mercurin treated animals and the observation is consistent with the no detectable mercury accumulation in brain, as showed by CV-AAS.

##### S1.2. α-Mercurin suggests myocardial integrity

Alongside, total histopathological score and histopathological evaluation of H&E-stained cardiac sections was performed as shown in Figure 4A–4B and Figure S13. Animals in the healthy control group showed well-preserved myocardial architecture, associated with clear cross-striations of intact cardiomyocytes having uniform nuclear morphology. Similar myocardial morphology was detected for α-Mercurin control group; cardiac muscles remained well organized with no cardiomyocyte degeneration, necrosis, interstitial fibrosis or vascular lesions. In contrast, ALL control group exhibited mild pathological alterations, including mild edema in the myocardial region with scattered inflammatory cell congestion within intramyocardial vessels. Nevertheless, these findings are consistent with systemic leukemic stress, where cardiomyocyte nuclei remained mostly preserved with no necrosis or fibrosis. Conversely, cytarabine treatment demonstrated cardiotoxic changes. There was notable myocardial swelling along with myocyte necrosis. Some regions had disorganized myofibers, accompanied by dense inflammatory infiltrates with myocarditis and disruption of vascular integrity. However, ALL animals treated with α-Mercurin showed largely preserved myocardial structure, with intact cardiomyocyte morphology; although, occasional inflammatory cell infiltration was observed in the interstitial spaces due to persistence of residual disease. The infiltrates did not disrupt continuity of the fibers and was not associated with edema; suggesting leukemic stress. Overall, histopathological analysis suggested no significant effect on cardiac tissue due to α-Mercurin treatment, comparing to cytarabine treatment.

##### S1.3. α-Mercurin reduces hepatic leukemic infiltration while preserving liver ultrastructure

Figure 4C–4D and Figure S14 exhibit liver sections from healthy control group showing completely preserved hepatic architecture. The hepatic cords are well-organized, radiating from the central vein with regular sinusoidal spacing. Characteristic polygonal morphology of hepatocytes also retained with centrally placed nuclei and prominent nucleoli. Neither necrosis, inflammation or fibrosis were observed. Similarly, animals of α-Mercurin control group showed identical histoarchitecture as healthy controls with regular sinusoidal arrangement and hepatocyte morphology. No structural distortion and inflammatory infiltrates were observed, suggesting that α-Mercurin did not induce any intrinsic hepatotoxicity at the given dose. In contrast, ALL control group demonstrated substantial pathological alterations. High level of extramedullary hematopoiesis was observed, within sinusoidal spaces, resulting in degeneration of lobular architecture, along with Kupffer cell hypertrophy. In terms of treatment, cytarabine administration in ALL animals produced a notable reduction in leukemic blast infiltration. However, cytarabine treatment also caused pronounced hepatocellular toxicity, where hepatocytes exhibited significant hydropic degeneration with focal necrosis. Karyorrhexis along with apoptotic hepatocytes were also prominent. Furthermore, there was also clear loss of polygonal architecture of hepatocytes along with Kupffer cell hypertrophy and hyperplasia, suggesting chemotherapy-induced hepatic injury. Minimal steatosis was also detected in some cases. Interestingly, liver histology of ALL animals treated with α-Mercurin showed considerable therapeutic benefit with reduced leukemic blast burden and no additional structural injury. Unlike cytarabine, α-Mercurin restored normal hepatic organization and no steatosis or fibrosis were observed, apart from minimal inflammatory response along with Kupffer cell hypertrophy, suggesting anti-leukemic activity and favourable hepatic tolerability.

##### S1.4. α-Mercurin maintains renal architecture with minimal tubular stress

Figure 4F–4G and Figure S15 display histopathological assessment of kidney sections to understand renal toxicity profile due to disease induction and treatment with cytarabine or α-Mercurin. Kidney sections of healthy control animals exhibited normal renal architecture and intact glomeruli with open Bowman's space in the cortex region. In both proximal and distal tubules, regular luminal architectures were observed having no inflammatory cells or protein casts. Additionally, the medullary region demonstrated well-preserved collecting ducts with no cytoplasmic vacuolization. Similar morphological observations were also recorded from the renal sections of α-Mercurin control animals, suggesting absence of intrinsic nephrotoxicity at the given dose. However, ALL-bearing animals exhibited significant renal alterations. Interstitial leukemic cell infiltration was found higher in cortex than medulla, leading to compression of adjacent tubules. Furthermore, glomerular tuft was found with mild hypercellularity and swelling; capillaries were also found to be congested with heteromorphic lymphocytes and neutrophils, suggesting endocapillary glomerulopathy. Focal tubular necrosis with pyknotic and karyorrhectic nuclei was also recorded, indicating tubular injury. Observably, ALL animals treated with cytarabine showed marked reduction in infiltrating blasts, compared to untreated ALL animals. However, they also showed mild tubular injury with epithelial flattening, cytoplasmic vacuolization and tubular dilation, along with occasional apoptotic / necrotic bodies. Glomeruli also showed inflammatory characteristics and mild tubular cell loss as well as mild swelling. Overall, cytarabine treatment improved kidney architecture, relative to ALL controls. Similarly, IV infusion of α-Mercurin in ALL animals improved renal physiology. Proximal tubules revealed mild epithelial swelling and scattered apoptotic / necrotic cells were also observed. Further, inflammation of glomerular tufts was also detected, while Bowman's space found to be preserved, suggesting similar physiology as cytarabine-treated group.

##### S1.5. α-Mercurin demonstrates preserved lung morphology

Figure 5A–5B and Figure S16 represent histopathological assessment of lung tissue from different groups. Healthy control animals revealed normal pulmonary architecture characterized with well-defined alveoli and thin alveolar septa. No evidence of alveolar or interstitial inflammation or edema was found. Pulmonary vessels also showed normal congestion along with regular Type II pneumocytes; alongside, bronchioles exhibited intact ciliated epithelium with sparsely distributed goblet cells. Histological investigation of lung from α-Mercurin control animals were also indistinguishable from healthy controls, indicating no evidence of pulmonary toxicity following α-Mercurin administration. In contrast, ALL control animals showed mild but consistent disease-related pulmonary alterations. Focal alveolitis was detected with mild inflammatory infiltrates in interstitial and perivascular region causing vascular congestion. Consequently, Type II pneumocyte hyperplasia was detected along with mild alveolar edema and occasional leukemic cell infiltration within pulmonary vessels. Further, bronchiolar architecture was found to be preserved with regular goblet cell distribution, while mild peribronchiolar inflammatory cells were found. Conversely, lungs from cytarabine-treated ALL animals exhibited minimal pathological alterations. A significant reduction in inflammatory infiltrates with minimal interstitial inflammation was found compared to ALL controls; though, occasional mild alveolar edema and rare Type II pneumocyte hyperplasia was observed. However, no hemorrhage and epithelial damage was found; accordingly, bronchiolar structure remained intact without any goblet cell hyperplasia. Similar to cytarabine, α-Mercurin treated ALL animals demonstrated preserved lung architecture, where alveolar structure was found to be intact with no inflammation or epithelial injury. However, very rare alveolar edema and occasional Type II pneumocyte hyperplasia were found. Bronchiolar epithelium and goblet cell distribution were found similar to healthy and α-Mercurin controls, suggesting pulmonary safety of α-Mercurin at given dose.

##### S1.6. α-Mercurin alleviates splenic pathology with reduction of leukemic infiltration

Histopathological investigation of spleen sections from different groups was performed and displayed in Figure 5C–5D and Figure S17. Datasets revealed distinct architectural and cellular alterations. Spleen histopathology from healthy control animals exhibited well-defined white pulps with intact periarteriolar lymphoid sheaths (PALS) surrounded by clear marginal zones (MZ). Normal sinusoidal organization with scattered megakaryocytes were also detected in the red pulp area, containing no inflammatory infiltrates and congestion or necrosis. Similarly, α-Mercurin control animals demonstrated normal splenic architecture as healthy controls with no histological signs of tissue injury and necrosis, suggesting absence of splenic toxicity at the given dose. In contrast, ALL control group demonstrated marked pathological alterations characterized by splenomegaly. Further, diffuse infiltration of leukemic blasts created atropic white pulp, damaging follicular architecture and reducing germinal centres significantly. Additionally, red pulp sinusoids were severely congested with large amount of extramedullary hematopoiesis, including occasional hemosiderin deposition. The collective evidences suggest high systemic leukemic burden and role of spleen as a secondary leukemic reservoir. Notably, cytarabine-treated

ALL animals exhibited a significant reduction in leukemic blast infiltration. Pathological evaluation suggested partial restoration of red pulp and white pulp architecture with visible margin. However, the white pulp showed poorly organized and depleted PALS with reduced germinal centres. Mild sinusoidal congestion was also detected along with increased apoptotic bodies and vacuoles, indicating reduced extramedullary hematopoiesis. Overall findings suggest effective decrease of disease burden with partial restoration of splenic architecture. Similarly, IV administration of α-Mercurin in ALL animals generated significant improvement. Accordingly, leukemic infiltration was largely reduced, accompanied by partial restoration of white pulp architecture and lymphoid follicles, in combination with decreased red pulp expansion and sinusoidal congestion. Further, distinct apoptotic or necrotic bodies and vacuoles with mild megakaryocytes were detected in red pulp, suggesting selective disappearance of dysfunctional extramedullary hematopoietic regions. Overall assessment depicted improved histopathological features of splenic structure for α-Mercurin treatment, comparing to cytarabine treatment, findings are in agreement with splenic immune profiling.

**S1.7. α-Mercurin preserves lymph node architecture**

Furthermore, histopathological evaluation of lymph nodes was performed to investigate disease progression and treatment-associated changes, results are shown in Figure 5E–5F and Figure S18. Lymph nodes from healthy control group exhibited intact architecture with well-defined capsule and clear demarcation between cortex, paracortex and medulla, where normal sized follicles are observed with occasional germinal centers (GC) and well-preserved mantle zones (MZ). Thin sinuses having minimal cellularity were seen with no atypical cells or necrosis. Correspondingly, α-Mercurin treated control animals showed preservation of normal lymph node architecture similar to healthy controls, suggesting that α-Mercurin alone does not exert any lymphotoxic effects at the given dose. In contrast, ALL control animals demonstrated significant pathological alterations, with diffused leukemic blast infiltration and high architectural distortion. Normal follicular structures were lost, along with expansion of medullary region. Leukemic blasts showed high nuclear-to-cytoplasmic ratio, condensed nucleoli as well as frequent mitotic figures; said observations are consistent with extensive leukemic involvement and disease progression. On the other hand, partial restoration of lymph node architecture was observed in cytarabine treated ALL animals. However, observed lymph nodes were markedly hypocellular, with significant lymphoid depletion and architectural distortion, including apoptotic bodies and cytoplasmic vacuoles. Atrophic follicles were also seen with partial reappearance of MZ and suppression of GC. Collective observations demonstrate cytotoxic elimination of leukemic blasts along with non-selective suppression of normal lymphoid populations. Similarly, lymph nodes from α-Mercurin treated ALL animals also exhibited partial restoration of overall architecture and moderate reduction in leukemic infiltration. Interestingly, follicles and GC were found to be reappeared along with an increased presence of morphologically matured lymphoid cells. Few apoptotic cells with cytoplasmic vacuolation were also found. Importantly, no significant lymphoid depletion was evident. Collective findings indicate anti-leukemic potential of α-Mercurin with potential for preservation of lymph node structure, suggesting physiological lymphoid homeostasis.

**S1.8. α-Mercurin restores bone marrow architecture**

Bone marrow (BM) and corresponding bone sections were examined to estimate leukemia burden, bone remodeling and treatment associated changes in the process of hematopoiesis; corresponding results are represented in Figure 6A–6B and Figure S19. BM sections from healthy control rats demonstrated normocellular marrow with a heterogeneous hematopoietic population. Myeloid (M), erythroid (E) and lymphoid precursors were also found at various stages of maturation with a normal M:E ratio. Histology of cortical bone suggested normal thickness with balanced osteoblast and osteoclast activity. Animals treated with α-Mercurin alone showed similar BM architecture like healthy controls with balanced osteoblast-osteoclast activity, suggesting no structural or functional disruption following given α-Mercurin dose. In contrast, BM sections from ALL control animals displayed marked hypercellularity with extensive infiltration of leukemic cells. The marrow was found predominantly occupied with blasts characterized by a high nuclear:cytoplasmic ratio, condensed chromatin and prominent nucleoli. Pronounced reduction in erythroid and megakaryocytic lineages was also observed, suggesting severe alterations in the process of hematopoiesis. Corresponding bone sections revealed irregular bone structures with increased osteoclastic activity, indicating ALL-associated disruption of bone architecture. Observably, BM sections from cytarabine treated ALL animals demonstrated significant reduction in leukemic blasts, which was accompanied by overall marrow hypocellularity, consistent with drug-induced myelosuppression and reduced mitotic figures. Bone histology showed notable reduction in osteoclastic activity with cytarabine treatment, though the level remains higher compared to healthy controls, indicating partial recovery of ALL-associated bone damage. Significantly, treatment of ALL animals with α-Mercurin resulted to a moderate reduction in leukemic blast density within BM, along with a partial restoration of heterogeneous hematopoietic population with reappearance of erythroid and myeloid precursors. Bone sections demonstrated reduced osteoclastic activity, depicting a more regular remodeling pattern compared to ALL controls, suggesting therapeutic recovery of ALL-induced bone pathology. Improvement in BM interface with homeostatic trend was evident in case of α-Mercurin treatment. Overall histopathological findings of BM demonstrated treatment-dependent reduction of leukemic burden and hematopoietic recovery; the same is reflected in terminal PB hematological profile and marrow cytomorphology.

**S1.9. α-Mercurin improves thymic integrity**

Moving towards histopathological examination of another primary lymphoid organ, group-dependent changes in thymic architecture was observed as shown in Figure 6D–6E and Figure S20. Thymus section of the healthy control group suggested well-defined architecture with clear distinction between cortex and medulla. Further, dense populations of thymocytes in cortex region were detected with no evidence of congestion, necrosis or inflammatory infiltration. Likewise, α-Mercurin control group demonstrated preserved thymic architecture and cell population, suggesting no adverse effect on thymic structure and function at the given dose. In contrast, ALL control animals showed observable thymic atrophy in combination with severe cortical lymphoid depletion and poor cortex-medulla distinction; alongside, vascular congestion and occasional inflammatory infiltrates were found, demonstrating ALL-dependent thymic impairment. Pronounced thymic alterations were also found in ALL animals treated with cytarabine, where thymus was severely reduced in size with apoptotic bodies and depleted cortical lymphocytes. Furthermore, the corticomedullary junction could not be clearly demarcated, while vascular congestion was also noted, suggesting chemotherapy-associated thymic suppression. In contrast, thymic sections from the ALL animals treated with α-Mercurin demonstrated partial restoration of thymic architecture compared to ALL control group, with an improved cortical thickness. However, moderate cortical lymphoid depletion persisted including mild residual vascular congestion, suggesting recovery of ALL-associated thymic damage, as also observed in immune-phenotyping by enhancement of DP thymocytes.

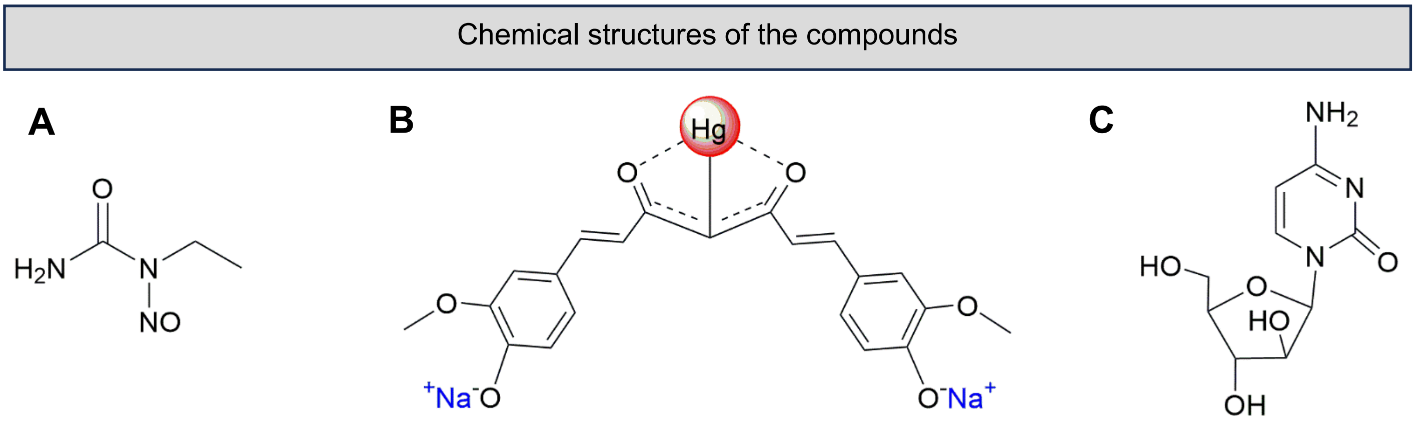

**Figure S1:** Chemical structures of the compounds used in this study. **A.** N-Nitroso-N-ethylurea (ENU), used for leukemia induction. **B.** α-Mercurin, organomercury derivative of curcumin; synthesized in our laboratory. **C.** Cytarabine (cytosine arabinoside; Ara-C), a clinically used standard chemotherapeutic drug for leukemia.

Synthesis scheme of α-Mercurin

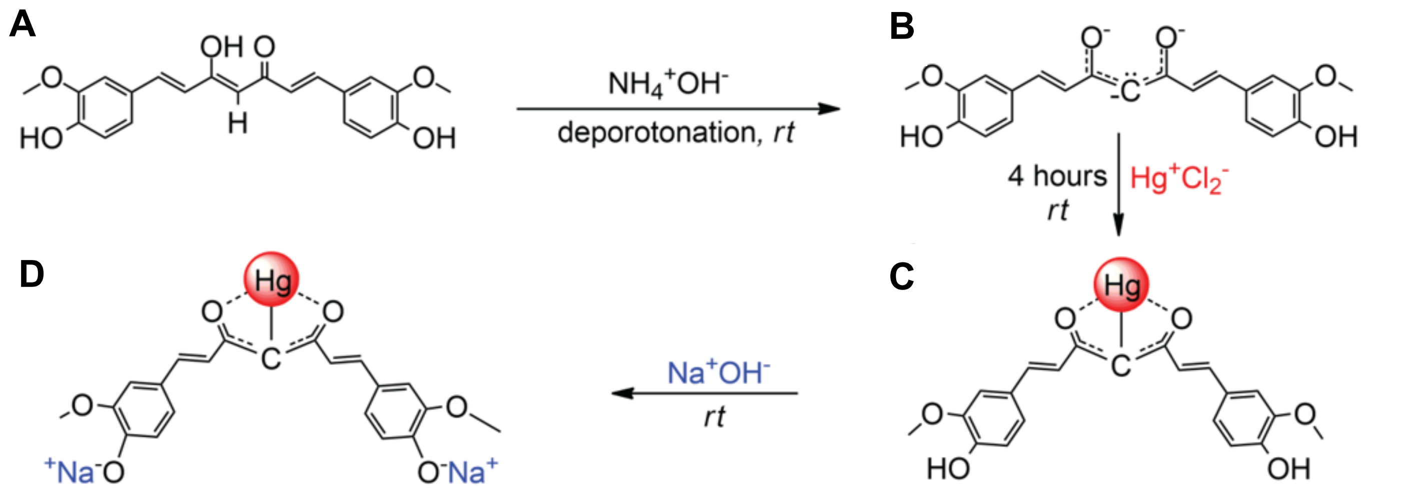

**Figure S2:** Reaction scheme for the synthesis of α-Mercurin. **A.** curcumin, **B.** curcumin in carbanion form (Intermediate), **C.** α-Mercurin, **D.** α-Mercurin in aqueous NaOH(1).

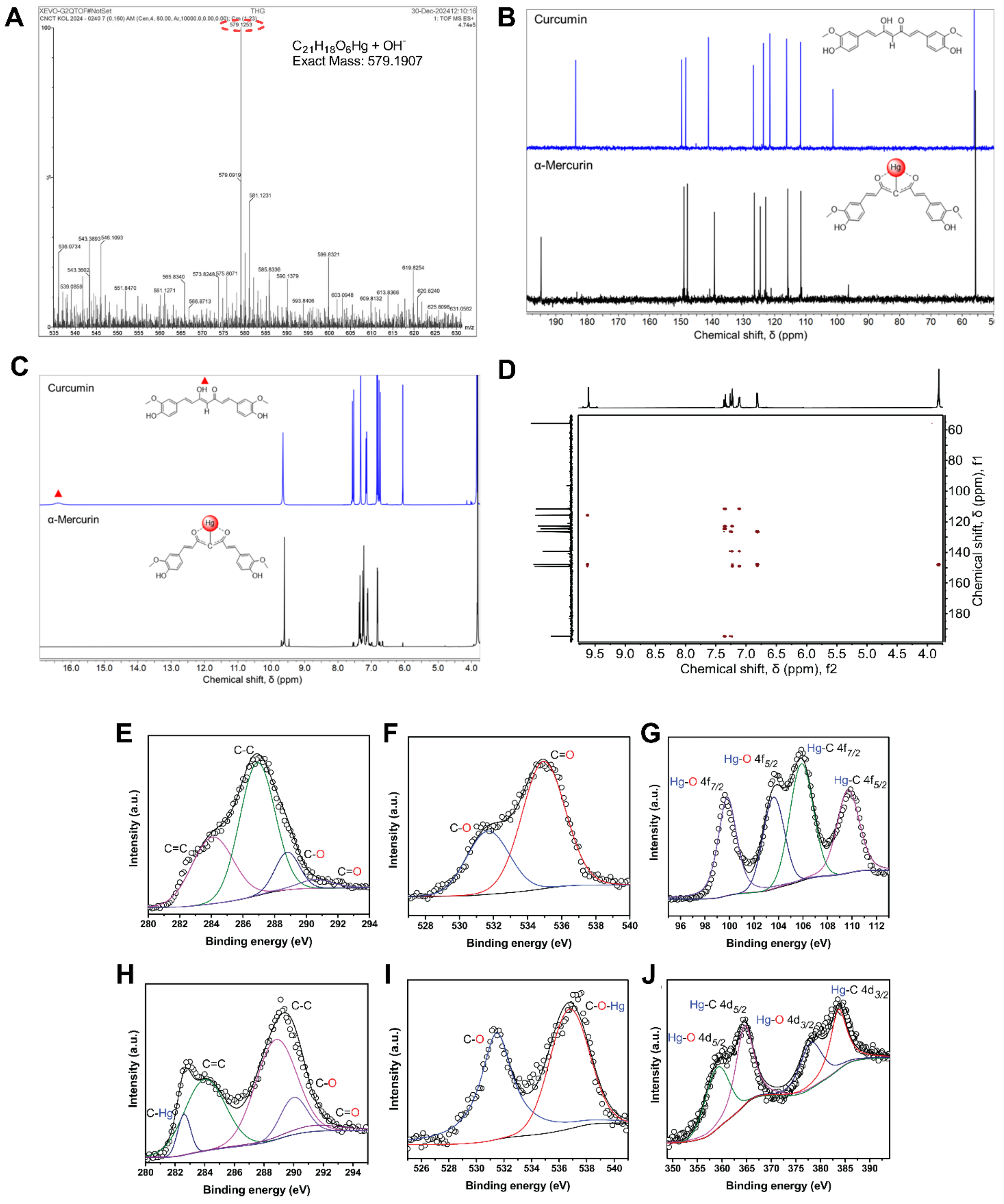

**Figure S3: A.** HR-MS spectrum of α-Mercurin. **B.** $^{13}C$ NMR spectra of α-Mercurin, comparing to curcumin. **C.** $^{1}H$ NMR spectra of α-Mercurin, comparing to curcumin. **D.** $^{13}C$-$^{1}H$ (f1/f2) HMBC NMR spectra of α-Mercurin. **E-F.** Deconvoluted XPS spectra of C1s and O1s core-level electrons of curcumin. **G-J.** Deconvoluted XPS spectra of Hg 4f, C 1s, O 1s, and Hg 4d core-level electrons of α-Mercurin; collectively demonstrating successful bonding of mercury atom to the α-carbon of curcumin(1).

Extended data of longitudinal hematological profile and terminal bone marrow smear

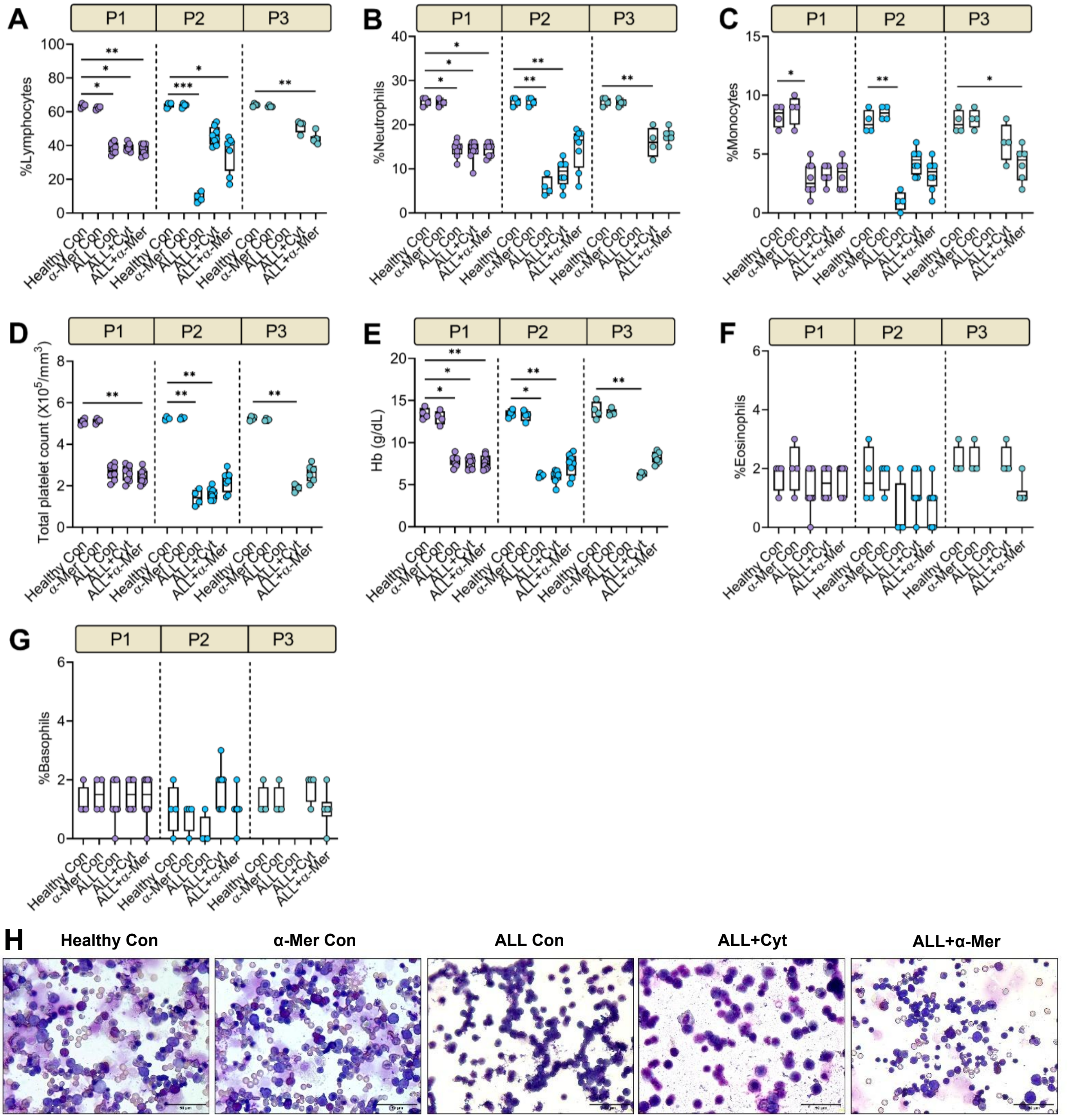

**Figure S4:** Graphical representation of **A.** %Lymphocytes, **B.** %Neutrophils, **C.** Monocytes, **D.** Total platelets, **E.** hemoglobin (Hb) concentration, **F.** %Eosinophils, **G.** %Basophils at Pull 1 (day 1, just before treatment), Pull 2 (day 14, post-treatment) and Pull 3 (day 28, post-treatment), across all groups; demonstrating disease progression and effect of treatment. Untreated ALL animals progressed rapidly and did not survive to Pull 3. P1: Pull 1, P2: Pull 2, P3: Pull 3. Data are presented as median with IQR (25 - 75 percentile) in box plots; whiskers represent minimum-maximum values. Kruskal-Wallis test followed by Dunn's multiple comparisons test was performed for statistical analysis. Sample sizes (n) varied across timepoints (Pull 1: n = 4-8; Pull 2: n = 4-8; Pull 3: n = 0-6) due to survival-associated attrition. Statistical significance is denoted as *p ≤ 0.05, **p ≤ 0.01, ***p ≤ 0.001, ****p ≤ 0.0001. **H.** Magnified view of Leishman-stained bone marrow smear. Scale bar: 50 μm.

#### Median %cell populations of longitudinal hematological profiling (Pull 1, 2, 3) till euthanasia (terminal)

| | Pull | Healthy Con | α-Mer Con | ALL Con | ALL+Cyt | ALL+α-Mer |
|---|---|---|---|---|---|---|
| Total WBC (X10³/mm³) | 1 | 4.70 | 4.90 | 30.70 | 30.20 | 32.80 |
| | 2 | 4.60 | 4.70 | 58.00 | 14.10 | 25.40 |
| | 3 | 4.80 | 4.80 | - | 8.40 | 19.10 |
| | Terminal | 4.90 | 4.90 | 59.20 | 11.00 | 16.50 |
| %Blasts | 1 | 0 | 0 | 41.5 | 41 | 42 |
| | 2 | 0 | 0 | 85 | 28 | 36.5 |
| | 3 | 0 | 0 | - | 17.5 | 31.5 |
| | Terminal | 0 | 0 | 85.5 | 13.5 | 30.5 |
| %Lymphocytes | 1 | 63.5 | 63 | 38.5 | 39 | 38.5 |
| | 2 | 64.5 | 64 | 9 | 45 | 41.5 |
| | 3 | 64 | 63 | - | 53 | 43.5 |
| | Terminal | 64 | 64 | 9 | 51.5 | 44 |
| %Neutrophil | 1 | 25.5 | 25 | 15 | 14.5 | 15 |
| | 2 | 25.5 | 25.5 | 5.5 | 9.5 | 16.5 |
| | 3 | 25.5 | 25 | - | 16 | 17.5 |
| | Terminal | 25 | 25 | 4 | 15 | 17 |
| %Eosinophil | 1 | 2 | 2 | 1 | 1.5 | 2 |
| | 2 | 1.5 | 2 | - | 1 | 1 |
| | 3 | 1.5 | 2 | | 2 | 1 |
| | Terminal | 1.5 | 2 | 0 | 1 | 1 |
| %Basophil | 1 | 1 | 1.5 | 1 | 1.5 | 1.5 |
| | 2 | 1 | 1 | 0 | 2 | 1 |
| | 3 | 1 | 1 | - | 2 | 1 |
| | Terminal | 1 | 0.5 | 0 | 1 | 1 |
| %Monocytes | 1 | 8.5 | 9 | 2.5 | 3 | 3.5 |
| | 2 | 7.5 | 8.5 | 1 | 4.5 | 3.5 |
| | 3 | 7.5 | 8 | - | 6 | 4.5 |
| | Terminal | 8 | 9 | 1 | 5 | 5 |
| Platelets count (X10⁵/mm³) | 1 | 5.04 | 5.14 | 2.72 | 2.65 | 2.34 |
| | 2 | 5.23 | 5.26 | 1.43 | 1.61 | 2.36 |
| | 3 | 5.25 | 5.16 | - | 1.95 | 2.65 |
| | Terminal | 5.30 | 5.23 | 0.88 | 1.97 | 2.60 |
| RBC count (X10⁶/mm³) | 1 | 7.525 | 7.585 | 5.155 | 5.11 | 5.215 |
| | 2 | 7.645 | 7.43 | 3.81 | 4.065 | 5.31 |
| | 3 | 7.765 | 7.6 | - | 4.35 | 5.465 |
| | Terminal | 7.905 | 7.71 | 3.21 | 4.45 | 5.595 |
| Hb conc. (g/dL) | 1 | 13.335 | 12.985 | 8.05 | 7.81 | 7.485 |
| | 2 | 13.375 | 13.285 | 6.055 | 6.21 | 7.6 |
| | 3 | 13.73 | 13.405 | - | 6.3 | 8.185 |
| | Terminal | 13.865 | 13.74 | 5.16 | 5.925 | 8.005 |

**Table S1:** Calculated median values of %blasts, total WBC, differential WBC, total RBC, platelet count and hemoglobin (Hb) concertation as found from longitudinal hematological profiling of peripheral blood smear at Pull 1 (day 1, just before treatment), Pull 2 (day 14, post-treatment) and Pull 3 (day 28, post-treatment), and terminally before euthanasia across all groups.

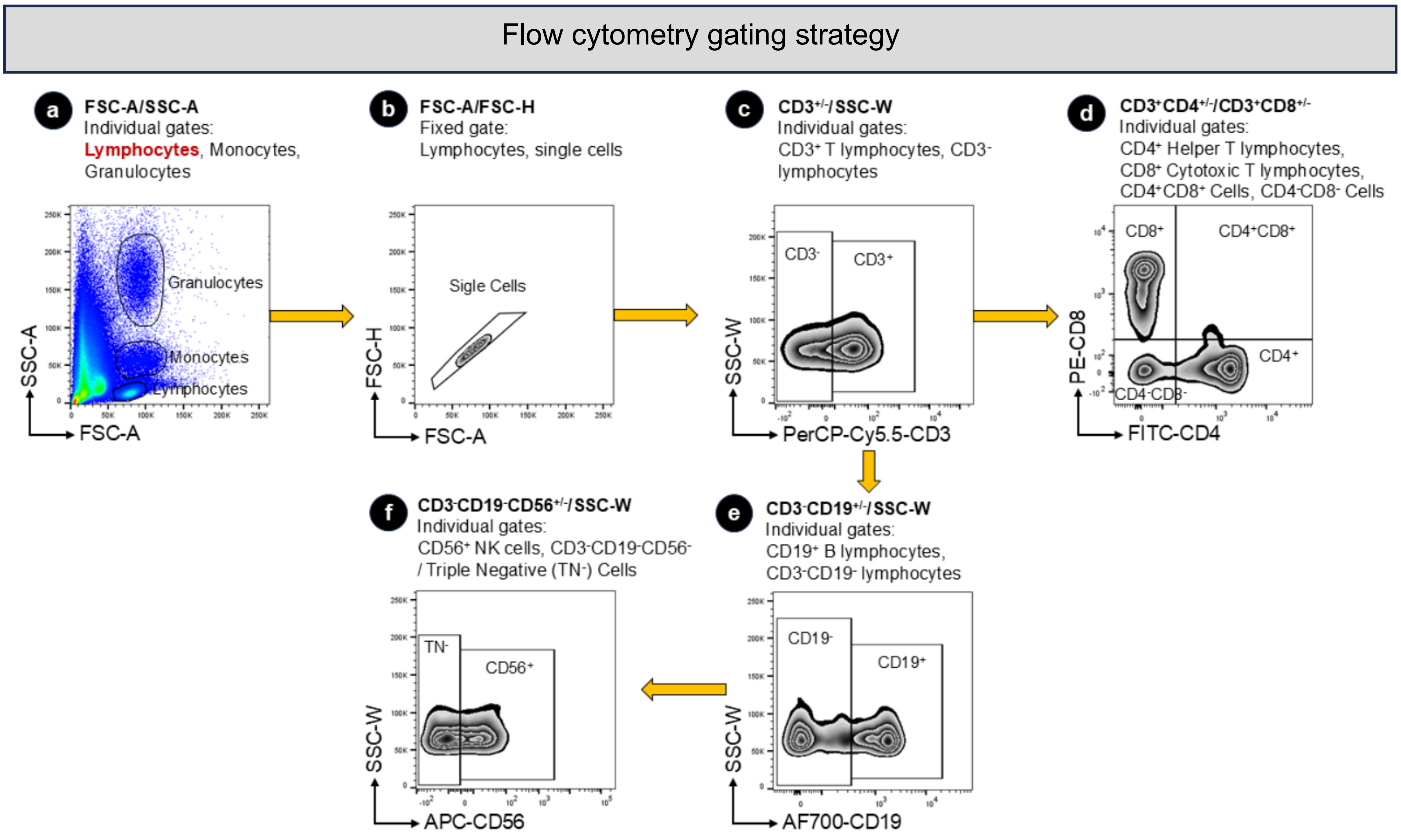

**Figure S5:** Representation of flow cytometry gating strategy. Gated lymphocyte compartment was further refined (FSC-A/FSC-H) to get single cells and then gated for CD3$^{+}$ T cells (PerCP-Cy5.5 positive) and CD3$^{-}$ cells. Following that, CD3$^{+}$ T cells were gated for CD4$^{+}$ (FITC positive) and CD8$^{+}$ (PE positive) cells. CD3$^{-}$ cells were differentiated to CD19$^{+}$ (A700 positive) and CD19$^{-}$ cells, followed by gating on CD19$^{-}$ cells to get CD56$^{+}$ (APC positive) and triple-negative (TN$^{-}$) cells. Absolute counts of CD3$^{+}$, CD19$^{+}$ and CD56$^{+}$ cells were presented as percent (%) of total lymphocytes, while CD4$^{+}$, CD8$^{+}$ and CD4$^{+}$CD8$^{+}$ cell counts are given as percent (%) of total CD3$^{+}$ T cell count.

Extended data of longitudinal immunophenotypic analysis of peripheral blood

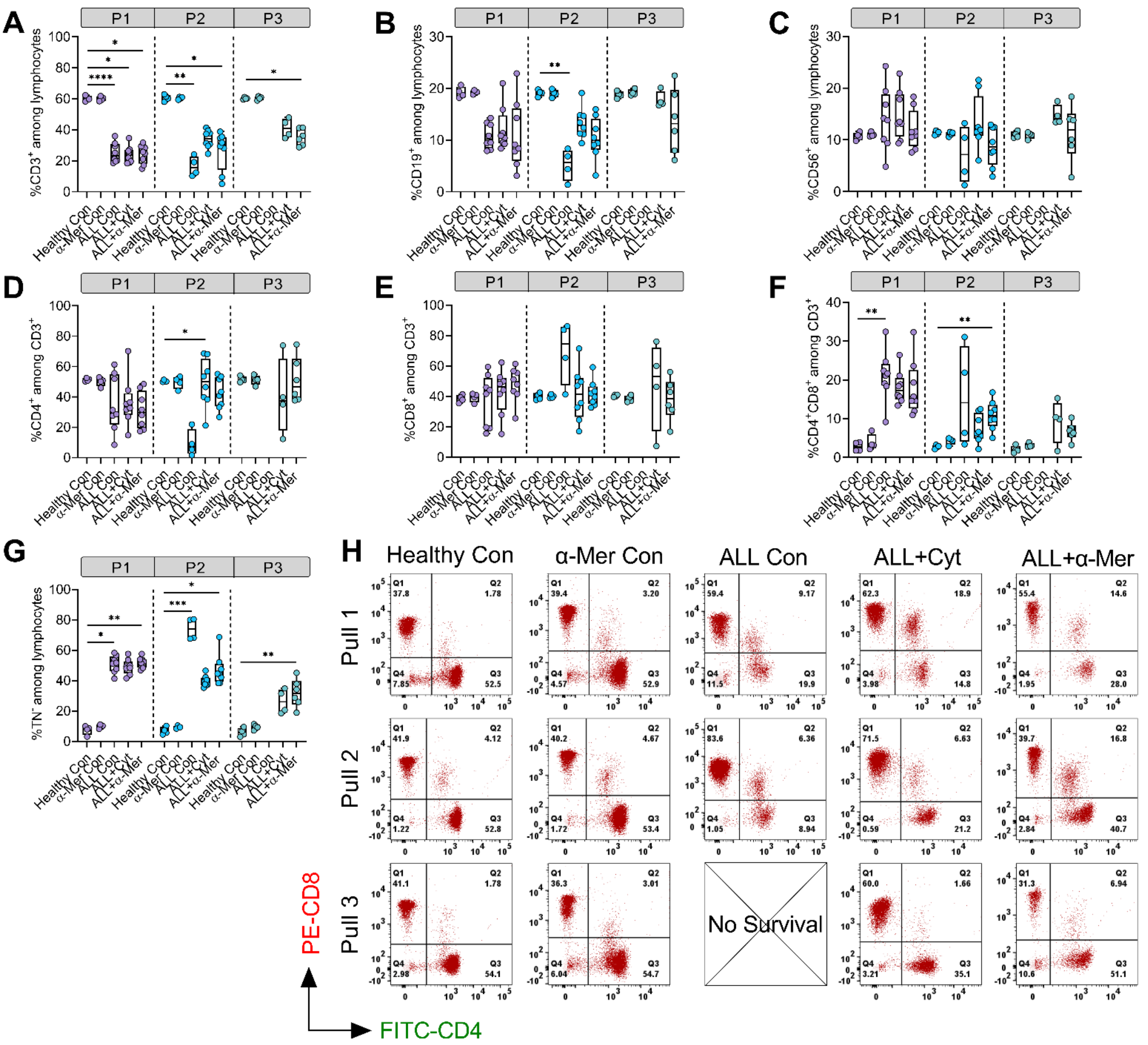

**Figure S6:** Panels showing graphical representation for **A.** %CD3+, **B.** %CD19+, **C.** %CD56+, **D.** %CD3+CD4+, **E.** %CD3+CD8+, **F.** %CD3+CD4+CD8+ and **G.** %TN- (triple-negative) cells at Pull 1 (day 1, just before treatment), Pull 2 (day 14 post-treatment) and Pull 3 (day 28 post-treatment), across all groups. Untreated ALL animals progressed rapidly and did not survive to Pull 3. P1: Pull 1, P2: Pull 2, P3: Pull 3. Data are presented as median with IQR (25 - 75 percentile) in box plots; whiskers represent minimum-maximum values. Kruskal-Wallis test followed by Dunn's multiple comparisons test was performed for statistical analysis. Sample sizes (n) varied across timepoints (Pull 1: n = 4-8; Pull 2: n = 4-8; Pull 3: n = 0-6) due to survival-associated attrition. Statistical significance is denoted as *p ≤ 0.05, **p ≤ 0.01, ***p ≤ 0.001, ****p ≤ 0.0001. **H.** Representative flow cytogram of CD3+ T cell subsets (%CD4+, %CD8+, %CD4+CD8+ and %CD4-CD8-) of peripheral blood at Pull 1 (day 1, just before treatment), Pull 2 (day 14, post-treatment) and Pull 3 (day 28, post-treatment), across all groups.

#### Median %cell populations with immune index, of peripheral blood at Pull 1, 2 and 3

| | | | | | | |
|---|---|---|---|---|---|---|
| $CD3^+$ | Pull | Healthy Con | α-Mer Con | ALL Con | ALL+Cyt | ALL+α-Mer |
| | 1 | 60.30 | 59.91 | 23.15 | 23.55 | 25.19 |
| | 2 | 60.85 | 60.62 | 15.75 | 34.25 | 30.50 |
| | 3 | 60.20 | 60.53 | - | 40.86 | 35.05 |
| $CD19^+$ | Pull | Healthy Con | α-Mer Con | ALL Con | ALL+Cyt | ALL+α-Mer |
| | 1 | 19.05 | 19.23 | 10.20 | 10.63 | 8.76 |
| | 2 | 19.19 | 19.04 | 5.63 | 12.90 | 9.96 |
| | 3 | 18.76 | 19.02 | - | 17.19 | 13.19 |
| $CD56^+$ | Pull | Healthy Con | α-Mer Con | ALL Con | ALL+Cyt | ALL+α-Mer |
| | 1 | 10.70 | 10.85 | 14.00 | 13.32 | 11.22 |
| | 2 | 11.30 | 11.25 | 7.15 | 11.79 | 8.68 |
| | 3 | 10.85 | 11.10 | - | 14.10 | 11.94 |
| $TN^-$ | Pull | Healthy Con | α-Mer Con | ALL Con | ALL+Cyt | ALL+α-Mer |
| | 1 | 9.85 | 9.86 | 51.51 | 48.87 | 51.95 |
| | 2 | 9.43 | 9.33 | 74.08 | 38.54 | 44.80 |
| | 3 | 10.20 | 9.75 | - | 26.05 | 31.45 |
| $CD4^+$ | Pull | Healthy Con | α-Mer Con | ALL Con | ALL+Cyt | ALL+α-Mer |
| | 1 | 51.17 | 50.48 | 29.96 | 34.10 | 30.65 |
| | 2 | 50.15 | 50.35 | 7.14 | 50.00 | 41.79 |
| | 3 | 51.12 | 51.27 | - | 40.65 | 46.60 |
| $CD8^+$ | Pull | Healthy Con | α-Mer Con | ALL Con | ALL+Cyt | ALL+α-Mer |
| | 1 | 38.58 | 40.17 | 42.95 | 46.26 | 51.23 |
| | 2 | 39.64 | 39.69 | 74.53 | 41.35 | 41.00 |
| | 3 | 40.61 | 38.06 | 0.00 | 48.75 | 43.11 |
| $CD4^+CD8^+$ | Pull | Healthy Con | α-Mer Con | ALL Con | ALL+Cyt | ALL+α-Mer |
| | 1 | 2.78 | 3.25 | 21.40 | 15.90 | 15.17 |
| | 2 | 2.79 | 4.48 | 14.14 | 5.89 | 10.72 |
| | 3 | 1.86 | 2.96 | - | 10.16 | 6.08 |
| $CD4^-CD8^-$ | Pull | Healthy Con | α-Mer Con | ALL Con | ALL+Cyt | ALL+α-Mer |
| | 1 | 8.26 | 6.95 | 5.46 | 1.76 | 2.12 |
| | 2 | 6.53 | 7.01 | 3.47 | 2.10 | 4.31 |
| | 3 | 5.62 | 7.35 | - | 2.28 | 2.74 |
| $CD4^+$:$CD8^+$ | Pull | Healthy Con | α-Mer Con | ALL Con | ALL+Cyt | ALL+α-Mer |
| | 1 | 1.33 | 1.28 | 0.70 | 0.75 | 0.60 |
| | 2 | 1.28 | 1.29 | 0.08 | 1.26 | 1.02 |
| | 3 | 1.27 | 1.35 | - | 0.84 | 1.04 |
| T:B | Pull | Healthy Con | α-Mer Con | ALL Con | ALL+Cyt | ALL+α-Mer |
| | 1 | 3.21 | 3.16 | 2.04 | 2.04 | 3.30 |
| | 2 | 3.21 | 3.21 | 3.17 | 2.85 | 2.87 |
| | 3 | 3.20 | 3.15 | - | 2.30 | 3.10 |
| Immune index | Pull | Healthy Con | α-Mer Con | ALL Con | ALL+Cyt | ALL+α-Mer |
| | 1 | 9.20 | 9.15 | 0.95 | 1.05 | 0.93 |
| | 2 | 9.61 | 9.71 | 0.26 | 1.60 | 1.23 |
| | 3 | 8.90 | 9.29 | - | 3.01 | 2.20 |

**Table S2:** Calculated median values of %immune cell populations along with immune cell ratios and immune index as found from longitudinal flow cytometric analysis of peripheral blood at Pull 1 (day 1, just before treatment), Pull 2 (day 14, post-treatment) and Pull 3 (day 28, post-treatment), across all groups.

### Extended data of terminal immunophenotypic analysis of peripheral blood

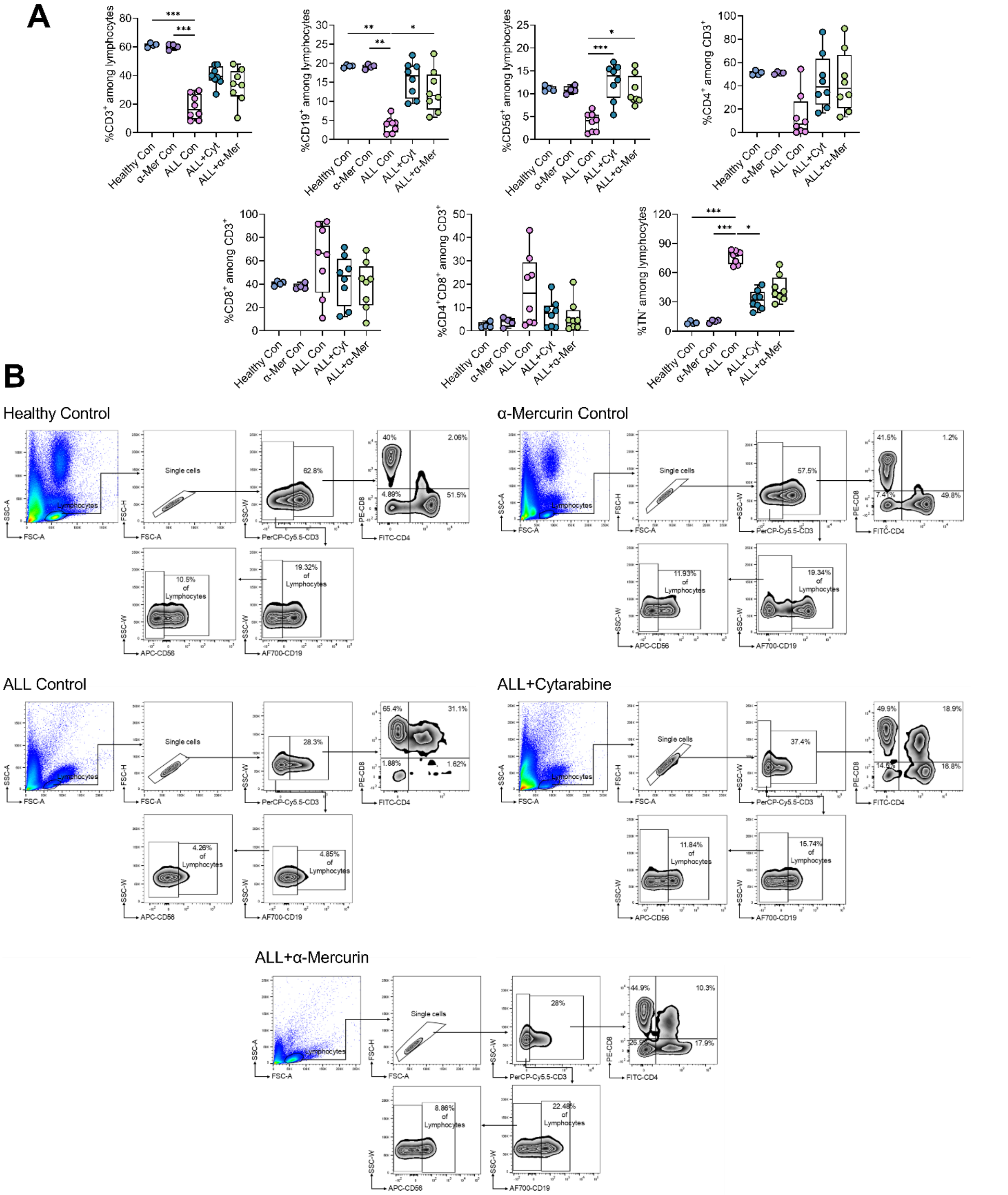

**Figure S7: A.** Graphical representation of percentages of cell populations (%$CD3^+$, %$CD19^+$, $CD56^+$, %$CD4^+$, %$CD8^+$, %$CD4^+CD8^+$ and %$TN^-$) investigated by flow cytometric analysis of lymphoid compartments from terminally collected peripheral blood samples. Data are presented as median with IQR (25 - 75 percentile) in box plots; whiskers represent minimum-maximum values. Kruskal-Wallis test followed by Dunn's multiple comparisons test was performed for statistical analysis. Statistical significance is denoted as *$p \leq 0.05$, **$p \leq 0.01$, ***$p \leq 0.001$, ****$p \leq 0.0001$. **B.** Representative flow cytograms across all groups demonstrating %$CD3^+$, %$CD3^+CD4^+$, %$CD3^+CD8^+$, %$CD3^+CD4^+CD8^+$ and %$CD3^+CD4^-CD8^-$, %$CD19^+$ and %$CD56^+$ cells of terminally collected peripheral blood samples.

#### Extended data of terminal immunophenotypic analysis bone marrow aspirate

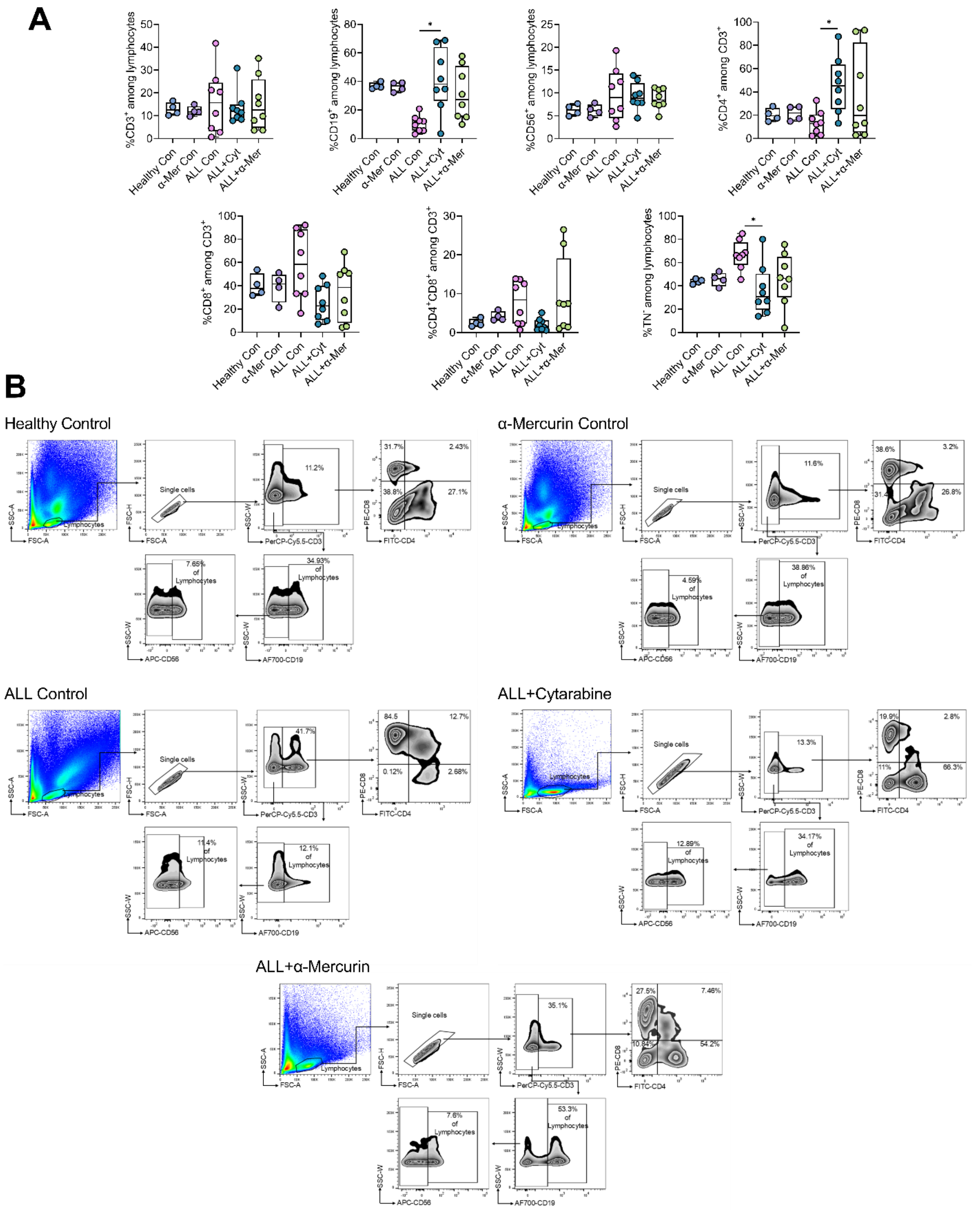

**Figure S8: A.** Graphical representation of percentages of cell populations (%CD3+, %CD19+, CD56+, %CD4+, %CD8+, %CD4+CD8+ and %TN-) investigated by flow cytometric analysis of lymphoid compartments from terminally collected bone marrow aspirates. Data are presented as median with IQR (25 - 75 percentile) in box plots; whiskers represent minimum-maximum values. Kruskal-Wallis test followed by Dunn's multiple comparisons test was performed for statistical analysis. Statistical significance is denoted as *p ≤ 0.05, **p ≤ 0.01, ***p ≤ 0.001, ****p ≤ 0.0001. **B.** Representative flow cytograms across all groups demonstrating %CD3+, %CD3+CD4+, %CD3+CD8+, %CD3+CD4+CD8+ and %CD3+CD4-CD8-, %CD19+ and %CD56+ cells of terminally collected bone marrow aspirates.

#### Extended data of terminal immunophenotypic analysis of thymus

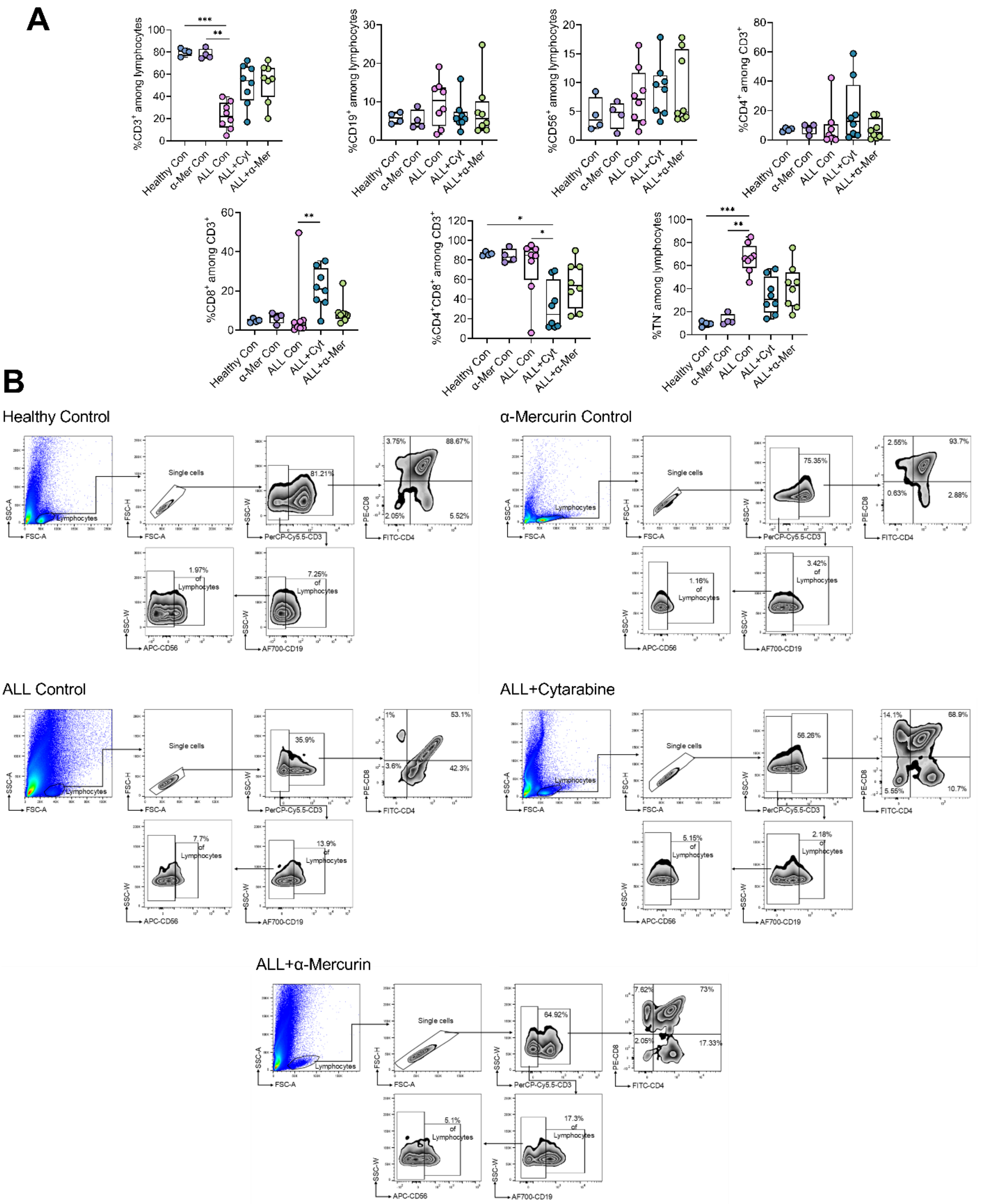

**Figure S9: A.** Graphical representation of percentages of cell populations (%CD3+, %CD19+, CD56+, %CD4+, %CD8+, %CD4+CD8+ and %TN-) investigated by flow cytometric analysis of lymphoid compartments from terminally collected single cells of thymus. Data are presented as median with IQR (25 - 75 percentile) in box plots; whiskers represent minimum-maximum values. Kruskal-Wallis test followed by Dunn's multiple comparisons test was performed for statistical analysis. Statistical significance is denoted as $*p \leq 0.05$, $**p \leq 0.01$, $***p \leq 0.001$, $****p \leq 0.0001$. **B.** Representative flow cytograms across all groups demonstrating %CD3+, %CD3+CD4+, %CD3+CD8+, %CD3+CD4+CD8+ and %CD3+CD4-CD8-, %CD19+ and %CD56+ cells of terminally collected thymocytes.

### Extended data of terminal immunophenotypic analysis of lymph nodes

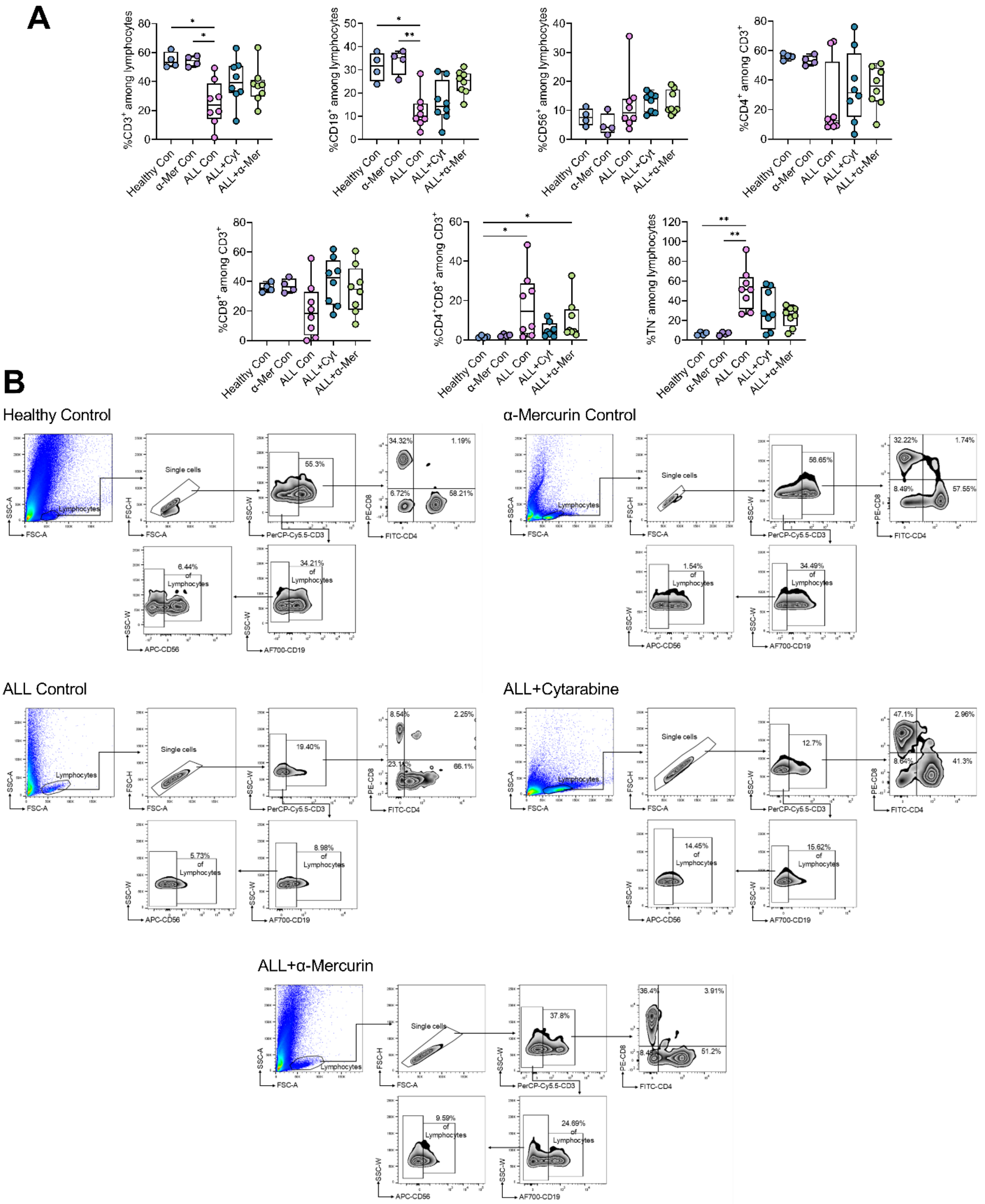

**Figure S10: A.** Graphical representation of percentages of cell populations ($\%CD3^+$, $\%CD19^+$, $CD56^+$, $\%CD4^+$, $\%CD8^+$, $\%CD4^+CD8^+$ and $\%TN^-$) investigated by flow cytometric analysis of lymphoid compartments from terminally collected single cells of lymph nodes. Data are presented as median with IQR (25 - 75 percentile) in box plots; whiskers represent minimum-maximum values. Kruskal-Wallis test followed by Dunn's multiple comparisons test was performed for statistical analysis. Statistical significance is denoted as $*p \le 0.05$, $**p \le 0.01$, $***p \le 0.001$, $****p \le 0.0001$. **B.** Representative flow cytograms across all groups demonstrating $\%CD3^+$, $\%CD3^+CD4^+$, $\%CD3^+CD8^+$, $\%CD3^+CD4^+CD8^+$ and $\%CD3^+CD4^-CD8^-$, $\%CD19^+$ and $\%CD56^+$ cells of terminally collected lymph nodes.

Extended data of terminal immunophenotypic analysis of spleen

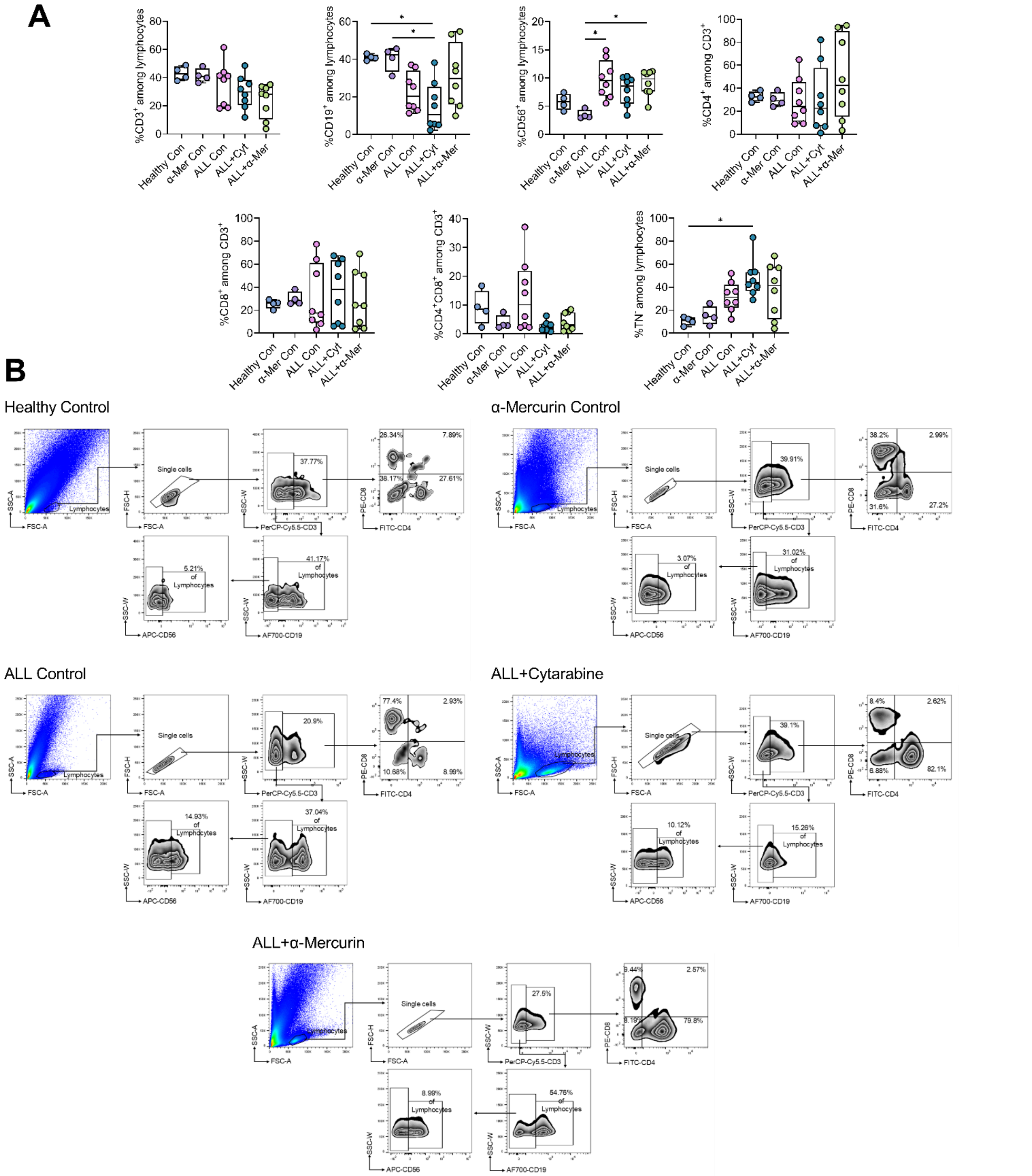

**Figure S11: A.** Graphical representation of percentages of cell populations (%CD3$^+$, %CD19$^+$, CD56$^+$, %CD4$^+$, %CD8$^+$, %CD4$^+$CD8$^+$ and %TN$^-$) investigated by flow cytometric analysis of lymphoid compartments from terminally collected single cells from spleen. Data are presented as median with IQR (25 - 75 percentile) in box plots; whiskers represent minimum-maximum values. Kruskal-Wallis test followed by Dunn's multiple comparisons test was performed for statistical analysis. Statistical significance is denoted as $^*p \leq 0.05$, $^{**}p \leq 0.01$, $^{***}p \leq 0.001$, $^{****}p \leq 0.0001$. **B.** Representative flow cytograms across all groups demonstrating %CD3$^+$, %CD3$^+$CD4$^+$, %CD3$^+$CD8$^+$, %CD3$^+$CD4$^+$CD8$^+$ and %CD3$^+$CD4$^-$CD8$^-$, %CD19$^+$ and %CD56$^+$ cells of terminally collected splenocytes.

#### Median %cell populations with immune index from immunophenotypic analysis

| | | | | | | | | | | | |
|---|---|---|---|---|---|---|---|---|---|---|---|
| **Peripheral Blood** | | | | | | | | | | | |
| | $CD3^+$ | $CD19^+$ | $CD56^+$ | $TN^-$ | $CD4^+$ | $CD8^+$ | $CD4^+CD8^+$ | $CD4^-CD8^-$ | $CD4^+$:$CD8^+$ | T:B | Immune index |
| Healthy Con | 61.35 | 19.29 | 10.91 | 7.94 | 50.50 | 40.80 | 1.88 | 5.90 | 1.24 | 3.19 | 11.63 |
| α-Mer Con | 60.80 | 19.15 | 10.76 | 10.35 | 50.95 | 37.10 | 3.96 | 7.26 | 1.39 | 3.18 | 8.70 |
| ALL Con | 16.10 | 4.18 | 4.08 | 79.08 | 7.27 | 66.35 | 16.16 | 4.53 | 0.10 | 5.13 | 0.31 |
| ALL+Cyt | 39.25 | 16.74 | 13.93 | 31.54 | 39.06 | 47.10 | 7.64 | 3.18 | 0.80 | 2.25 | 2.21 |
| ALL+α-Mer | 33.70 | 11.52 | 9.06 | 39.54 | 37.70 | 43.00 | 4.43 | 5.64 | 0.94 | 2.64 | 1.53 |
| **Bone Marrow Aspirate** | | | | | | | | | | | |
| | $CD3^+$ | $CD19^+$ | $CD56^+$ | $TN^-$ | $CD4^+$ | $CD8^+$ | $CD4^+CD8^+$ | $CD4^-CD8^-$ | $CD4^+$:$CD8^+$ | T:B | Immune index |
| Healthy Con | 12.50 | 36.23 | 6.28 | 44.04 | 19.45 | 37.81 | 2.20 | 36.14 | 0.51 | 0.35 | 1.27 |
| α-Mer Con | 11.70 | 36.79 | 6.10 | 46.00 | 21.96 | 41.60 | 3.76 | 29.85 | 0.65 | 0.32 | 1.18 |
| ALL Con | 15.70 | 10.76 | 9.00 | 66.32 | 12.85 | 58.58 | 8.38 | 17.58 | 0.32 | 1.95 | 0.52 |
| ALL+Cyt | 11.85 | 38.19 | 8.82 | 30.52 | 45.37 | 22.72 | 1.55 | 24.65 | 1.72 | 0.33 | 2.33 |
| ALL+α-Mer | 12.53 | 27.39 | 8.47 | 46.42 | 19.70 | 38.60 | 7.36 | 18.30 | 0.65 | 0.50 | 0.86 |
| **Thymus** | | | | | | | | | | | |
| | $CD3^+$ | $CD19^+$ | $CD56^+$ | $TN^-$ | $CD4^+$ | $CD8^+$ | $CD4^+CD8^+$ | $CD4^-CD8^-$ | $CD4^+$:$CD8^+$ | T:B | Immune index |
| Healthy Con | 80.70 | 5.82 | 3.52 | 9.77 | 7.10 | 4.66 | 85.78 | 1.88 | 1.38 | 14.37 | 9.22 |
| α-Mer Con | 76.60 | 4.39 | 4.82 | 11.26 | 8.48 | 6.81 | 82.73 | 1.98 | 1.15 | 18.47 | 7.81 |
| ALL Con | 21.95 | 10.32 | 7.09 | 59.99 | 2.31 | 2.70 | 84.80 | 7.38 | 0.35 | 2.78 | 0.66 |
| ALL+Cyt | 54.23 | 5.58 | 9.23 | 34.81 | 12.82 | 21.00 | 24.45 | 13.99 | 0.47 | 9.71 | 1.89 |
| ALL+α-Mer | 56.35 | 5.64 | 4.88 | 24.99 | 5.01 | 7.92 | 53.88 | 27.29 | 0.36 | 9.99 | 3.06 |
| **Lymph Nodes** | | | | | | | | | | | |
| | $CD3^+$ | $CD19^+$ | $CD56^+$ | $TN^-$ | $CD4^+$ | $CD8^+$ | $CD4^+CD8^+$ | $CD4^-CD8^-$ | $CD4^+$:$CD8^+$ | T:B | Immune index |
| Healthy Con | 53.33 | 31.68 | 7.53 | 6.39 | 56.14 | 35.40 | 1.37 | 7.84 | 1.57 | 1.69 | 15.12 |
| α-Mer Con | 54.52 | 35.14 | 4.11 | 6.82 | 52.99 | 36.38 | 2.25 | 6.97 | 1.46 | 1.55 | 13.81 |
| ALL Con | 23.85 | 9.93 | 9.13 | 51.33 | 12.40 | 18.35 | 14.59 | 40.08 | 0.42 | 2.12 | 0.92 |
| ALL+Cyt | 39.36 | 14.23 | 13.92 | 26.53 | 31.38 | 42.41 | 3.77 | 10.16 | 0.80 | 2.77 | 2.77 |
| ALL+α-Mer | 37.40 | 25.45 | 10.45 | 26.23 | 36.01 | 34.70 | 4.58 | 21.38 | 1.30 | 1.47 | 2.64 |
| **Spleen** | | | | | | | | | | | |
| | $CD3^+$ | $CD19^+$ | $CD56^+$ | $TN^-$ | $CD4^+$ | $CD8^+$ | $CD4^+CD8^+$ | $CD4^-CD8^-$ | $CD4^+$:$CD8^+$ | T:B | Immune index |
| Healthy Con | 42.78 | 40.91 | 5.80 | 11.05 | 32.57 | 26.07 | 8.63 | 32.73 | 1.31 | 1.05 | 8.31 |
| α-Mer Con | 40.65 | 42.39 | 3.17 | 14.52 | 29.14 | 27.76 | 2.90 | 32.88 | 0.99 | 1.05 | 5.95 |
| ALL Con | 39.85 | 20.34 | 9.52 | 31.22 | 24.10 | 17.50 | 10.09 | 12.93 | 1.28 | 1.42 | 2.28 |
| ALL+Cyt | 29.86 | 10.49 | 8.59 | 43.85 | 22.66 | 38.09 | 2.04 | 15.39 | 0.53 | 3.99 | 1.28 |
| ALL+α-Mer | 28.00 | 29.73 | 9.85 | 41.29 | 42.42 | 24.06 | 3.17 | 20.26 | 1.77 | 3.00 | 1.47 |

**Table S3:** Calculated median values of %immune cell populations along with immune cell ratios and immune index as found from flow cytometric analysis of terminally collected peripheral blood and single cells from lymphoid and myeloid organs (Bone marrow, thymus, lymph nodes and spleen).

#### Extended Data of Brain Histopathological Assessment

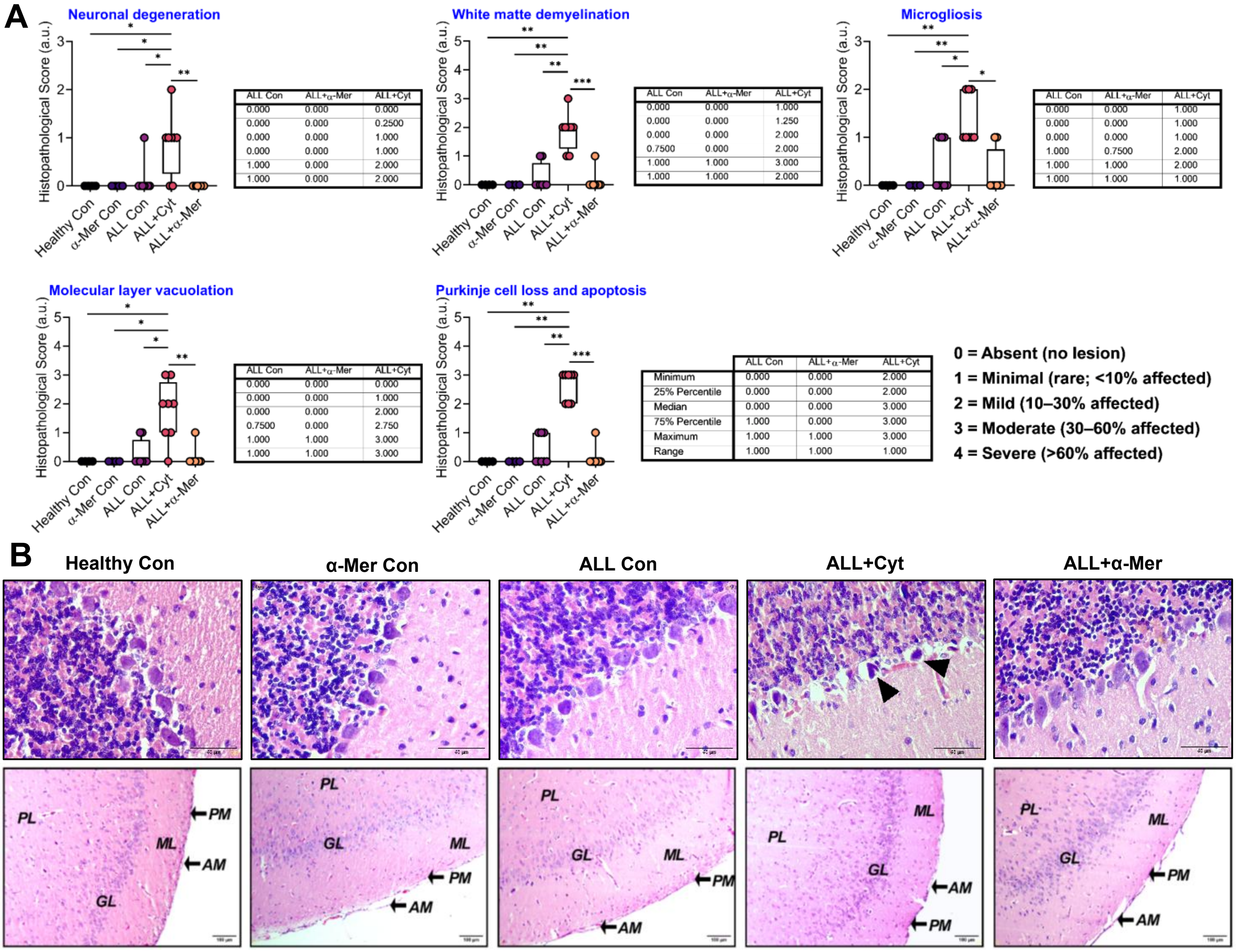

**Figure S12: A.** Graphical representation of individual H&E histopathological score of brain sections across all groups. Score 0 suggests no lesion, 1: <10% affected, 2: 10-30% affected, 3: 30-60% affected and 4: >60% or severely affected. Data are presented as median with IQR (25 - 75 percentile) in box plots; whiskers represent minimum-maximum values. Kruskal-Wallis test followed by Dunn's multiple comparisons test was performed for statistical analysis. Statistical significance is denoted as *$p \leq 0.05$, **$p \leq 0.01$, ***$p \leq 0.001$, ****$p \leq 0.0001$. **B.** Extended representation of H&E-stained sections of cerebellum and cerebrum across all groups. Cytarabine treatment showing significant Purkinje cell injury, and white matter demyelination suggesting treatment-induced neurotoxicity. In contrast, α-Mercurin treated ALL rats maintains intact neuronal morphology and cerebellar architecture with no detectable neurotoxic lesions. Scale bar: 50, 100 µm. Black arrowhead: Purkinje cell injury, ML: Molecular Layer, GL: Granular Layer, PL: Pyramidal Layer, AM: Arachnoid Mater, PM: Pia Mater.

#### Extended Data of Cardiac Histopathological Assessment

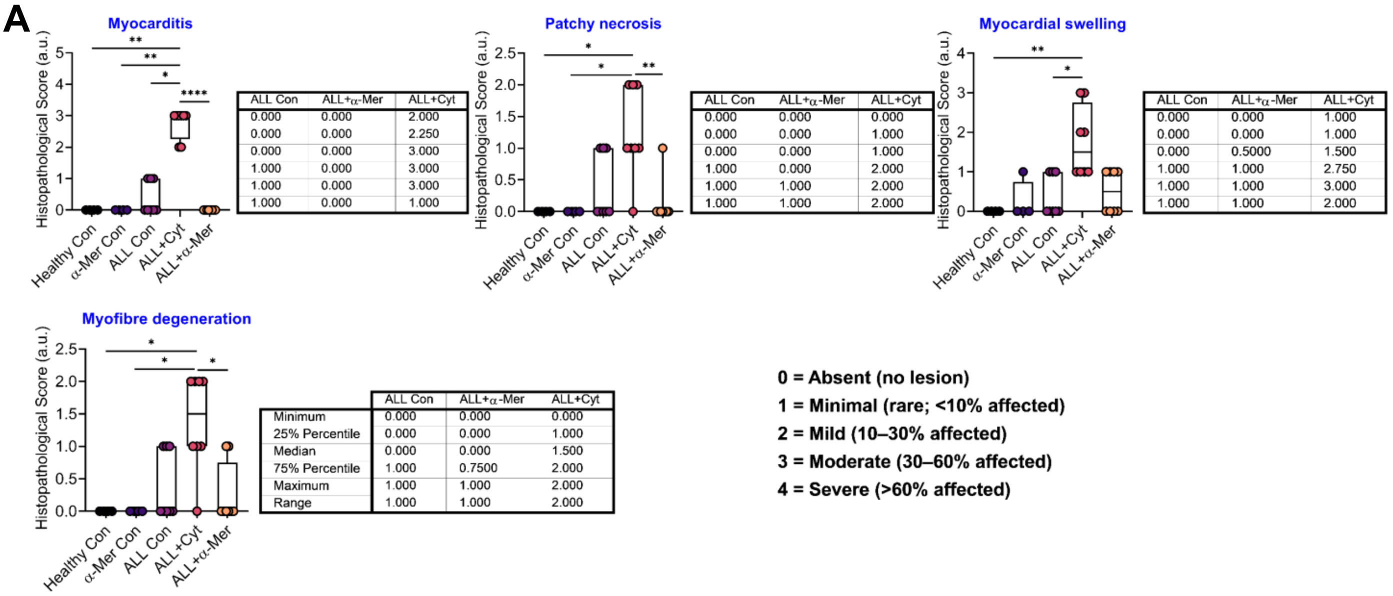

Myocarditis

| ALL Con | ALL+α-Mer | ALL+Cyt |
|---|---|---|
| 0.000 | 0.000 | 2.000 |
| 0.000 | 0.000 | 2.250 |
| 0.000 | 0.000 | 3.000 |
| 1.000 | 0.000 | 3.000 |
| 1.000 | 0.000 | 3.000 |
| 1.000 | 0.000 | 1.000 |

Patchy necrosis

| ALL Con | ALL+α-Mer | ALL+Cyt |
|---|---|---|
| 0.000 | 0.000 | 0.000 |
| 0.000 | 0.000 | 1.000 |
| 0.000 | 0.000 | 1.000 |
| 1.000 | 0.000 | 2.000 |
| 1.000 | 1.000 | 2.000 |
| 1.000 | 1.000 | 2.000 |

Myocardial swelling

| ALL Con | ALL+α-Mer | ALL+Cyt |
|---|---|---|
| 0.000 | 0.000 | 1.000 |
| 0.000 | 0.000 | 1.000 |
| 0.000 | 0.5000 | 1.500 |
| 1.000 | 1.000 | 2.750 |
| 1.000 | 1.000 | 3.000 |
| 1.000 | 1.000 | 2.000 |

Myofibre degeneration

| | ALL Con | ALL+α-Mer | ALL+Cyt |
|---|---|---|---|
| Minimum | 0.000 | 0.000 | 0.000 |
| 25% Percentile | 0.000 | 0.000 | 1.000 |
| Median | 0.000 | 0.000 | 1.500 |
| 75% Percentile | 1.000 | 0.7500 | 2.000 |
| Maximum | 1.000 | 1.000 | 2.000 |
| Range | 1.000 | 1.000 | 2.000 |

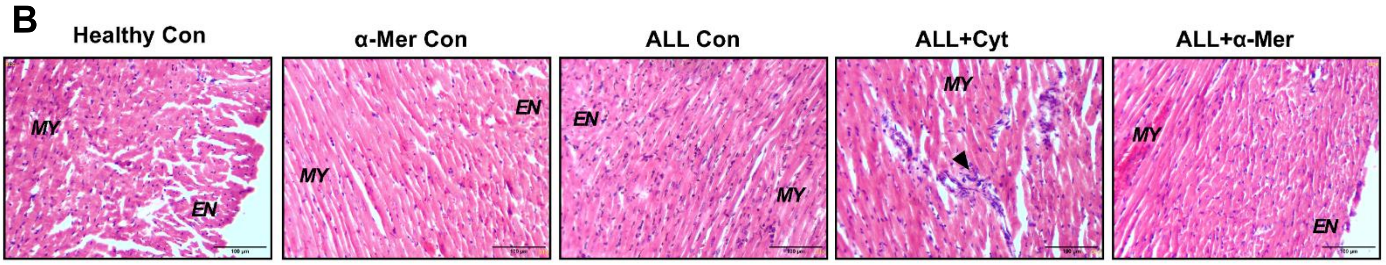

**Figure S13: A.** Graphical representation of individual H&E histopathological score of cardiac sections across all groups. Score 0 suggests no lesion, 1: <10% affected, 2: 10-30% affected, 3: 30-60% affected and 4: >60% or severely affected. Data are presented as median with IQR (25 - 75 percentile) in box plots; whiskers represent minimum-maximum values. Kruskal-Wallis test followed by Dunn's multiple comparisons test was performed for statistical analysis. Statistical significance is denoted as *p ≤ 0.05, **p ≤ 0.01, ***p ≤ 0.001, ****p ≤ 0.0001. **B.** Extended representation of H&E-stained cardiac sections across all groups. Cytarabine treatment exhibits pronounced myocardial injury as shown by myocarditis. In contrast, α-Mercurin-treated ALL depicts largely preserved myocardial structure. Scale bar: 100 µm. MY: Myocardium, EN: Endocardium, Black arrowhead: Myocarditis.

#### Extended Data of Liver Histopathological Assessment

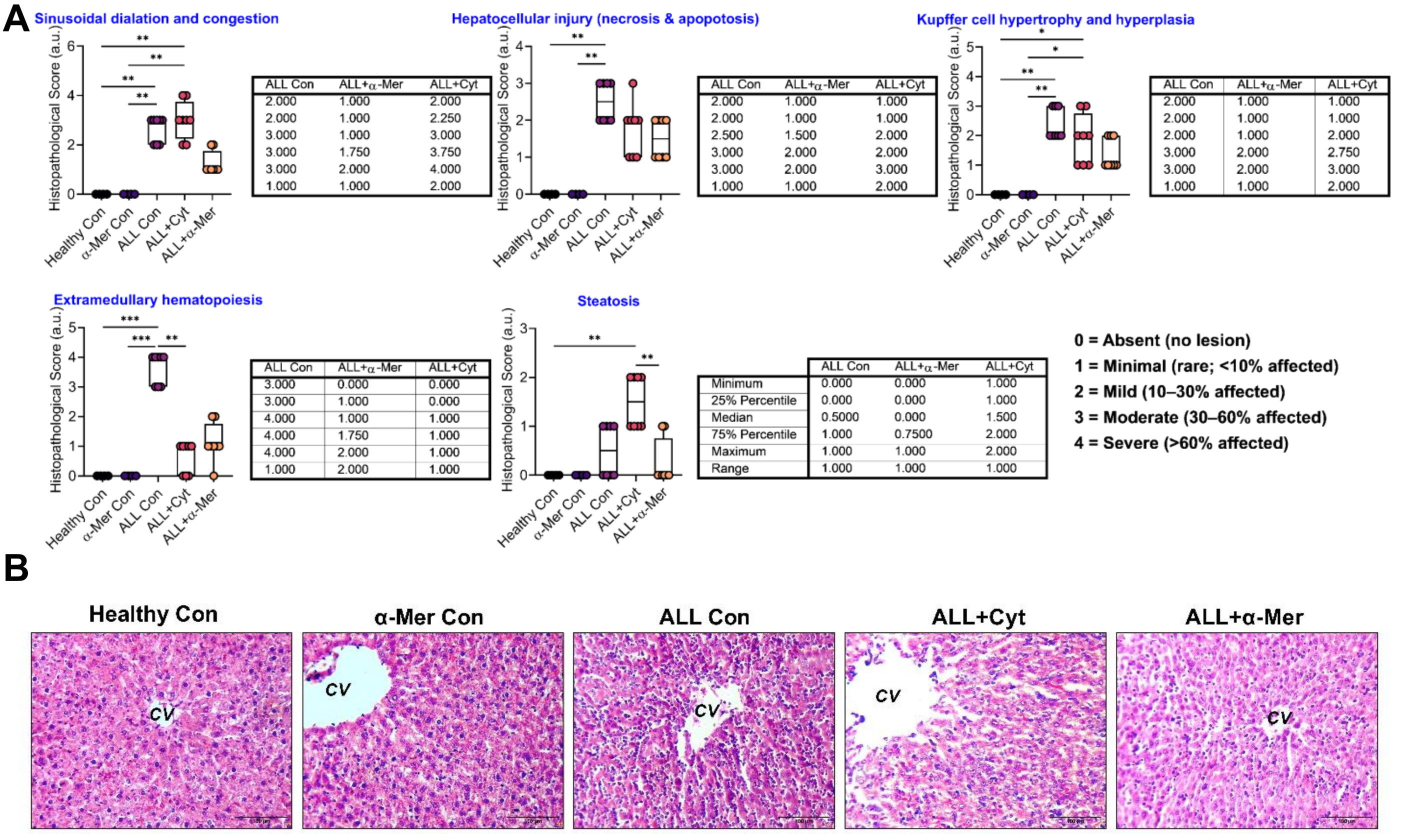

**Figure S14: A.** Graphical representation of individual H&E histopathological score of liver sections across all groups. Score 0 suggests no lesion, 1: <10% affected, 2: 10-30% affected, 3: 30-60% affected and 4: >60% or severely affected. Data are presented as median with IQR (25 - 75 percentile) in box plots; whiskers represent minimum-maximum values. Kruskal-Wallis test followed by Dunn's multiple comparisons test was performed for statistical analysis. Statistical significance is denoted as *p ≤ 0.05, **p ≤ 0.01, ***p ≤ 0.001, ****p ≤ 0.0001. **B.** Extended representation of H&E-stained sections of liver across all groups. Treatment with cytarabine reduces the disease burden, simultaneously it also exerts marked hepatocellular toxicity. However, no intrinsic hepatotoxicity is detected following α-Mercurin treatment. Scale bar: 100 μm. CV: central vein

#### Extended Data of Kidney Histopathological Assessment

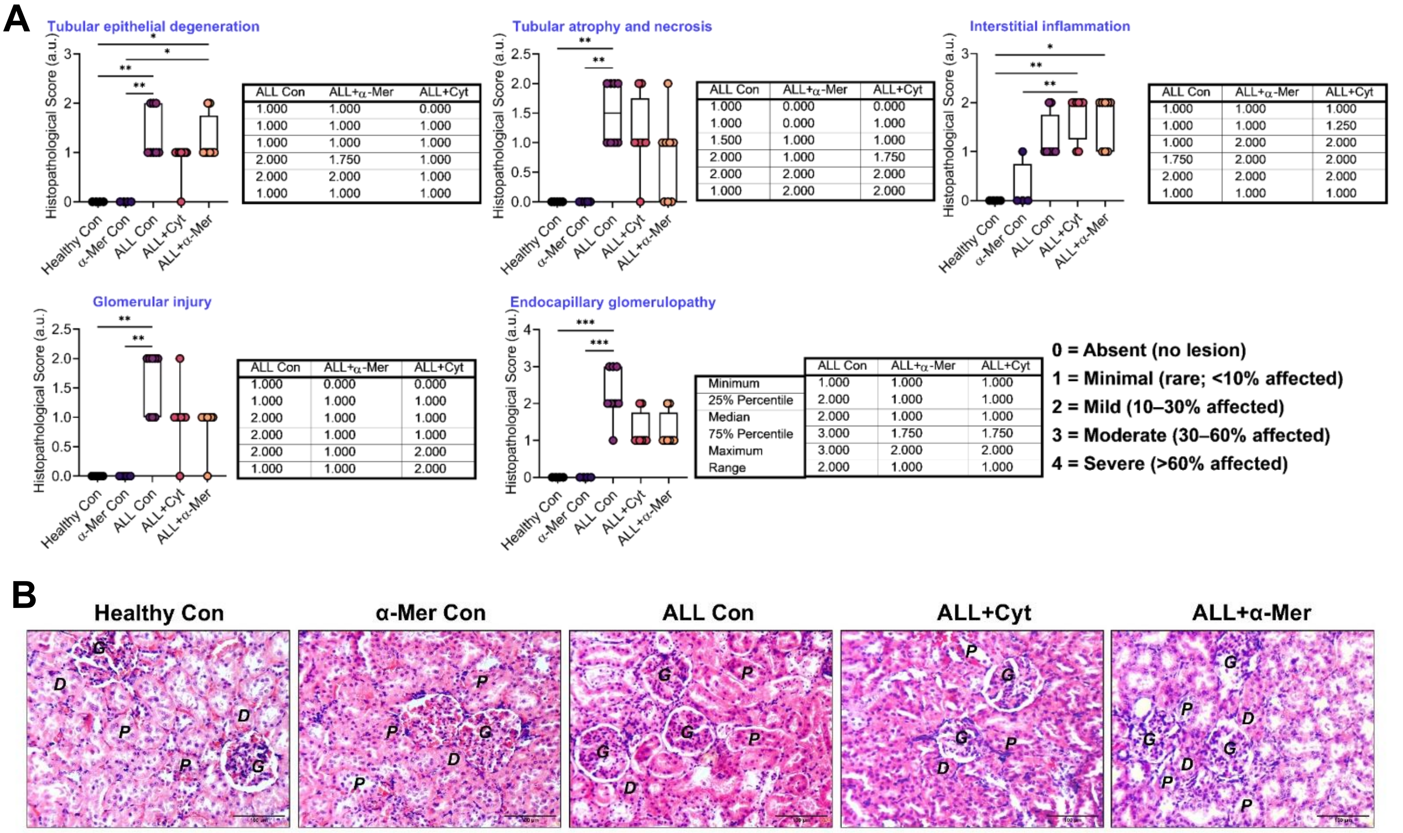

**Figure S15: A.** Graphical representation of individual H&E histopathological score of kidney sections across all groups. Score 0 suggests no lesion, 1: <10% affected, 2: 10-30% affected, 3: 30-60% affected and 4: >60% or severely affected. Data are presented as median with IQR (25 - 75 percentile) in box plots; whiskers represent minimum-maximum values. Kruskal-Wallis test followed by Dunn's multiple comparisons test was performed for statistical analysis. Statistical significance is denoted as *p ≤ 0.05, **p ≤ 0.01, ***p ≤ 0.001, ****p ≤ 0.0001. **B.** Extended representation of H&E-stained sections of kidney across all groups. Both cytarabine and α-Mercurin treatments suggest reduced leukemic burden and associated inflammatory changes, showing no evidence of intrinsic nephrotoxicity. Scale bar: 100 µm. G: glomerulus, P: proximal tubule, D: distal tubule.

#### Extended Data of Lung Histopathological Assessment

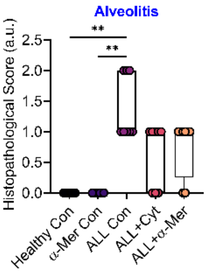

| ALL Con | ALL+α-Mer | ALL+Cyt |
|---|---|---|
| 1.000 | 0.000 | 0.000 |
| 1.000 | 0.2500 | 0.000 |
| 1.000 | 1.000 | 1.000 |
| 2.000 | 1.000 | 1.000 |
| 2.000 | 1.000 | 1.000 |
| 1.000 | 1.000 | 1.000 |

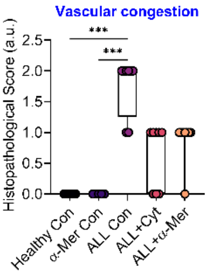

| ALL Con | ALL+α-Mer | ALL+Cyt |
|---|---|---|
| 1.000 | 0.000 | 0.000 |
| 1.250 | 1.000 | 0.000 |
| 2.000 | 1.000 | 1.000 |
| 2.000 | 1.000 | 1.000 |
| 2.000 | 1.000 | 1.000 |
| 1.000 | 1.000 | 1.000 |

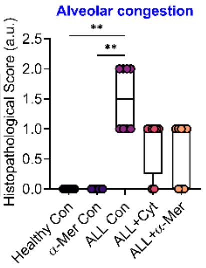

| ALL Con | ALL+α-Mer | ALL+Cyt |
|---|---|---|
| 1.000 | 0.000 | 0.000 |
| 1.000 | 0.000 | 0.2500 |
| 1.500 | 1.000 | 1.000 |
| 2.000 | 1.000 | 1.000 |
| 2.000 | 1.000 | 1.000 |
| 1.000 | 1.000 | 1.000 |

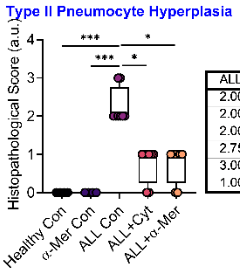

| ALL Con | ALL+α-Mer | ALL+Cyt |
|---|---|---|
| 2.000 | 0.000 | 0.000 |
| 2.000 | 0.2500 | 0.2500 |
| 2.000 | 1.000 | 1.000 |
| 2.750 | 1.000 | 1.000 |
| 3.000 | 1.000 | 1.000 |
| 1.000 | 1.000 | 1.000 |

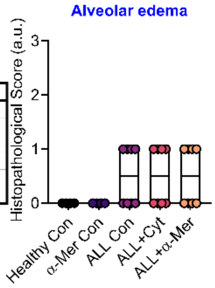

| | ALL Con | ALL+α-Mer | ALL+Cyt |
|---|---|---|---|
| Minimum | 0.000 | 0.000 | 0.000 |
| 25% Percentile | 0.000 | 0.000 | 0.000 |
| Median | 0.5000 | 0.5000 | 0.5000 |
| 75% Percentile | 1.000 | 1.000 | 1.000 |
| Maximum | 1.000 | 1.000 | 1.000 |
| Range | 1.000 | 1.000 | 1.000 |

**0 = Absent (no lesion)**
**1 = Minimal (rare; <10% affected)**
**2 = Mild (10–30% affected)**
**3 = Moderate (30–60% affected)**
**4 = Severe (>60% affected)**

**Figure S16:** Graphical representation of individual H&E histopathological score of lung sections across all groups. Score 0 suggests no lesion, 1: <10% affected, 2: 10-30% affected, 3: 30-60% affected and 4: >60% or severely affected. Data are presented as median with IQR (25 - 75 percentile) in box plots; whiskers represent minimum-maximum values. Kruskal-Wallis test followed by Dunn's multiple comparisons test was performed for statistical analysis. Statistical significance is denoted as $*p \leq 0.05$, $**p \leq 0.01$, $***p \leq 0.001$, $****p \leq 0.0001$.

#### Extended Data of Spleen Histopathological Assessment

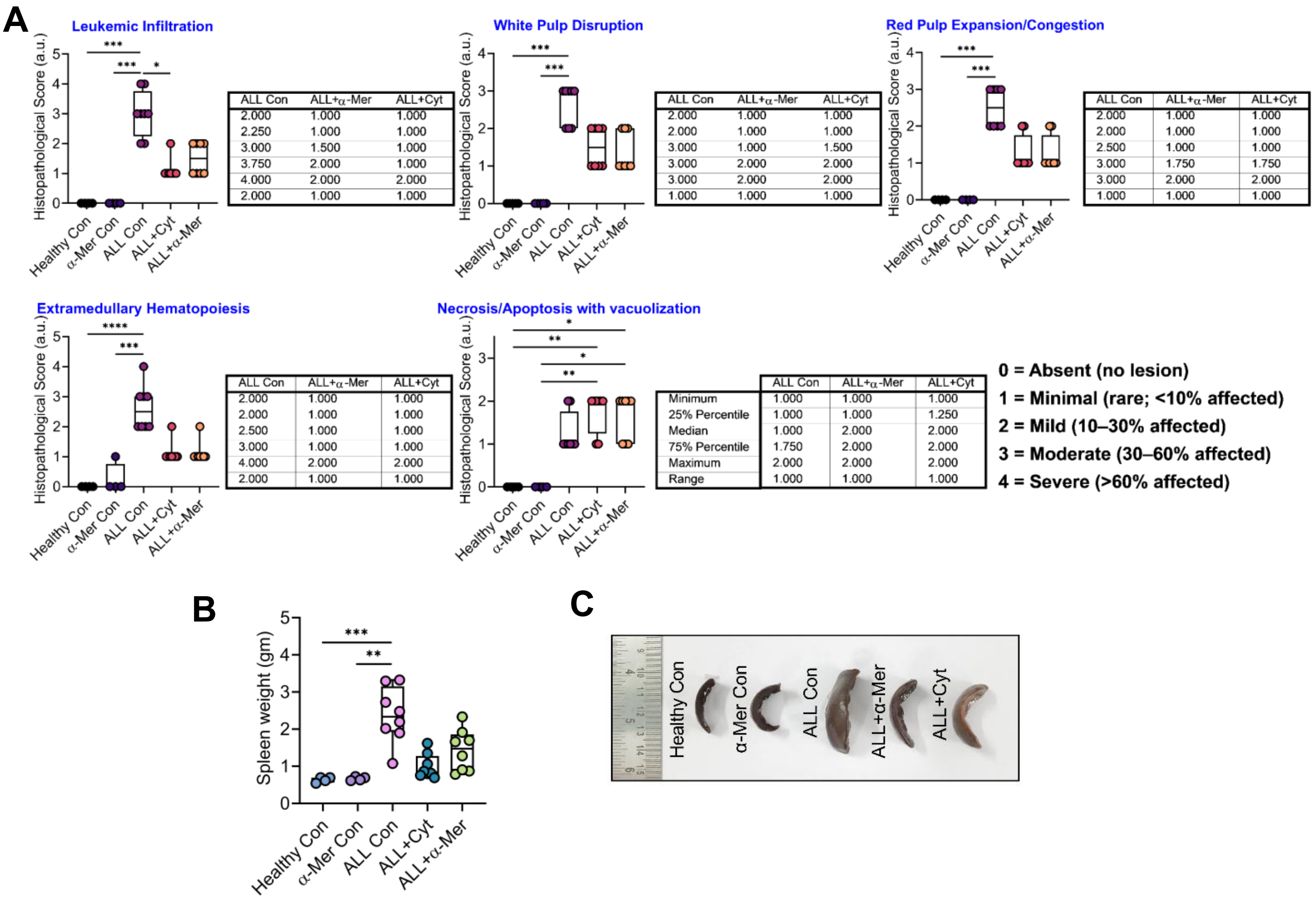

**Figure S17: A.** Graphical representation of individual H&E histopathological score of spleen sections across all groups. Score 0 suggests no lesion, 1: <10% affected, 2: 10-30% affected, 3: 30-60% affected and 4: >60% or severely affected. Data are presented as median with IQR (25 - 75 percentile) in box plots; whiskers represent minimum-maximum values. Kruskal-Wallis test followed by Dunn's multiple comparisons test was performed for statistical analysis. Statistical significance is denoted as *p ≤ 0.05, **p ≤ 0.01, ***p ≤ 0.001, ****p ≤ 0.0001. **B.** Spleen weight across experimental groups, suggesting marked splenomegaly in ALL controls, that is partially reduced by α-Mercurin treatment, similar to cytarabine. **C.** Representative comparison of spleen size, showing marked splenomegaly in ALL controls, which is reduced toward normal levels following treatment.

#### Extended Data of Lymph Node Histopathological Assessment

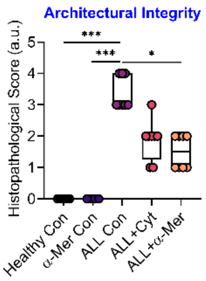

| ALL Con | ALL+α-Mer | ALL+Cyt |
|---|---|---|
| 3.000 | 1.000 | 1.000 |
| 3.000 | 1.000 | 1.250 |
| 3.000 | 1.500 | 2.000 |
| 4.000 | 2.000 | 2.000 |
| 4.000 | 2.000 | 3.000 |
| 1.000 | 1.000 | 2.000 |

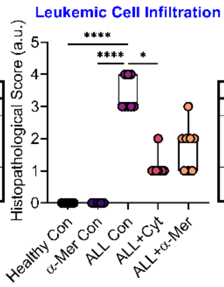

| ALL Con | ALL+α-Mer | ALL+Cyt |
|---|---|---|
| 3.000 | 1.000 | 1.000 |
| 3.000 | 1.000 | 1.000 |
| 3.000 | 2.000 | 1.000 |
| 4.000 | 2.000 | 1.000 |
| 4.000 | 3.000 | 2.000 |
| 1.000 | 2.000 | 1.000 |

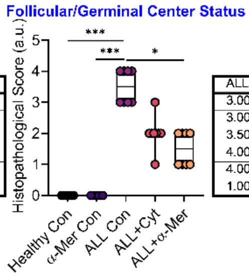

| ALL Con | ALL+α-Mer | ALL+Cyt |
|---|---|---|
| 3.000 | 1.000 | 1.000 |
| 3.000 | 1.000 | 2.000 |
| 3.500 | 1.500 | 2.000 |
| 4.000 | 2.000 | 2.000 |
| 4.000 | 2.000 | 3.000 |
| 1.000 | 1.000 | 2.000 |

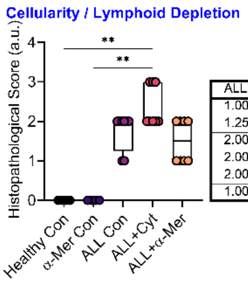

| ALL Con | ALL+α-Mer | ALL+Cyt |
|---|---|---|
| 1.000 | 1.000 | 2.000 |
| 1.250 | 1.000 | 2.000 |
| 2.000 | 1.500 | 2.000 |
| 2.000 | 2.000 | 3.000 |
| 2.000 | 2.000 | 3.000 |
| 1.000 | 1.000 | 1.000 |

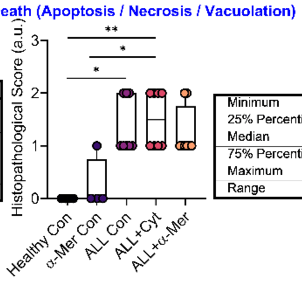

| | ALL Con | ALL+α-Mer | ALL+Cyt |
|---|---|---|---|
| Minimum | 1.000 | 1.000 | 1.000 |
| 25% Percentile | 1.000 | 1.000 | 1.000 |
| Median | 1.000 | 1.000 | 1.500 |
| 75% Percentile | 2.000 | 1.750 | 2.000 |
| Maximum | 2.000 | 2.000 | 2.000 |
| Range | 1.000 | 1.000 | 1.000 |

**0 = Absent (no lesion)**
**1 = Minimal (rare; <10% affected)**
**2 = Mild (10–30% affected)**
**3 = Moderate (30–60% affected)**
**4 = Severe (>60% affected)**

**Figure S18:** Graphical representation of individual H&E histopathological score of lymph node sections across all groups. Score 0 suggests no lesion, 1: <10% affected, 2: 10-30% affected, 3: 30-60% affected and 4: >60% or severely affected. Data are presented as median with IQR (25 - 75 percentile) in box plots; whiskers represent minimum-maximum values. Kruskal-Wallis test followed by Dunn's multiple comparisons test was performed for statistical analysis. Statistical significance is denoted as *p ≤ 0.05, **p ≤ 0.01, ***p ≤ 0.001, ****p ≤ 0.0001.

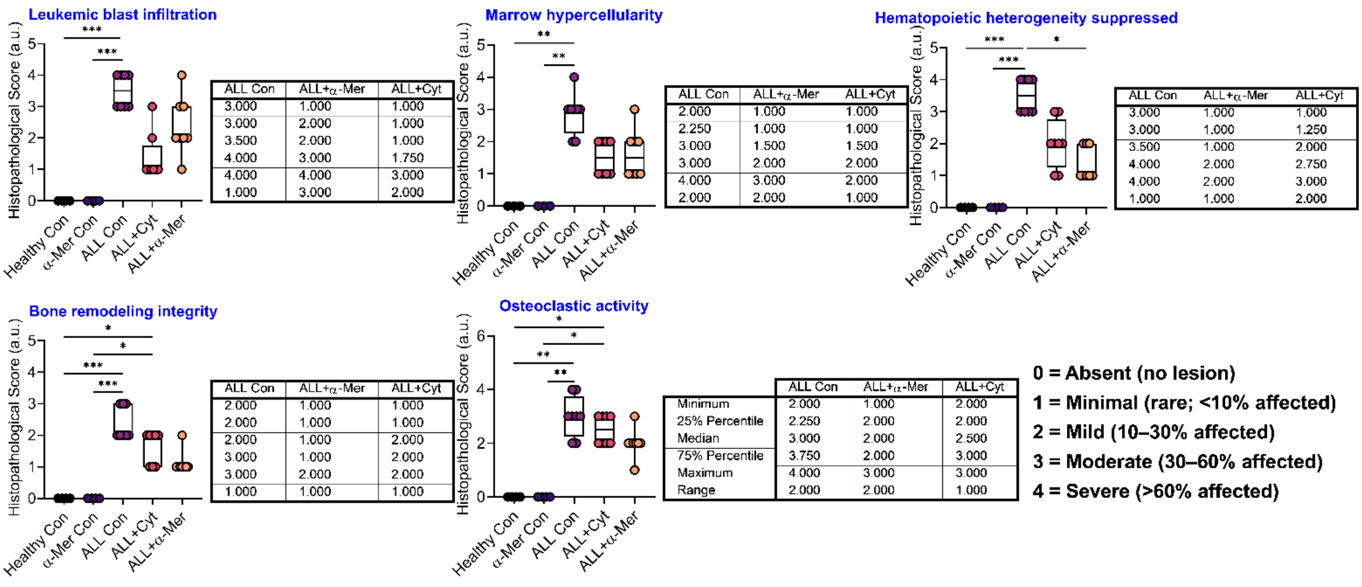

**Leukemic blast infiltration**

| ALL Con | ALL+α-Mer | ALL+Cyt |
|---|---|---|
| 3.000 | 1.000 | 1.000 |
| 3.000 | 2.000 | 1.000 |
| 3.500 | 2.000 | 1.000 |
| 4.000 | 3.000 | 1.750 |
| 4.000 | 4.000 | 3.000 |
| 1.000 | 3.000 | 2.000 |

**Marrow hypercellularity**

| ALL Con | ALL+α-Mer | ALL+Cyt |
|---|---|---|
| 2.000 | 1.000 | 1.000 |
| 2.250 | 1.000 | 1.000 |
| 3.000 | 1.500 | 1.500 |
| 3.000 | 2.000 | 2.000 |
| 4.000 | 3.000 | 2.000 |
| 2.000 | 2.000 | 1.000 |

**Hematopoietic heterogeneity suppressed**

| ALL Con | ALL+α-Mer | ALL+Cyt |
|---|---|---|
| 3.000 | 1.000 | 1.000 |
| 3.000 | 1.000 | 1.250 |
| 3.500 | 1.000 | 2.000 |
| 4.000 | 2.000 | 2.750 |
| 4.000 | 2.000 | 3.000 |
| 1.000 | 1.000 | 2.000 |

**Bone remodeling integrity**

| ALL Con | ALL+α-Mer | ALL+Cyt |
|---|---|---|
| 2.000 | 1.000 | 1.000 |
| 2.000 | 1.000 | 1.000 |
| 2.000 | 1.000 | 2.000 |
| 3.000 | 1.000 | 2.000 |
| 3.000 | 2.000 | 2.000 |
| 1.000 | 1.000 | 1.000 |

**Osteoclastic activity**

| | ALL Con | ALL+α-Mer | ALL+Cyt |
|---|---|---|---|
| Minimum | 2.000 | 1.000 | 2.000 |
| 25% Percentile | 2.250 | 2.000 | 2.000 |
| Median | 3.000 | 2.000 | 2.500 |
| 75% Percentile | 3.750 | 2.000 | 3.000 |
| Maximum | 4.000 | 3.000 | 3.000 |
| Range | 2.000 | 2.000 | 1.000 |

**Figure S19:** Graphical representation of individual H&E histopathological score of bone along with bone marrow sections across all groups. Score 0 suggests no lesion, 1: <10% affected, 2: 10-30% affected, 3: 30-60% affected and 4: >60% or severely affected. Data are presented as median with IQR (25 - 75 percentile) in box plots; whiskers represent minimum-maximum values. Kruskal-Wallis test followed by Dunn's multiple comparisons test was performed for statistical analysis. Statistical significance is denoted as *p ≤ 0.05, **p ≤ 0.01, ***p ≤ 0.001, ****p ≤ 0.0001.

#### Extended Data of Thymus Histopathological Assessment

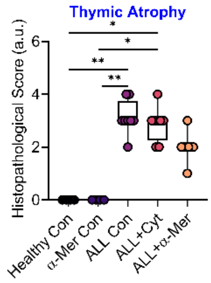

| ALL Con | ALL+α-Mer | ALL+Cyt |
|---|---|---|
| 2.000 | 1.000 | 2.000 |
| 3.000 | 2.000 | 2.250 |
| 3.000 | 2.000 | 3.000 |
| 3.750 | 2.000 | 3.000 |
| 4.000 | 3.000 | 4.000 |
| 2.000 | 2.000 | 2.000 |

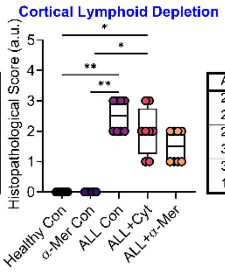

| ALL Con | ALL+α-Mer | ALL+Cyt |
|---|---|---|
| 2.000 | 1.000 | 1.000 |
| 2.000 | 1.000 | 1.250 |
| 2.500 | 1.500 | 2.000 |
| 3.000 | 2.000 | 2.750 |
| 3.000 | 2.000 | 3.000 |
| 1.000 | 1.000 | 2.000 |

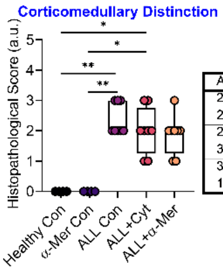

| ALL Con | ALL+α-Mer | ALL+Cyt |
|---|---|---|
| 2.000 | 1.000 | 1.000 |
| 2.000 | 1.250 | 1.250 |
| 2.000 | 2.000 | 2.000 |
| 3.000 | 2.000 | 2.750 |
| 3.000 | 3.000 | 3.000 |
| 1.000 | 2.000 | 2.000 |

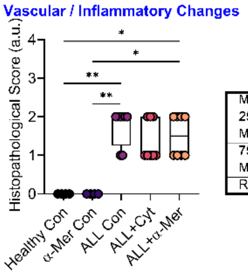

| | ALL Con | ALL+α-Mer | ALL+Cyt |
|---|---|---|---|
| Minimum | 1.000 | 1.000 | 1.000 |
| 25% Percentile | 1.250 | 1.000 | 1.000 |
| Median | 2.000 | 1.500 | 1.000 |
| 75% Percentile | 2.000 | 2.000 | 2.000 |
| Maximum | 2.000 | 2.000 | 2.000 |
| Range | 1.000 | 1.000 | 1.000 |

**0 = Absent (no lesion)**
**1 = Minimal (rare; <10% affected)**
**2 = Mild (10–30% affected)**
**3 = Moderate (30–60% affected)**
**4 = Severe (>60% affected)**

**Figure S20:** Graphical representation of individual H&E histopathological score of thymus sections across all groups. Score 0 suggests no lesion, 1: <10% affected, 2: 10-30% affected, 3: 30-60% affected and 4: >60% or severely affected. Data are presented as median with IQR (25 - 75 percentile) in box plots; whiskers represent minimum-maximum values. Kruskal-Wallis test followed by Dunn's multiple comparisons test was performed for statistical analysis. Statistical significance is denoted as *p ≤ 0.05, **p ≤ 0.01, ***p ≤ 0.001, ****p ≤ 0.0001.
